# Experimental evolution unveils a role for structural variants and sex-specificity in the genetic architecture of lifespan

**DOI:** 10.64898/2026.10.28.762418

**Authors:** Athanasios Toros, Mats E. Pettersson, Angela P. Fuentes-Pardo, Alexandre Rêgo, Carl-Johan Rubin, Ahmed Sayadi, Mirko Đorđević, Uroš Savković, Biljana Stojković, Elina Immonen, Marek Kucka, Yingguang Frank Chan, Leif Andersson, Göran Arnqvist

## Abstract

Insights into the genetic basis of lifespan evolution can link evolutionary theory with an understanding of health, disease and adaptation. We leverage the power of >200 generations of experimental evolution in a seed beetle to unveil the genomic architecture of lifespan evolution. Replicated populations evolved under regimes favoring either a short or long life, resulting in a near doubling of lifespan. Whole genome re-sequencing revealed that short- and long-lived lines were strongly differentiated (global *F*_ST_ = 0.41) and more than 700 genes showed consistent signs of divergent selection. Remarkably, short- and long-lived lines were consistently divergent in a large number of regions harboring copy-number variants (CNVs), involving >600 genes. The evolution of a long life involved many distinct biological processes, such as DNA fidelity, mitotic integrity, reproduction, metabolism, immunity and stress response. A pedigree-based genome scan for quantitative trait loci (QTLs) uncovered several QTLs with large effects on lifespan, but the genetic basis of a long life was dominated by a very large number of regions with medium and small effects. The genome-wide spatial association with QTL effects was, if anything, stronger for CNVs than for sequence variants. The genetic effects of QTLs on lifespan were positively correlated overall between the sexes, but four regions showed pronounced sex-differences. Our results show that structural variants play a significant role in the evolution of an extended lifespan and the highly polygenic genomic architecture is to some extent sex-specific, consistent with differences in physiology and life history tradeoffs between the sexes.

**Significance Statement:** Why some populations evolve long lives while others do not remains a central question in evolutionary biology and biomedicine. Using more than 200 generations of experimental evolution in seed beetles, we show that lifespan can evolve dramatically through changes spread across hundreds of genomic regions rather than a few major genes. A striking finding is that copy-number variants, a form of structural genetic variation often overlooked in studies of aging, contributed strongly to lifespan evolution. Genes involved in DNA maintenance, reproduction, metabolism, immunity, and stress responses were all affected. The genetic basis of lifespan was also partly sex-specific, highlighting that males and females may face different evolutionary and physiological constraints on aging.

## Introduction

Understanding why organisms age is a fundamental and longstanding challenge in biology. While the field of biogerontology strives to gain a detailed physiological understanding using experimental tools and bottom-up methods (1), evolutionary biologists strive to understand ageing by studying the evolution and genetics of lifespan through a top-down approach (2). The former implicitly assumes that a few key pathways regulate lifespan, but the latter instead predicts that lifespan is a polygenic trait with a complex genetic architecture.

Although both approaches have generated significant progress, there are still important gaps in our understanding of ageing (3). For example, while experimental studies show that a substantial fraction of the genome can regulate lifespan (4, 5), we know much less about the genetic basis of actual lifespan evolution (6, 7).

Much of what we know about the genetic architecture of adaptive evolution is based on segregating single-nucleotide polymorphisms (SNPs). Yet, there is a rising recognition that structural variation in the genome, in the form of variation in copy number variants (CNVs; deletions, insertions, duplications), orientation (inversions), and chromosomal location (translocations), represents an important source of genetic variation in the adaptive evolution of life history traits (8, 9). In particular, CNVs are very abundant in most taxa and it has been predicted that CNVs may play an even more important role than sequence variants in adaptation (10, 11). Major inversions have been associated with variation in lifespan in some insects (12–14) and CNVs are known to be associated with ageing and lifespan in humans (15, 16). However, the relative contribution of sequence variants and CNVs to lifespan evolution is currently an open question, in part due to the methodological challenges with detecting CNVs in genome-wide data.

Females live longer than males in most animals and life history theory predicts that optimal female lifespan should often exceed that in males (17, 18). Comparative studies show that sexual dimorphism in lifespan can evolve rapidly (19), yet the fact that males and females share a common genome should constrain the evolution of sexual dimorphism (20–22) and the underlying causes of sex differences in aging are largely unknown (23). An improved understanding of sex differences in lifespan may provide novel insights into both the evolution of and the mechanisms of aging (23–26). Male and female physiologies differ and it has been predicted that sexually antagonistic pleiotropy may contribute to ageing, in particular if alleles have opposing effects on lifespan in the sexes (27). Unfortunately, few studies have characterized sex-specificity in the genetic architecture of lifespan. Quantitative genetic studies have suggested that the genetic basis for the regulation of lifespan is shared between males and females to a large extent (28–31) while studies of flies have revealed a more complex genetic architecture with sex- and context-specificity (6, 27, 32–34).

Experimental evolution (hence, EE) studies of lifespan and ageing can provide direct assessments of the genetic basis of lifespan evolution under controlled conditions, which can help us unmask the ultimate processes that are responsible for the evolution of ageing and a restricted lifespan (35). The replication of EE populations also allows assessment of the repeatability of adaptive evolution of complex traits based on standing genetic variation (36). A number of EE studies have provided crucial insights into the phenotypic correlates of lifespan evolution (37), but only three EE experiments have combined EE with whole genome resequencing (evolve-and-resequence) to characterize the genetic architecture of evolved differences in lifespan (38–40). All three were performed with *Drosophila melanogaster* and each implicated large regions of the genome, harbouring many hundreds of potentially causal genes, as being involved in lifespan evolution. While it is clear that evolve and resequence can provide important insights into the evolution of lifespan, it is equally clear that we need studies of a larger diversity of taxa including non-model organisms (37).

Here, we employ nearly 40 years (>200 generations) of divergent EE in the seed beetle *Acanthoscelides obtectus* and amalgamate a bottom-up and a top-down genomic approach to characterize the genetic basis of lifespan evolution from standing genetic variation and assess its repeatability. Replicated EE lines, derived from a common founding population established in 1983, have been maintained under conditions allowing for reproduction either early (E) or late (L) in life. This has resulted in a striking decelerated senescence and near doubling of lifespan in L lines, which has been accompanied by an increased juvenile developmental time, larger adult body size, higher lipid content, a lower metabolic rate, a metabolic machinery more reliant on fatty acid oxidation, increased resistance to oxidative stress, altered mitochondrial bioenergetics and a shift in reproductive effort towards later ages in L compared with E lines (41–48). Interestingly, the evolution of lifespan under EE has been more rapid in females, resulting in the evolution of increased sexual dimorphism in lifespan in L lines (Fig. 1). Immonen et al. (49) showed that while >1,600 genes were differentially expressed in L and E, the differences in expression were also overall more pronounced in females. Here, we first scaffold a previous genome assembly and then used whole genome population resequencing of our replicated EE lines to assess genetic differentiation between E and L lines and to identify regions of the genome experiencing selective sweeps. We then make a dedicated genome-wide effort to assess the prediction that CNVs play a central role for the evolution of lifespan. To test the prediction that the genetic architecture of lifespan is sex-specific, we conduct a large genotype-phenotype association experiment of lifespan and body size. Finally, we amalgamate our complimentary attempts to better characterize the functional genetic architecture of lifespan evolution.

**Fig. 1.**
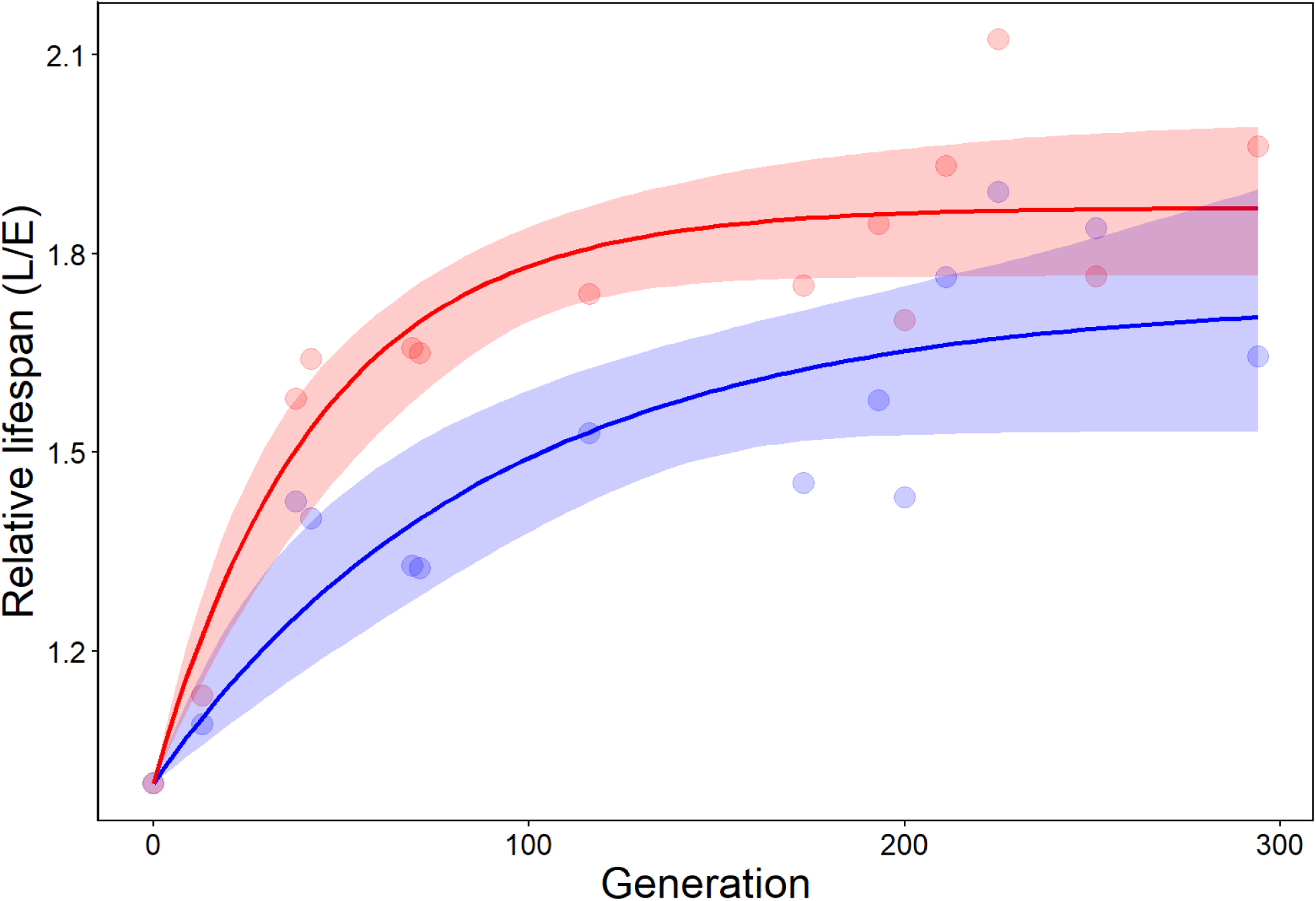
Mean divergence in lifespan between L and E lines in females (red) and males (blue) over the course of experimental evolution. Solid lines represent the fitted function (relative lifespan = 1 + *a*[1 − *b ^generation^*]) and shaded areas the 95% bootstrap CI. Lifespan in females responded more rapidly and to a greater degree than in males, and sexual dimorphism in lifespan was larger in L lines than in E lines (SDI [{female lifespan / male lifespan }-1] in L = 0.27 and in E = 0.11; paired *t*-test, *t*_12_ = 6.89, P < 0.001). Data from (49).

## Results

### Genetic differentiation

Our Hi-C genome assembly contained some 90% of all scaffolds within 10 chromosome-length scaffolds (SI; Fig. S1). We combined DNA of 50 males and 50 females in sex-specific pools for each of the 8 replicate lines (2 EE treatments × 4 replicate lines per treatment × 2 sex-specific pools per replicate line = 16 pools) and performed whole-genome sequencing to ∼100X coverage per pool. We recovered in total 7,977,586 well supported SNPs in the resequencing data. Overall, the global genomic differentiation between E and L lines was as high as *F*_ST_ = 0.41 and all pair-wise comparisons between E and L lines were *F*_ST_ ≍ 0.4 (Fig. 2). The two sex-specific pools from the same line were near identical and the four replicate lines within each EE regimes were similar to one another (Fig 2). The multivariate repeatability (49) of SNP frequencies, which relates the disparity between the four replicate lines to the overall difference between E and L regimes, was high among both replicate E (*R* = 0.75; 95% bootstrap CI: 0.73 - 0.76) and L lines (*R* = 0.67; 95% bootstrap CI: 0.65 - 0.70), providing evidence for a high evolutionary consistency at the SNP level across replicates in the two regimes. This shows that the E and L regimes were more than twice as different from one another as were within-regime replicate lines from one another. Genome scans showed that all chromosomes exhibited multiple local regions with SNPs fixed or nearly fixed for alternative variants in E and L lines (Figs. 3, S7). To attest to the genome-wide nature of differentiation, we found 37,250 SNPs with dAF ≥ 0.9 between all E versus all L lines which were located in or near (±40 kb) 1,335 genes.

**Fig. 2.**
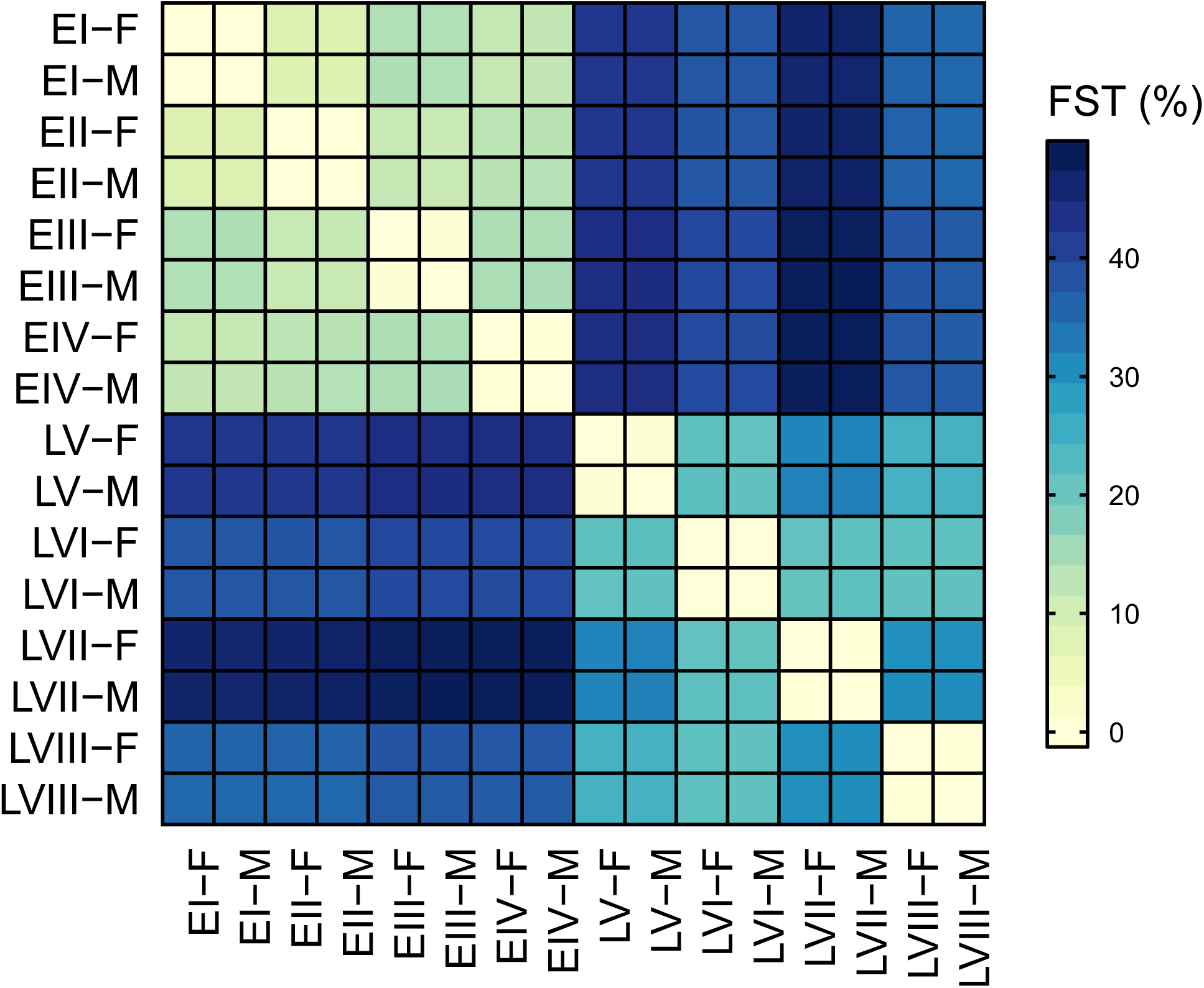
A matrix of pair-wise genetic differentiation (*F*_ST_) between samples from the E and L lines, based on the frequency of almost 8 M SNPs. Each of the eight lines is here represented by two biological replicate samples (denoted F and M).

**Fig. 3.**
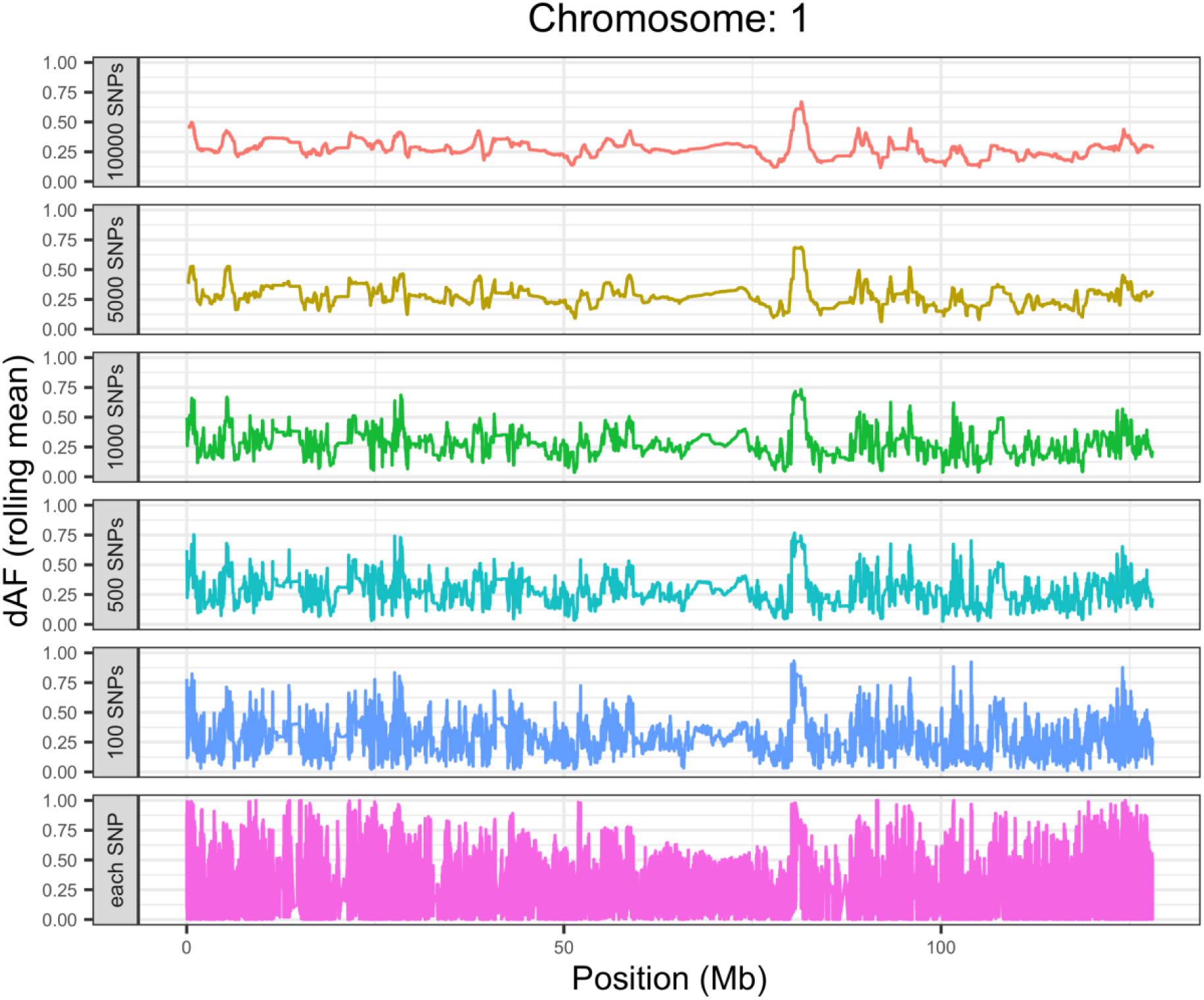
Differentiation between E and L lines was near genome-wide, here exemplified with a genome scan of the difference in SNP allele frequency (dAF) between superpools of E and L lines along chromosome 1, shown as a rolling average of various window sizes (window sizes bottom up: 1, 100, 500, 1000, 5000 and 10000 bp) (see also Fig. S7).

### Selective sweeps

On average 247 (SD = 65) regions, covering in total 129 Mbp (SD = 21), showed evidence of recent selective sweeps in the eight samples from E lines. The corresponding metrics in samples from L lines were 336 (SD = 131) and 135 Mbp (SD = 25). This corresponds to on average >10% of genomic coverage and regions called were located in all chromosomes.

Sweep calls showed positional consistency between sex-specific samples from the same line (ECDF ∼0.6 – 0.7). Within-regime comparisons showed moderate colocalization (ECDF ∼ 0.2-0.4), while inter-regime correlations were very low (ECDF ∼0 – 0.1) (Fig. S8). Importantly, the similarity in the spatial distribution of sweep calls between lines sharing EE regime was significantly higher than that between lines from different regimes (pseudo-*F*₁,₆ = 2.60, *R*² = 0.30, P = 0.027) and a PCoA ordination of sweep calls of all samples showed a very clear separation of E and L samples along PCo1 (Fig. S8) (based on the Jaccard similarity matrix of spatial distribution of sweep calls). We conclude that divergent selection between regimes, rather than differential effects of drift across replicated lines, is apparently a major determinant of our SNP based sweep calls.

We considered annotated genes overlapping a sweep in at least 6 out of 8 samples of an EE regime as candidates, excluding all blacklisted regions. A total of 656 and 136 genes were retained for E and L, respectively (39 overlapping), representing approximately 4.2% and 0.9% of the filtered gene universe and suggesting that sweeps in L lines targeted less gene-dense regions. Enrichment of GO terms are illustrated in Fig. S28-S34. In the short-lived lines, most enriched GO terms formed a large but cohesive cluster associated with the regulation of cell division, developmental processes, morphogenesis and metabolic processes. For example, several genes are involved in the regulation of the activity of GTPases, which controls pathways important for growth both at the cellular and organismal level. A more specific cluster is formed by proteins involved in RNA processing and translation, echoing the upregulation of RNA processing genes previously documented in the E lines (49). Overall, the pattern of functional enrichment is well aligned with the higher growth, developmental and metabolic rates seen in E lines (46).

The long-lived L lines showed a distinct pattern of enrichment in regions consistently under selection. Functional clusters involving lipid metabolism, stress response and behavior are consistent with phenotypic data showing that L lines have evolved to rely more metabolically on fatty acid oxidation (46), are more resistant to oxidative stress (48) and show altered reproductive behavior (50). Interestingly, a large cluster is associated with DNA repair and the G2/M phase transition in the cell cycle (Fig. S33 B). This is a major pre-mitotic checkpoint, where DNA and protein damage is assessed as well as mitotic spindle assembly and segregation, consistent with elevated selection for fidelity in long-lived lines.

### Copy-number variants

Multivariate ordination showed that E and L lines were markedly separated based on normalized genome-wide coverage profiles (Fig. S9). Moreover, regions showing differences in coverage between E and L occurred in all chromosomes (Fig. S10) and were distinct and localized.

By comparing coverage in samples from the E and L regimes (Fig. S11), we identified some 3-4000 putative CNVs in each EE regime, most of which were called as deletions relative to the other EE regime (Table 1). The average length was approximately 3-5 kb, with some calls reaching over 100kb (Fig. S12). In total, 238 functionally annotated and non-blacklisted genes resided in CNV regions in E lines and 363 in L lines, corresponding to approximately 1.46% and 2.22% of the filtered gene universe. Functional enrichment of these are illustrated in Fig. S28-S34.

**Table 1.** CNVs identified in each selective regime. Genes were considered affected if at least 50% of their length overlapped with a CNV and only if functionally annotated.

| Type<sup>a</sup> | E lines |  | L lines |  |
| --- | --- | --- | --- | --- |
|  | Deletions | Gains | Deletions | Gains |
| Number of calls<sup>b</sup> | 2552 | 449 | 3454 | 325 |
| Mean length (bp) | 4769 | 3645 | 4993 | 3173 |
| Number of affected genes<sup>c</sup> | 218 | 22 | 338 | 25 |
| Proportion of gene universe<sup>d</sup> | 1.33% | 0.13% | 2.07% | 0.15% |

<sup>a</sup>Relative to the other EE regime
<sup>b</sup>Filtered for significance, fold change and length.
<sup>c</sup>In non-black-listed regions.
<sup>d</sup>For filtered universe.

Enriched GO terms for CNVs in the E lines overlapped substantially with those highlighted in regions under selection, in that they include key regulators of cell division, developmental processes, morphogenesis, metabolic processes and transcription. For example, modulators of circadian rhythm and calcium ion transport are both key in cell-cycle regulation and the BMP signaling pathway is involved in multiple aspects of the regulation of development and growth. Two distinct clusters involve proteins that relate to reproduction, such as male genital development and sperm storage as well as the regulation of ovulation and egg-laying (Fig. S34 A and C). This is consistent with the fact that both post-mating sexual selection (51) and age-specific fecundity (52) differ between the E and L lines. Another striking enrichment is that involving mitochondrial ATP synthesis, given that the functional characteristics of the ETC complexes in the inner mitochondrial membrane is known to differ between E and L lines (44) and genes related to the ETC and to ATP synthesis have been found to be differentially expressed in E and L lines (49).

In the L lines, enriched GO terms for CNVs also showed apparent overlap with those identified in the sweep analyses presented above. Again, a major cluster is composed of genes related to mitotic fidelity, involving for example microtubule polymerization and organization as well as spindle assembly and chromosomal organization (Fig. S34 B). The expression of genes involved in mitotic spindle organization and assembly has previously been found to be upregulated in L lines (49) and has been associated with long life in other taxa (40). Other identified shared gene functions are the regulation of behavior and stress response. Enriched GO terms also included wound healing and immune response as well as protein ubiquitination, the latter being critical in maintaining protein homeostasis. A suite of genes related to immune function have previously been found to be upregulated in L relative to E lines (49). Finally, we note that several enriched functions, including response to oxidative stress and hypoxia as well as wound healing, overlap with the GO term “determination of adult lifespan” (Fig. S34 D).

### Intersecting functional signals of lifespan evolution

While there was significant overlap between regions showing signs of selective sweeps and those showing differentiated CNVs, this overlap was restricted and the genome-wide ECDF-based relative-distance correlation between sweep calls and CNV calls was generally low (dCor < 0.1; Fig. S38). To some extent this no doubt reflects the fact that CNV and sweep calls were both conservative, but also the standard exclusion of regions with atypical depth-of-coverage (DOC) in the SNP based sweep calls and the relative sparsity of well-supported SNPs in regions with atypical DOC.

Although the two types of regions thus harbour largely distinct gene sets, we wished to more formally assess any functional concordance between genes in sweep regions with those in regions showing CNVs, as well as with those previously found to show differential expression in E and L lines (49). We hypothesized that functional clusters overrepresented among the three gene classes (sweeps, CNVs and DEGs) may overlap. To test this hypothesis, we applied a custom functional analysis, that integrates multiple candidate gene sets (layers: selection-based candidates and expression-based candidates) within a single scoring framework (SI).

At a false-discovery-rate (FDR) threshold of 0.05, this approach identified 256 GO terms (restricted to “biological process”) significant in the E regime and 289 significant in the L regime, with an additional 98 terms significant in both regimes (“shared”). Clustering of significant GO terms by gene-set overlap yielded modules that showed a sizeable degree of functional overlap. A large set of functional clusters were identified as being associated with the EE of lifespan, which were enriched in more than one data layer and which involved a large number of functional categories canonically associated with a long life (Fig. S35-S37). At a functional level, thus, genes involved in selective sweeps overlapped considerably with those affected by CNVs and with those showing differential expression between E and L. For example, one functional cluster enriched in both E and L lines for sweeps, CNVs as well as for DEGs (Fig. S37 A) is associated with processes related to melanin biosynthesis. These are involved in the conversion of tyrosine into melanin, which is essential for both cuticle formation, pigmentation and hardening as well as for the phenoloxidase (PO) cascade. The former is essential for minimizing dehydration and the latter is key in both insect immunity as well as in oxidative stress tolerance (53), all with direct links to lifespan.

Further inspection of the genes that reside inside these functional clusters identified a very large number of multi-evidence candidate genes (Table S4), many of which have been implicated in ageing in other taxa. To highlight a few, two RecQ helicases (*RecQ4* and *RecQ5*) appeared in multiple data layers. These enzymes play central roles in maintaining genome stability and are linked to lifespan in both humans (54), *Drosophila* (55) and yeast (56). Several other candidates, such as *smc6*, *lig1*, *tam, ddb1* and *rad50*, also contribute importantly to genome integrity. Others include *ubiquitin*, a central component of proteostasis with well documented effects on aging (57), and a *yellow* gene which plays a central role in cuticle formation and hardening in insects and where loss of function causes dehydration and reduced lifespan in other beetles (58). *Adk1* is a core metabolic buffer enzyme, with likely effects on lifespan through its central role in cellular energy homeostasis and adenine nucleotide metabolism. An interesting candidate is also the heat shock protein *l2efl* (lethal 2 essential for life), as this key chaperone binds misfolded proteins and promotes proteostasis and stress resistance. It is located in a selective sweep in L lines, is markedly upregulated in expression in L lines and also shows a strong QTL signal (Table S4). Finally, *chico* is a central component of the insulin/IGF signalling pathway and a canonical lifespan gene (59) with documented effects on lifespan in several taxa (60).

### QTL mapping

To assess whether evolved genetic differences between EE regimes are associated with lifespan, we conducted a genotype – phenotype association study based on a large E/L intercross pedigree where 1334 recombinant F2s were genotyped and phenotyped for sex, lifespan and body size. The genome-wide additive QTL effect estimates of the L allele, from interval mapping (IM) models, show that markers overall exhibit directional effects concordant with expectations for both weight (mean additive L allele effect = 0.057 mg, SD = 0.045, SE =0.001) and lifespan (mean L effect = 0.51 days, SD = 0.42, SE =0.007). The distribution of the effects of the L allele across all markers was markedly skewed to positive values and often multi-modal across nearly every chromosome for both traits (Fig. S13-S16), consistent with a few loci with large effects and a very large number of loci with medium or small effects. Our IM models showed significant QTLs in four chromosomes (Fig. S26). QTL effects for weight and lifespan were positively correlated across all autosomes (r = 0.27– 0.93), suggesting wide-spread pleiotropic effects (Fig. S17).

The distribution of sex-specific effect estimates showed a similar pattern. Mean effects of the L allele were positive for both males and females for weight (males: 0.049 mg, SD = 0.047, SE =0.001; females: 0.051 mg, SD = 0.062, SE =0.001) (Fig. S18) and lifespan (males: 0.32 days, SD = 0.38, SE =0.006; females: 0.60 days, SD = 0.55, SE =0.009) (Fig. 4). Genome-wide correlations of QTL effects across all markers between sexes were quite high for both weight (r = 0.49) and lifespan (r = 0.60) (Fig. S21-S22). Yet, overall QTL effects were stronger in females than males for lifespan (paired *t* = 25.16, df = 6608.2, *P* < 0.001) but not for weight (paired *t* = 1.39, df = 6956.9, *P* = 0.165). Within individual chromosomes, the correlation between male and female QTL effects varied substantially for weight (r = 0.16– 0.77) and even more so for lifespan (r = −0.70–0.88) where chromosome 1 and 7 even showed a negative correlation (Fig. S23-S24). Unlike the moderate-to-high positive correlations between weight and lifespan in sex-averaged models, sex-specific correlations were generally high in most chromosomes in females but were lower overall in males (Fig. S25).

**Fig. 4.**
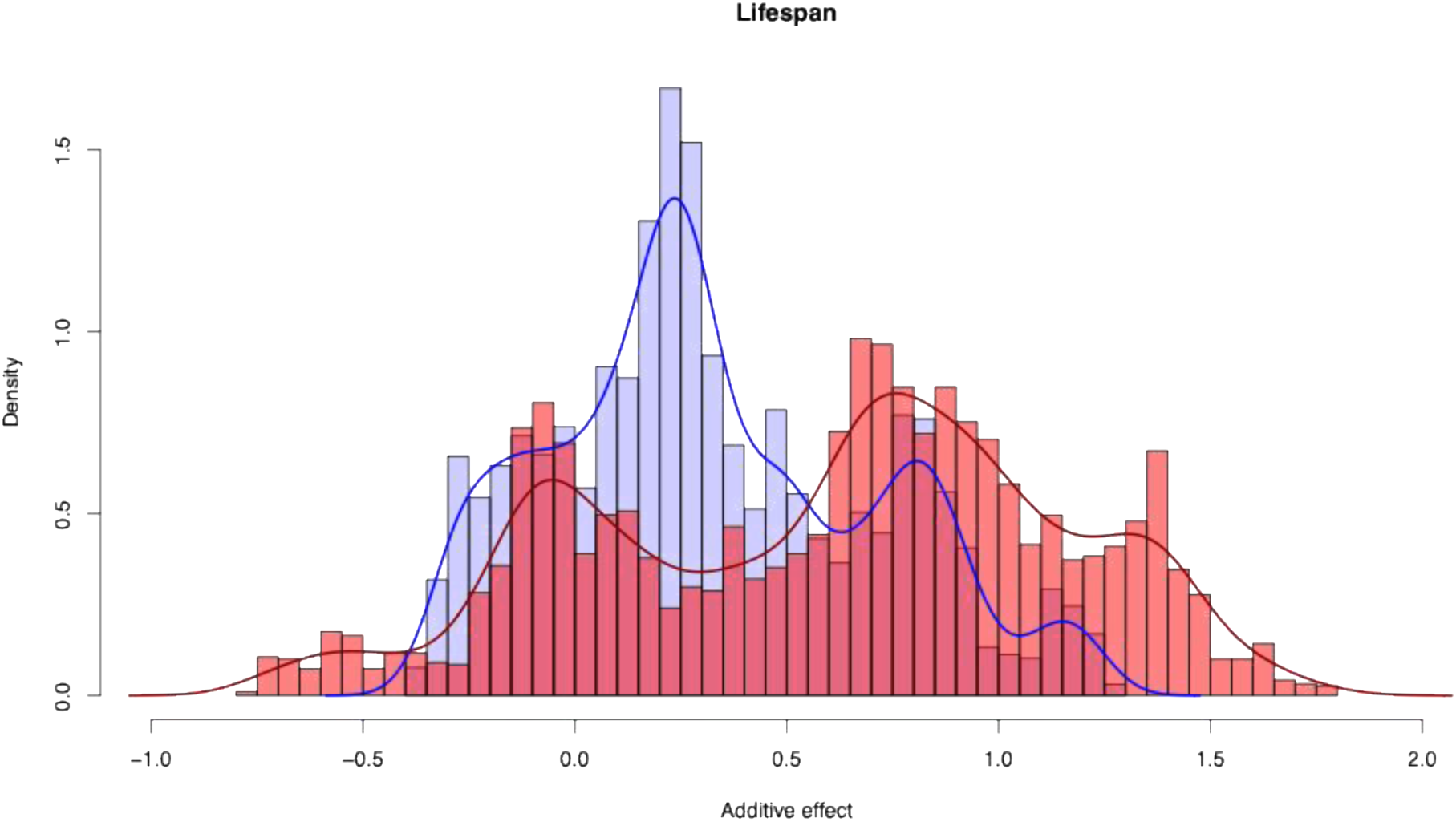
Frequency distributions of the additive effects of the L allele for lifespan (in days) in males (blue) and in females (red) for all QTL markers, estimated separately for the sexes in interval mapping models. Positive values mean that the L allele is associated with a long lifespan. Distributions are displaced from zero and a very large number of markers show medium and small effects on life span.

Standard interval mapping with sex as an additive covariate identified QTLs for body size on chromosomes 2, 4, and 9 (Fig. S26-S27). Here, multiple markers exceeded the permutation-based LOD threshold on each chromosome, spanning variable proportions. For example, chromosome 2 had significant markers spanning ≈70% of the chromosome, while chromosome 4 markers spanned only ≈4% of the chromosome. Lifespan QTLs showed a somewhat similar pattern, with peaks on chromosomes 2, 4, and 9 but also showed two distinct marginal peaks on chromosome 7 (Fig. S26-S27). Sex-specific QTL models revealed QTLs unique to each sex and sometimes distinct from those identified by the sex-averaged model (Fig. 5). For body weight, females showed a single LOD peak on chromosome 2, while males had peaks on chromosomes 2, 3, 4, and 9. For lifespan, LOD peaks also differed in position but appeared on chromosomes 2, 4, and 7 in both males and females. A striking case is chromosome 7, where the sex-averaged model showed relatively marginal effects on lifespan (Fig. S26) but where QTL effects were negatively correlated across sexes (r = −0.7; Fig. S24) and the sex-specific models revealed two clear sex-specific peaks located on opposite terminal regions of the chromosome (Fig. 5).

**Fig. 5.**
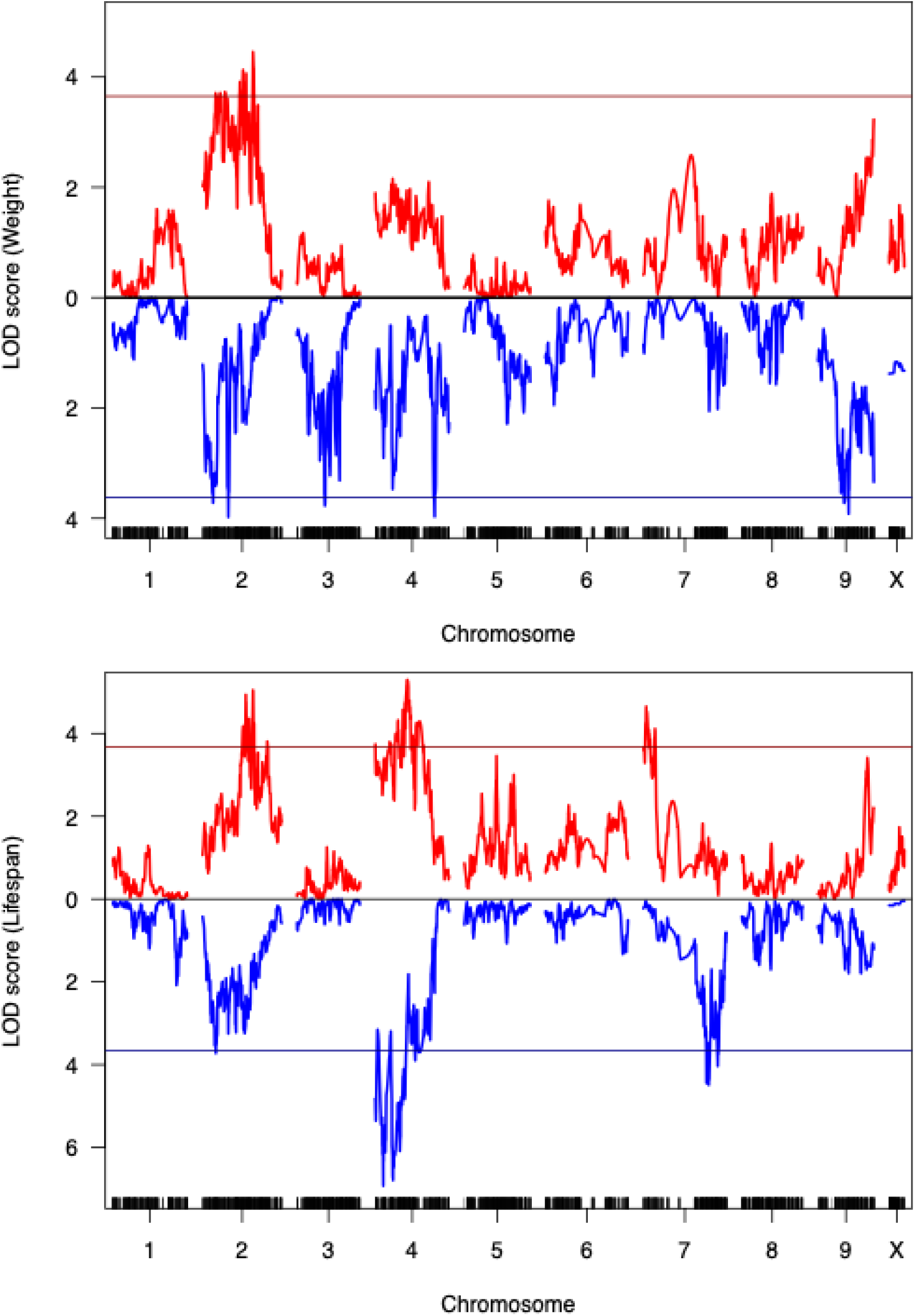
Sexual dimorphism in the genomic architecture of body size and lifespan. The figure shows LOD scores for body weight (top) and lifespan (bottom) plotted for both males (blue) and females (red), from interval mapping models based on 1334 recombinant F2 individuals. The solid horizontal line denotes the LOD threshold determined by permutation tests for autosomes. Females showed a single significant LOD peak for weight, while males had five significant peaks. For lifespan, LOD peaks were somewhat more concordant between sexes, although differing in position, appearing on chromosomes 2, 4, and 7 in both sexes.

Multiple QTL mapping (MQM) models with sex as an interactive covariate are better able to account for linked effects, thus increasing LOD peak resolution (61). These models detected QTLs not identified by standard interval mapping and confirmed the presence of multiple QTLs on several chromosomes. Six significant QTLs for body weight were identified on chromosomes 2, 4, 6, and 9, collectively accounting for 8.2 % of sex-specific variance in body weight. Six significant QTLs for lifespan, collectively accounting for 10.0 % of the sex-specific variance in lifespan, were located on chromosomes 2, 4, 5, 6, 7, and 9 (Fig. 6). While none of the QTLs for body weight showed a significant interaction with sex, four out of six lifespan QTLs exhibited significant interactions with sex (on chromosomes 2, 5, 7, and 9) (Table S3). These MQM models thus confirm a polygenic architecture of these traits as well as important differences between the sexes in terms of genetic effects on lifespan.

**Fig. 6.**
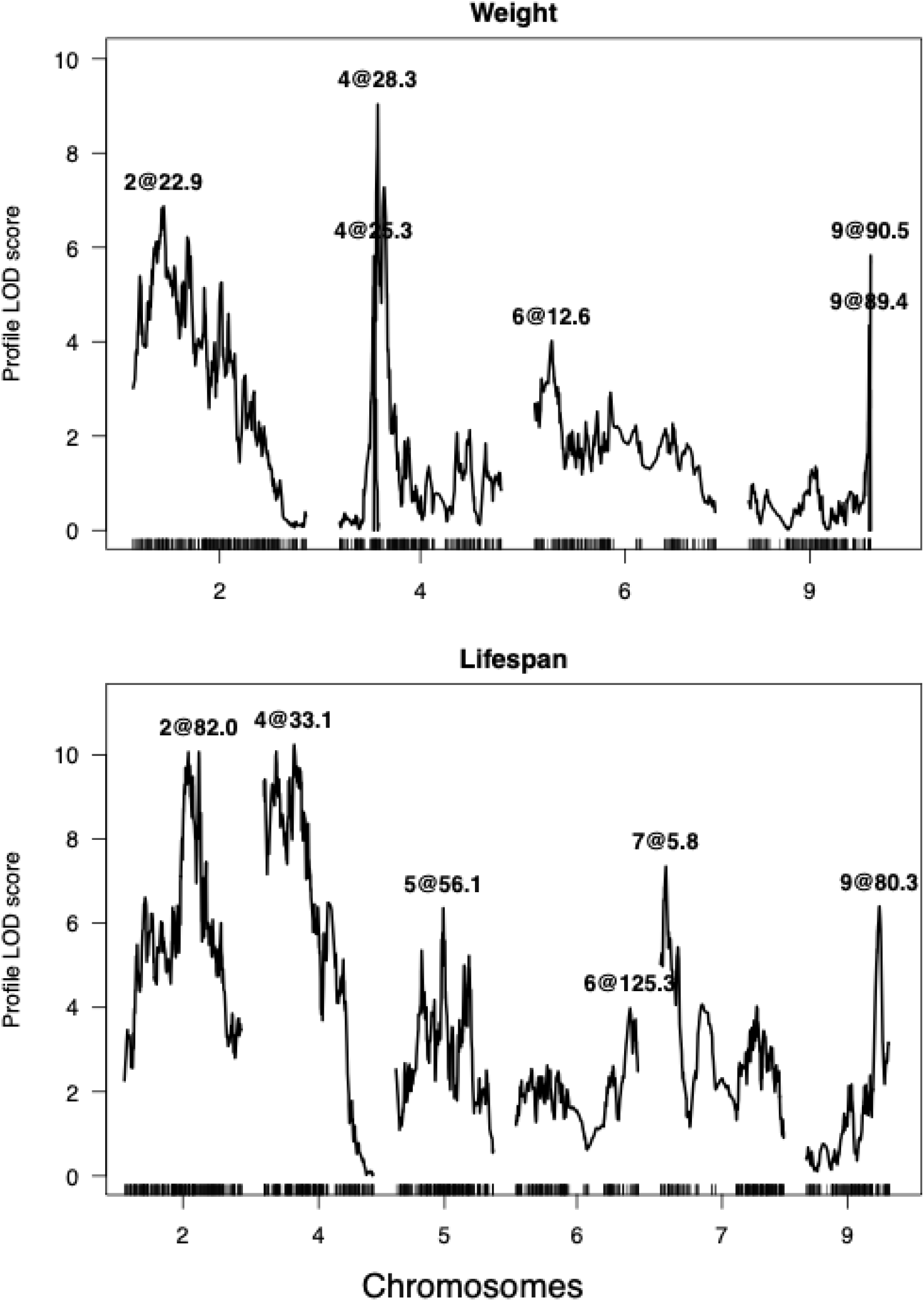
The LOD profile generated by multiple QTL mapping models. Significant QTLs are denoted by their chromosome and position (in cM). Here, LOD profiles for each QTL are calculated by comparing the full model with the model in which the QTL of interest is omitted.

The six top lifespan QTLs were further inspected for overlap with CNVs, DOC in F0 pedigree founders, selective sweeps, recombination rates and differential gene expression (49). The region delineated by QTL 2@82.0 showed several CNVs and the DOC was clearly divergent among F0 founders. The region harbours the protein draper, which is upregulated in E females and involved in phagocytosis and apoptosis. The QTL 4@33.1 carries a 1.2 kbp deletion in L lines and is flanking several regions exhibiting selective sweeps. This deletion is also present in the F0 L founders. This region harbours the galactose-1-phosphate uridylyltransferase gene, a key enzyme in galactose metabolism, which is also upregulated in the abdomen of E females relative to L females. The QTL 5@56.1 shows two selective sweep regions in L, and there are two flanking major deletions in E lines (150 kbp and 135kbp) involving several annotated genes, as well as several smaller deletions in L lines. The E and L F0 founders also differ markedly in the region with these CNVs. This region covers several DEGs, such as elongation of very long chain fatty acids protein 1 and fatty acyl-CoA reductase. The QTL 6@125.3 is a large (5Mbp) but well-defined peak, with a number of relatively small CNV calls and a proximate selective sweep region in E lines. The E and L F0 founders clearly differ in DOC in this region. It harbours 9 DEGs, among these DNA primase large subunit and two copies of pachytene checkpoint protein 2. The latter protein monitors double-strand-break events and is key in meiotic recombination checkpoint signaling. The absence of recombination events within this QTL but a high recombination rate in the flanking regions leads us to hypothesize that it represents a chromosomal inversion. The QTL 7@5.8 holds three deletions in the L lines (3-5 kbp) and an adjacent sweep region in L lines. It harbours four DEGs, of which two are involved with transmembrane ion signalling (ionotropic glutamate receptor activity, piezo-type mechanosensitive ion channel component) and one is a mitochondrial tricarboxylate transport protein. Finally, the QTL 9@80.3 is 2.5 Mbp region showing a number of CNVs and a clearly divergent DOC in the E and L F0 founders. A region close to this QTL was called as sweep region in E lines. It carries one DEG, an orthologue of tetratricopeptide repeat protein 39B. In summary, the major QTLs were structurally complex, most were either close to or enclosed regions showing sweeps, harboured many CNV calls in the EE lines or showed contrasting DOC profiles in E and L F0 founders. Moreover, the QTLs harboured DEGs annotated as being involved in key phenotypes such as lipid metabolism, carbohydrate metabolism, cell cycle regulation and DNA repair.

### Intersecting QTLs with signals of evolutionary differentiation

To assess whether variation in the strength of the genotype–phenotype association (represented by the LOD score from our global IM model) was related to evolutionary differentiation between E and L (represented by our focal regions: selective sweep and CNV calls) across the genome, we performed a series of spatial correlation analyses using StereoGene (62). These genome-wide correlations summarize large-scaled rather than regional patterns of covariation, which means that they are not simply built by a small set of large-effect sites, and tests for non-random associations using spatial permutation tests. For lifespan, the mean genome-wide correlations were 0.328 (SE = 0.014) [0.249] for CNVs in E lines, 0.363 (SE = 0.015) [0.298] for CNVs in L lines, 0.297 (SE = 0.027) [0.273] for sweeps in E lines and 0.249 (SE = 0.032) [0.296] for sweeps in L lines [partial correlation after accounting for genome-wide covariance with mappability] (all *P* < 3.42 × 10^−14^). For body size, the correlations were 0.33 (SE = 0.014) [0.236] for CNVs in E lines, 0.371 (SE = 0.015) [0.281] for CNVs in L lines, 0.299 (SE = 0.027) [0.222] for sweeps in E lines and 0.249 (SE = 0.032) [0.24] for sweeps in L lines (all *P* < 2.67 × 10^−14^).

The overlap between CNV and sweep calls was not as high as one might expect (see above), and their spatial correlation was also restricted (0.286 [SE = 0.024] in L lines and 0.292 [SE = 0.021] in E lines). Yet, proximity between focal regions could generate confounding effects. We thus also estimated partial spatial correlations between focal regions and LOD score, where the influence of one focal region on the correlation between LOD score and the other focal region was removed. These partial correlations were ∼10% lower than the bivariate correlations for CNVs and LOD score, while the partial correlations between were ∼50% lower for sweep calls and LOD score. Although these results should be interpreted with some caution, as the sweep and CNV tracks differ in densities and interval-size distributions, they are least suggestive of a stronger direct association between CNVs and LOD score than between selective sweep calls and LOD score. In summary, the spatial correlations between genome tracks were sizeable. The correlations were much larger than expected by chance, and estimates of direct associations were stronger between LOD score and CNV calls than between LOD score and sweep calls.

## Discussion

Lifespan represents a key phenotype in life history research and is also of substantial interest in applied sciences and medicine. Here, we use experimental evolution (EE) in a comprehensive effort to shed light on the genetic architecture of evolved differences in lifespan and assess the role of structural variation (SV) and sex-specificity. We address five main points. First, the degree of complexity *per se* is of importance to the field. Our findings expose a highly polygenic genetic basis for evolved differences in lifespan in a non-model organism. Second, although it is becoming increasingly clear that SVs can form a major component of segregating genetic variation, the potential general role of SVs in adaptation is still an open question. We show that CNVs are associated with the EE of lifespan, implying that SVs contribute significantly to the adaptive evolution of life histories. Third, classic models for the evolution of ageing make distinct predictions about which classes of genes and variants should be involved, and we evaluate our results in light of these predictions. Fourth, the extent to which the genetic basis for adaptive evolution in complex traits is repeatable is debated, and we use our results to shed light on this issue. Finally, studies of the biological basis of ageing often ignore sex. We show that the trajectory of lifespan evolution differs between sexes, as does the genetic architecture of the evolved difference in lifespan. This has implications for sex-specific life history adaptation but also for the evolution of ageing more generally.

### Genomic complexity

The genetic architecture of adaptive lifespan evolution has been characterized in *D. melanogaster*, where three independent EE experiments each identified > 1,000 candidate genes (38–40). Much less is known about the molecular genetic basis of lifespan in other insects (42). We found that in *A. obtectus*, the evolved difference in lifespan between the EE regimes was associated with SNP differentiation in or near some 1,300 genes. Although linked selection is no doubt responsible for a considerable fraction of this highly polygenic signal, we note that recombination rate was normal in *A. obtectus*, the duration of the experiment was very long and differentiated regions were distributed along all chromosomes. Further, the selective sweep analyses identified some 750 genes in regions under consistent selection across replicate lines and our QTL mapping revealed a polygenic basis for evolved differences in lifespan. Although only a handful of QTL peaks were statistically significant in isolation, the distribution of allelic effects was consistent with the presence of a very large number of small and medium effect QTLs. Importantly, the genome-wide spatial correlation between LOD scores and evolutionary differentiation between E and L confirmed this, as this is built by genome-wide effects rather than regional effects of a few large-effect variants (62). The nature of this correlation is thus consistent with the interpretation that a very large number of QTLs with small or medium sized effect are causal, despite being statistically non-significant, as they were also spatially associated with differentiation between E and L. Thus, it is clear that the phenotypic effects of EE seen in *A. obtectus*, including a near doubling of lifespan, is due to evolutionary modifications involving at the very least many hundred genes. Overall, the high complexity of the genetic architecture is thus on par with that observed in *D. melanogaster*.

Genomic complexity is not only built by polygenic inheritance, but also to a high degree by patterns of epistasis, gene-by-environment interactions and pleiotropy. While such effects are very difficult to fully characterize with sequencing data (63–65), our QTL mapping provides a foundation for assessing the degree of pleiotropy between lifespan and body size. We found that the correlation between genetic effects on lifespan and body size across all markers was sizeable across all autosomes. This implies that many genetic effects on functions such as metabolism, growth and development generate downstream effects on both key life history traits, adding to the complexity of both. Yet, we note that genetic effects are not entirely shared as the major QTL peaks for the two traits were distinct, and pleiotropic constraints are thus to some extent restricted. Finally, we note that earlier quantitative genetic studies in less advanced generations of our EE lines not only provided evidence for additive genetic variance for lifespan (52) but also revealed a substantial amount of epistatic variance (64). While QTL mapping studies such as ours are not ideally suited for detecting and characterizing complex epistatic interactions, due to a near infinite QTL model universe and thus limited statistical power (66, 67), the results of our modelling bore hallmarks of epistasis. Most notably, the genotype-phenotype map from the interval mapping was in several cases altered when including information on genotype in markers in other chromosomes (i.e., in our MQM models), as predicted when interactions across linkage-groups affects lifespan. These findings underpin the general conclusion that lifespan is a highly complex trait.

### Structural variants

It has been predicted that segregating genetic variation in CNVs should provide important raw material for adaptive evolution (8, 9). Although a role for CNVs in adaptation has been uncovered in EE experiments in single-cell organisms (68, 69), very few EE studies have addressed this possibility in metazoans (70) and none have systematically gaged the effects of CNVs for lifespan evolution.

We detected in total several thousand of regions showing putative CNVs that showed consistent differentiation between E and L lines, with marginal overlap with sweep regions. These CNV regions harboured several hundred annotated genes and enrichment analyses revealed an overrepresentation of functional categories (i) generally implicated in the evolution of ageing in model systems, (ii) matching phenotypes known to be divergent between long- and short-lived lines and/or (iii) known to show differential expression in our EE regimes. These observations lend considerable support to an important role for at least a fraction of the CNVs in mediating life history evolution in our species. Moreover, the functional overlap between genes in differentiated CNV regions and those earlier shown to exhibit differential expression (49), including for example ATP synthesis, stress response, immune function and mitotic spindle organization and assembly, suggests that the effects of CNVs seen at least in part involves effects on transcription. CNVs can affect gene expression in several distinct ways (71) and associations between CNVs and expression have been documented through e.g. the mapping of CNVs as eQTLs (72) as well as through experimental approaches (73, 74). The observation that the CNV regions detected in our study were relatively gene-poor, in conjunction with the fact that the genome of *A. obtectus* is very repeat-rich (>60%; 49), further suggests that the effects of many CNV regions involve regulatory effects. Our inspection of the major QTLs substantiated this suggestion, as QTLs contained both structural complexity and differentially expressed genes. This aligns well with recent work in humans, showing that a large fraction of the total genetic variation in many complex traits is caused by non-coding variants with effects on gene expression (75).

More generally, our analyses suggest that at least 3% of the filtered gene universe in the wild population used to found our EE lines is residing in segregating CNVs with potential effects on lifespan, and including genes nearby but not within CNVs would result in a greater fraction. This implies that a significant fraction of genetic variation for this life history trait involves SVs. It should be noted that the pipeline used here for detection of CNV regions involved in the evolution of lifespan, based on contrasting depth-of-coverage information in the two EE regimes, is relatively conservative. Yet, although calling CNVs based on depth-of-coverage in ordinary short-read data may not be the most sophisticated method available, it has been shown to perform well relative to other pipelines (76, 77) and can be applied to poolseq data (78). The fact that the depth-of-coverage profile to a large extent was divergent between EE regimes, but consistent across replicate populations within regimes (Fig. S9), places considerable confidence in the pipeline used.

Our analyses of the links between the strength of the genotype–phenotype association across the genome and differentiation between E and L lines provided additional insights. First, the spatial correlation between LOD score and sweep/CNV calls was sizeable. This provides evidence that regions with evolutionary differentiation tended to be those with phenotypic effects on lifespan, as expected under adaptive EE. Second, the genome-wide nature of these correlations (62) shows that a large number of QTL markers with phenotypic effects harbour CNVs. Third, given that the overlap between regions showing signs of consistent selective sweeps and those showing differentiated CNVs was marginal, the fact that the genome-wide correlations between LOD score and CNV calls was if anything higher than those between LOD score and sequence-based sweep calls suggests that the effects of structural variants on the adaptive evolution of lifespan are at the very least on a par with those of sequence variants. Our work thus implies that the genetic architecture of adaptive evolution of lifespan is built by structural variation to a large extent. Yet, we stress that a precise quantitative comparison of the relative importance of sequence variation and SVs is difficult, because CNV and sweep calls are by necessity based on different methods and interrogate different genetic compartments.

### The evolution of ageing

Virtually all organisms age at some rate and fully understanding the evolution of ageing and restricted lifespan remains an ongoing challenge in evolutionary biology. Ageing results from the fact that the strength of natural selection on survival declines with adult age (79), which in turn results from age-independent extrinsic mortality. There are two non-mutually exclusive corollaries of this. One is age-specific antagonistic pleiotropy (80): alleles or traits that are beneficial early in life but deleterious late in life may be favoured, because selection is stronger early in life. This antagonism involves trade-offs either between functional effects on various life history traits or between competing demands in resource investment to reproduction and somatic maintenance (81). The other is that any alleles or mutations with detrimental effects expressed only late in life may accumulate in a population, simply because natural selection is too weak to eliminate them (82).

While it is generally difficult to determine the relative contribution of antagonistic pleiotropy and mutation accumulation to aging, EE can offer important insights (37). Indirect phenotypic responses of the EE of lifespan involving symptomatic trade-offs, for example a reduction of early life fecundity in long-lived lines, is generally interpreted as evidence for antagonistic pleiotropy (3). In *A. obtectus*, previous assays of indirect phenotypic responses of our EE have documented effects on several life-history traits involved in trade-offs, such as early life fecundity, growth, body size, metabolic rate and resistance to stress. These observations point to a central role for antagonistic pleiotropy.

The functional genetic architecture of evolved differences in lifespan can also offer crucial insights (3, 35, 40). The EE of lifespan under antagonistic pleiotropy is predicted to target genes enriched for pathways involved in canonical life history trade-offs (83). In contrast, mutation accumulation does not predict enrichment for any particular life-history function, and affected genes are expected to scatter across random cellular processes (3). Three aspects of our findings support a major role for antagonistic pleiotropy. First, the identity of enriched functions included processes involved in a spectrum of canonical life history trade-offs. This included growth, developmental and metabolic rates, as well as immune function, regulation of reproduction, maintenance and DNA fidelity. Second, the overlap in enriched functional categories across different evidence layers (gene expression, selective sweeps, CNVs, QTLs) in itself implicate selection and canonical life history trade-offs in lifespan extension. Third, the high genomic repeatability of lifespan evolution across replicate EE lines also points to antagonistic pleiotropy, as mutation accumulation is predicted to result in a more pronounced role for stochasticity: mutations neutral early in life would be free to differentiate in replicate populations through drift (3). This is especially true for E lines, where the effects of drift should be more pronounced due to the fact that E lines underwent more generations but where repeatability instead was largest.

Our multi-evidence enrichment analyses show that a large set of biological processes underpin the EE of lifespan in *A obtectus*. There are some interesting overlaps with similar EE studies in *Drosophila*. A dedicated search for homologs in *A. obtectus* of the 20 genes that are candidates in all three *Drosophila* EE studies (40) recovered 10 homologs. Two of these, FBgn0261555 (CG42673) predicted to mediate nitric-oxide synthase binding and FBgn0083981 (RunxA), have both been associated with oxidative stress resistance in *Drosophila* (84) and are both associated with CNV variants in the *A. obtectus* genome. This may contribute to the higher oxidative stress resistance seen in our L lines (48). Yet, although many higher order functions are shared with similar studies in *Drosophila*, the specific pathways sometimes differ. For example, while genes implicated in immune function play an important role both *Drosophila* and in *A. obtectus*, the effects are mainly mediated by the Toll signalling pathway in *Drosophila* (40) but primarily by other immune-related pathways in *A. obtectus* (49). Similarly, while nutrient sensing pathways that affects growth, such as the Insulin-like Growth Factor and the Target of Rapamycin signalling pathways, are key in *Drosophila* (39, 40), other pathways that regulate growth, development and reproduction are more central in *A. obtectus*. These observations suggest that although many key life history trade-offs are shared, the precise evolutionary trajectories differ across species. This likely reflects differences in ecology and physiology between taxa. For example, adult seed beetles are aphagous capital breeders and the EE of a long life is associated with the growth of a larger adult body size (85) and a higher lipid content (47) in *A. obtectus*. This allows more efficient storage of long-term energy resources and lower water requirements (46), both of which are important for a long life in warm and dry conditions. In contrast, *Drosophila* are to some extent income breeders and nutrient sensing pathways are thus more central terms of food consumption, digestion and metabolic requirements (86), and the EE of a long life shows no or little association with body size (87, 88).

The extent to which adaptive evolution involves deterministic or contingent forces is a long-standing issue in biology. Evolve-and-resequence experiments from standing genetic variation such as ours allow direct assessments of the degree of repeatability of adaptive evolution (36). A common prediction is that the genomic repeatability of shifts underlying complex traits such as lifespan should be low, because of genetic redundancy (89, 90).

Somewhat surprisingly, we found that the multivariate genome-wide repeatability of SNP frequency differentiation between our E and L lines was high (*R* = 0.67 – 0.75), on par with that observed earlier for gene expression (*R* = 0.56 - 0.72; 49). Moreover, a large number of genomic regions showed signals of selective sweeps and of CNV differentiation which were consistent across replicate lines. The repeatability observed for both SNPs and CNVs is, no doubt, in part due to linked selection. Yet, we note that recombination rate in *A. obtectus* was normal (genome-wide average 1.07 cM/Mb), our experiment lasted for 329 (E) and 213 (L) generations, all chromosomes showed differentiation and differentiation correlated with phenotypic effects on lifespan genome-wide, suggesting that the effects of linked selection is predominating. Our functional analyses of both sweeps and CNVs, showing that focal regions were enriched with genes involved in canonical life-history traits, instead suggest that deterministic forces can have predictable effects on the genomic architecture even of complex polygenic life history traits.

### Sex-specificity of lifespan evolution

Among our EE lines, females have diverged more between regimes in lifespan (Fig. 1), rate of senescence (85) and gene expression (49). This implies that the genetic architecture of lifespan must be at least partly private to the two sexes, allowing the rapid evolution of sexual dimorphism in lifespan, despite a fairly high genetic correlation between lifespan in the two sexes both in general (91) and in seed beetles (*r*_mf_ = 0.33 – 0.83; 28, 30). The intersexual correlation between the effects on lifespan across all QTLs in our analyses (*r*_mf_ = 0.60) is well aligned with these findings (Fig. S22). Yet, we detected distinct major QTL peaks in four chromosomes with significantly different effects in males and females and the LOD profile was distinct in several chromosomal regions. Here, chromosome 7 is especially interesting. While this chromosome harboured multiple QTLs for lifespan in both sexes, the regions of the two sex-specific QTLs are non-overlapping (Fig. 5) and the genetic effects on lifespan were even negatively correlated across sexes (Fig. S24). These results echo a handful of previous QTL mapping studies of lifespan in *Drosophila*, of which a few used recombinant inbred lines (32, 92, 93) and one used EE lines (94), all of which documented QTLs with sex-specific effects. In our case, the fact that clusters of sex-specific QTLs are located on opposite arms of chromosome 7 suggests that genomic rearrangements and sex-specific adaptation may have generated genomic “neighbourhoods” with sex-specific effects on lifespan (95).

A genetic architecture that allows sexual dimorphism to evolve is built by loci with sex-specific phenotypic effects. However, very little is known about the loci that facilitate sexual dimorphism in lifespan (6). A closer inspection of the genomic structure of the four QTLs with significantly different effects in the sexes, in conjunction with previous work on gene expression (49), suggest that the effects seen occur through sex-biased genes. For example, the central part of QTL 2@82.0a in chromosome 2 shows a 1.6 kbp region identified as both being under a selective sweep and carrying a CNV. A closer inspection of DOC in this region shows that it represents a CNV gain in all replicate L lines, as well as in the L F0 individuals, and the pattern of mapped paired-reads is consistent with a series of tandem duplications of non-coding DNA (Fig. 7). This region is located downstream of an EF-hand domain-containing family member gene, which is strongly and significantly differentially expressed in E and L beetles: its expression in the abdomen of young females (but not in males) is approximately 100 times higher in L lines compared to E lines. This protein acts as a Ca²⁺ sensor pivotal to calcium signaling, such as the calcium-dependent calcineurin signaling cascade. Calcineurin is known to promote the induction of innate immune responses in *Drosophila* (96) and innate immune function is indeed much elevated in females in seed beetles (97) which may contribute to the sex specific effect on lifespan. However, calcium signaling and calcineurin are also key regulators of egg activation and oocyte maturation (98) and an even more likely effect is that an increased expression of this gene in young L females acts to delay egg production and thus to prolong life. In both cases, however, evidence suggests that the evolution of sexual dimorphism in lifespan is due to a structural variant with effects on the expression of a gene with sex-specific/limited phenotypic effects.

**Fig. 7.**
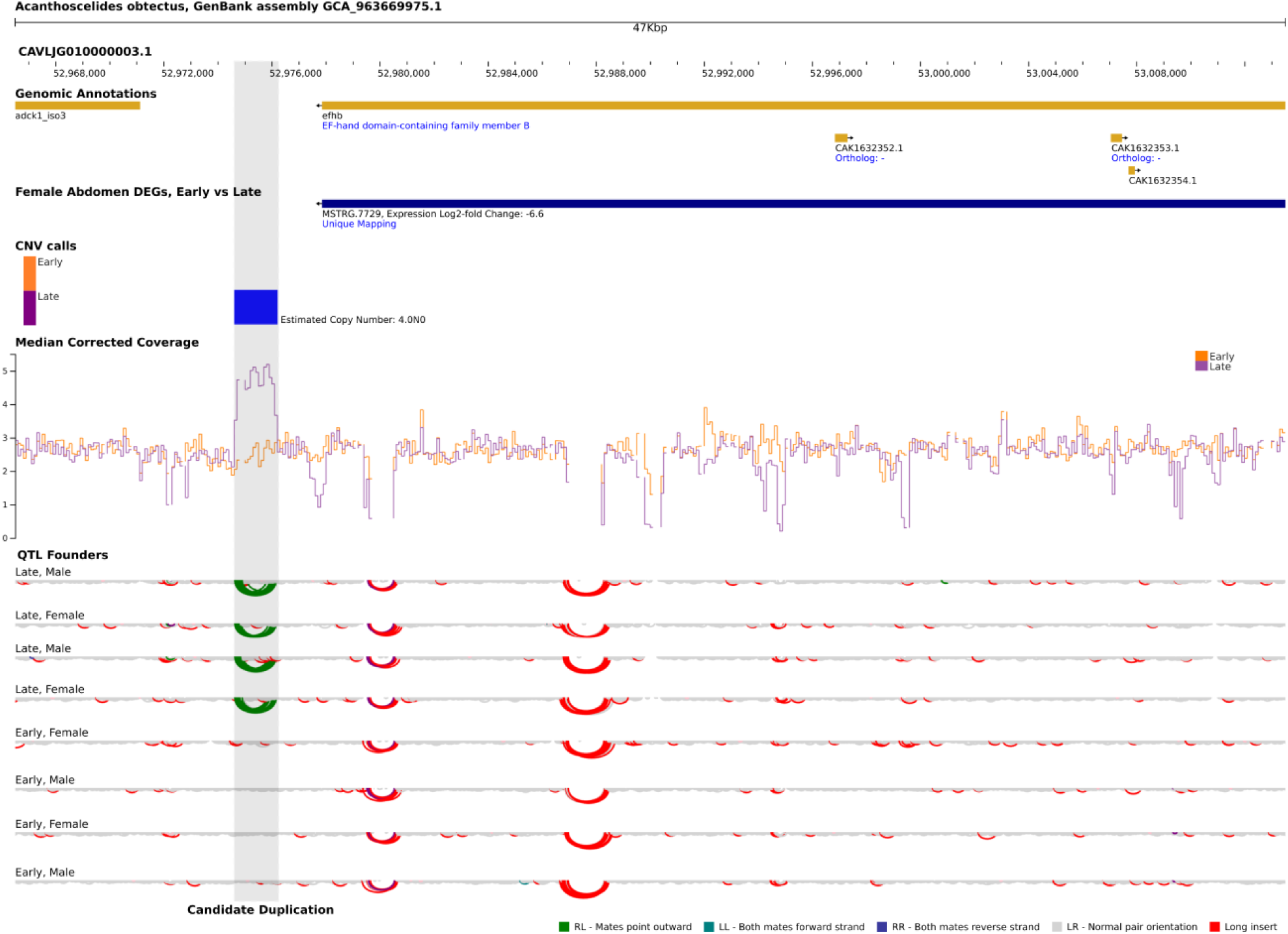
Candidate tandem duplication at QTL 2@82.0. This QTL, identified by the multiple-QTL mapping (MQM) model, shows sex-specific effects on lifespan. The shaded region marks a putative duplication that colocalizes with the QTL peak, supported by increased copy number and sequencing coverage in the L lines, as well as by discordant read-pair patterns in the L pedigree founders consistent with a tandem duplication. The region is also centred on a selective sweep detected consistently in the L but not the E lines (not shown in the plot). The duplication lies 1.6 kbp downstream of a uniquely mapped EF-hand domain-containing gene (MSTRG.7729), which is strongly differentially expressed between selection regimes in females (Early vs Late: log₂FC = −6.6) but not in males. Collectively, these observations identify this region as a candidate contributor to the sex-specific lifespan effect at this QTL. Females homozygous for this tandem duplication show a lifespan extension of 11% (Table S3).

Vieira et al. (27) suggested that the antagonistic pleiotropy hypothesis of ageing can be extended to sex-antagonistic effects. Sexually antagonistic (SA) alleles are those with opposing fitness effects in males and females (20) and this could in theory affect the evolution of lifespan in two ways (99). First, allelic effects on lifespan may be shared by the sexes but optimal lifespan may differ. Second, allelic effects on lifespan may be opposing in the sexes, which was in fact true for chromosome 1 and 7 in *A. obtectus*, but optimal lifespan shared. In both cases, allelic effects on fitness will be opposing in the two sexes and adaptive evolution of lifespan in one sex will be constrained by selection in the other. Needless to say, the potential for SA pleiotropy to affect lifespan evolution is expanded when considering pleiotropic effects on other life-history traits and the effects of sexual selection (18, 99). Variants that increase early life reproductive success in one sex may, for example, reduce lifespan in the other.

Although genes with SA effects are very prevalent (100), the degree to which SA pleiotropy constrains the evolution of lifespan is generally difficult to assess empirically (99, 101). In our experiments, our EE conditions have distinct effects in males and females and this has left footprints in sex-specific evolution of lifespan. In our E lines, male and female fitness is gained exclusively during day 1-2 and selection will favour a live-fast-die-young life history in both sexes (18). In our L lines, where oviposition is allowed only after day 10, females must survive to day 10 to leave any genetic contribution to the next generation. Because males and females are allowed to mate already from day 1 and female seed beetles store sperm for extended periods (102), males can mate and gain reproductive success already from day 1. This means that optimal lifespan is shifted upwards more in females than in males under our L conditions and we predict that selection in males will constrain the evolution of a long life in females. The sex-specific response to EE (Fig. 1) not only shows that the underpinning of lifespan is partly private to the two sexes but it is also consistent with this prediction, in line with a role for SA pleiotropy in constraining the adaptive evolution of lifespan during the course of our EE.

Lifespan generally exhibits substantial amounts of genetic variance (103), which is not expected if alleles that prolong life would be unconditionally beneficial and work on *Drosophila* have shown wide-spread context-specificity of genes affecting lifespan (6, 7). Antagonistic pleiotropy through functional trade-offs between different life history traits, such as early-life fecundity and late-life survival, can generate net balancing selection and thus contribute to the maintenance of genetic variation in longevity (104). Importantly, the conditions under which such balancing selection maintains variation are expanded when selection on different life history traits differ between the sexes (105) and Vieira et al. (27) suggested that SA antagonistic pleiotropy involving lifespan may contribute to the maintenance of genetic variation in loci affecting lifespan. Our results are broadly consistent with such a scenario. Segregating variation in loci affecting lifespan was evidently considerable in the wild population used to found our EE lines and selection on lifespan is known to be sex-specific in these beetles (101, 106, 107). It is therefore likely that SA antagonistic pleiotropy contributes to the maintenance of genetic variation in regions affecting lifespan, and our results moreover suggests that a fraction of these involve structural variants.

## Conclusions

We amalgamate long-term EE and QTL mapping to dissect the genomic architecture of adaptive lifespan evolution in an insect model system. We uncover a highly complex polygenic architecture, involving many hundreds of regions throughout the genome, with clear evidence of both pleiotropy and epistatic interactions across regions. Genes within candidate regions were consistently enriched for canonical functional modules, such as DNA fidelity, immunity, stress tolerance, protein maintenance, circadian rhythm, mitochondrial function, reproduction, growth and metabolism. Collectively, our results show that SVs are important for the adaptive evolution of lifespan, possibly on par with the importance of sequence variants. Moreover, the genomic architecture of lifespan differed in males and females such that the evolution of lifespan in one sex may have unanticipated effects on lifespan in the other.

## Materials and Methods

### Experimental evolution

The seed beetle *A. obtectus* (Coleoptera, Bruchinae) is a widely distributed agricultural pest, infesting *Phaseolus vulgaris* and several other legume crops. They lay their eggs at or near dried beans and the larvae bore into the bean where they complete their development and emerge as adults after 3-4 weeks. A large common base population of *A. obtectus* for all EE lines was established in 1983 by mass mating three local subpopulations collected in the field near Belgrade (41). A detailed description of the EE regimes can be found in Tucić et al. (41, 52). Briefly, the EE lines were initiated in 1986 by placing beetles under two divergent EE regimes (Fig. S40). Under the E life history regime, adult beetles were allowed to mate and lay eggs for the first 48 h of their adult life only.

Under the L regime, beetles could mate but egg laying was permitted only after 10 days of adult life by withholding food resources for offspring (bean seeds) until that point. This thus represents a bi-directional EE design, favoring a short life and early reproduction (E; N = 4 lines) or a long life and late reproduction (L; N = 4 lines), as peak reproduction normally occurs in this species during day 3-6 of adult life (108). Each of the two EE regimes were replicated with four independent lines. Each generation, the E lines were started with approximately 4-500 newly emerged adults placed in rearing jars provided with a surplus of beans. Each L line was started with 1,000 adults kept together in 10 separate vials (approximately 100 individuals per vial) without beans. Beetles that survived to day 10 (∼ 4-500) were then transferred to a common rearing jar provided with beans. The EE lines were maintained in a dark incubator on *P. vulgaris* beans and under adult aphagy at 30°C. Average lifespan depends on environmental conditions, but is approximately 9 and 6 days in E lines and 17 and 9 in L lines (in females and males, respectively) under aphagy (44, 85).

### Methods

We conducted a series of efforts aimed at collectively characterizing the adaptive genetic architecture of the E and L lines (see Fig. S40). We give only a very schematic overview of the methods used here and refer to the Supplementary Materials and Methods for a detailed account. We first assembled a 1.1 Gb chromosome-length genome assembly of *A. obtectus*., using HiC sequencing. We then assessed genome-wide divergent evolution between E and L lines using a Pool-seq approach in generation 329 (E) and 213 (L), where replicated samples were sequenced to ∼100X. Pool-HMM (109, 110) was employed to identify autosomal regions of the genome subjected to selective sweeps. This method involves a likelihood-based scan of the genome for regions where the SNP allele frequency spectrum (AFS) in a sample is biased toward extreme allele frequencies and then implements a Hidden Markov Model for identifying and delineating regions showing signs of sweeps. We used a depth-of-coverage (DOC) strategy to call CNVs, by identifying regions where DOC was consistently significantly different between all E and all L lines. To assess whether evolved genetic differences between EE regimes translate into key phenotypes, we performed a genotype – phenotype association study. Based on a large E/L pedigree, 1334 recombinant F2s were phenotyped (sex, lifespan and body size) and genotyped in 100 kb QTL bins. Interval mapping (IM) and multiple QTL mapping (MQM) models were then used to assess the genotype – phenotype association across the genome. In addition, we asked whether variation in the strength of the genotype – phenotype association across the genome (LOD score) was related to evolutionary differentiation between E and L (sweep and CNV calls). Finally, we conducted a functional Gene Ontology (GO) enrichment analyses of selective sweep and CNV intervals, following standard procedures. To complement these analyses, we implemented a custom multi-evidence Gene Ontology (GO) term analysis to explore whether the functional properties of genes in selective sweeps, CNVs and differentially expressed genes (E versus L) overlap.

### Data, Materials, and Software Availability

The annotated genome assembly is available at ENA (accession GCA_963669975.1). All read data is available at NCBI (BioProject ID PRJNA1540466) and processed data at [Figshare](https://doi.org/10.17044/scilifelab.34059444). Code describing all analyses is available at [GitHub](https://github.com/Goran-Arnqvist/UUBeetleLab/tree/main/Lifespan_selection). An interactive genome browser is publicly available at the [Swedish Reference Genome Portal](https://genomes.scilifelab.se/acanthoscelides_obtectus/).

## Acknowledgements

We wish to thank Johanna Rönn, Gunilla Engström, Yvonne Kallberg and Rebecca Nylund for invaluable help with the experiments and data, and to Locke Rowe for comments on the manuscript. We are also deeply indebted to the late Nikola Tucić for initiating the experimental evolution lines used in this study. This work was funded by the Swedish Research Council (2019-03611 and 2023-03730 to G.A., 2017-02907 to L.A.), the European Research Council (AdG-294333 to G.A.), the Knut and Alice Wallenberg Foundation (KAW 2023.0160 to L.A.), the Ministry of Education, Science and Technological Development of the Republic of Serbia (451-03-136/2025-03/200007) and the Onassis Foundation (to A.T.). Sequencing was performed by the Uppsala University SNP&SEQ Technology Platform as part of the National Genomics Infrastructure at the Science for Life Laboratory, financially supported by the Swedish Research Council and the Knut and Alice Wallenberg Foundation. Bioinformatic work was supported by the National Bioinformatics Infrastructure Sweden, including the Wallenberg Advanced Bioinformatics Infrastructure. The computations were performed on resources provided by the National Academic Infrastructure for Supercomputing in Sweden (NAISS), partially funded by the Swedish Research Council through grant agreement no. 2022-06725, at UPPMAX.

## Author contributions

M.D., U.S. and B.S. managed the long-term experimental evolution; G.A., L.A., M.E.P., A.P.F.-P., C.-J. R., A.T., A.R. and E.I. designed the research; A.S. made the genome assembly; M.K. and Y.F.C prepared samples; A.T., M.E.P., A.P.F.-P., A.R. analyzed data; all authors contributed to the SI; G.A. wrote the first manuscript draft; all authors reviewed and edited the paper.

## References

1. Lemaître, J. F., Moorad, J., Gaillard, J. M., Maklakov, A. A., and Nussey, D. H. (2024) A unified framework for evolutionary genetic and physiological theories of aging. PLoS Biol. 22(2):e3002513.

2. Li, S., Vazquez, J. M., and Sudmant, P. H. (2023) The evolution of aging and lifespan. Trends Genet. 39(11):830–843.

3. Maklakov, A. A., and Chapman, T. (2019) Evolution of ageing as a tangle of trade-offs: Energy versus function. Proc. R. Soc. B 286(1911):20191604.

4. Ivanov, D. K., Escott-Price, V., Ziehm, M., Magwire, M. M., Mackay, T. F. C., Partridge, L., and Thornton, J. M. (2015) Longevity GWAS using the Drosophila genetic reference panel. J. Gerontol. A Biol. Sci. Med. Sci. 70(12):1470–1478.

5. Timmers, P. R., Mounier, N., Lall, K., Fischer, K., Ning, Z., Feng, X., et al. (2019) Genomics of 1 million parent lifespans implicates novel pathways and common diseases and distinguishes survival chances. eLife 8:e39856.

6. Mackay, T. F. C., Roshina, N. V., Leips, J. W., and Pasyukova, E. G. (2005) Complex genetic architecture of Drosophila longevity. In Handbook of the Biology of Aging, pp. 181–216. Academic Press.

7. Pallares, L. F., Lea, A. J., Han, C., Filippova, E. V., Andolfatto, P., and Ayroles, J. F. (2023) Dietary stress remodels the genetic architecture of lifespan variation in outbred Drosophila. Nat. Genet. 55(1):123–129.

8. Chakraborty, M., Emerson, J. J., Macdonald, S. J., and Long, A. D. (2019) Structural variants exhibit widespread allelic heterogeneity and shape variation in complex traits. Nat. Commun. 10(1):4872.

9. Wellenreuther, M., Mérot, C., Berdan, E., and Bernatchez, L. (2019) Going beyond SNPs: The role of structural genomic variants in adaptive evolution and species diversification. Mol. Ecol. 28(6).

10. Dorant, Y., Cayuela, H., Wellband, K., Laporte, M., Rougemont, Q., Mérot, C., et al. (2020) Copy number variants outperform SNPs to reveal genotype–temperature association in a marine species. Mol. Ecol. 29(24):4765–4782.

11. Wilson, J., Bieker, V. C., van Boheemen, L., Connallon, T., Martin, M. D., Battlay, P., and Hodgins, K. A. (2025) Copy number variation contributes to parallel local adaptation in an invasive plant. Proc. Natl. Acad. Sci. U.S.A. 122(10):e2413587122.

12. Butlin, R. and Day, T. (1985) Adult size, longevity and fecundity in the seaweed fly, *Coelopa frigida*. Heredity 54:107–110.

13. Rodriguez, C., Fanara, J. J., and Hasson, E. (1999) Inversion polymorphism, longevity, and body size in a natural population of *Drosophila buzzatii*. Evolution 53(2):612–620.

14. Paris, M., Durmaz Mitchell, E., Kerdaffrec, E., Rubin, D., Spichtig, C., Zurbriggen, F., et al. (2025) Multiple forms of balancing selection maintain inversion polymorphism. Heredity 1–14.

15. Nygaard, M., Debrabant, B., Tan, Q., Deelen, J., Andersen-Ranberg, K., De Craen, A. J. M., et al. (2016) Copy number variation associates with mortality in long-lived individuals: A genome-wide assessment. Aging Cell 15(1):49–55.

16. Auwerx, C., Lepamets, M., Sadler, M. C., Patxot, M., Stojanov, M., Baud, D., et al. (2022) The individual and global impact of copy-number variants on complex human traits. Am. J. Hum. Genet. 109(4):647–668.

17. Bonduriansky, R., Maklakov, A., Zajitschek, F., and Brooks, R. (2008) Sexual selection, sexual conflict and the evolution of ageing and lifespan. Funct. Ecol. 443–453.

18. Maklakov, A. A. and Lummaa, V. (2013) Evolution of sex differences in lifespan and aging: Causes and constraints. BioEssays 35(8):717–724.

19. Lemaître, J. F., Ronget, V., Tidière, M., Allainé, D., Berger, V., Cohas, A., et al. (2020) Sex differences in adult lifespan and aging rates of mortality across wild mammals. Proc. Natl. Acad. Sci. U.S.A. 117(15):8546–8553.

20. Arnqvist, G. and Rowe, L. (2005) Sexual conflict. Princeton University Press.

21. Poissant, J., Wilson, A. J., and Coltman, D. W. (2010) Sex-specific genetic variance and the evolution of sexual dimorphism: A systematic review of cross-sex genetic correlations. Evolution 64(1):97–107.

22. Immonen, E., Hämäläinen, A., Schuett, W., and Tarka, M. (2018) Evolution of sex-specific pace-of-life syndromes: Genetic architecture and physiological mechanisms. Behav. Ecol. Sociobiol. 72:1–23.

23. Bronikowski, A. M., Meisel, R. P., Biga, P. R., Walters, J. R., Mank, J. E., Larschan, E., et al. (2022) Sex-specific aging in animals: Perspective and future directions. Aging Cell 21(2):e13542.

24. Tatar, M. and Carey, J. R. (1994) Sex mortality differentials in the bean beetle: Reframing the question. Am. Nat. 144(1):165–175.

25. Hägg, S. and Jylhävä, J. (2021) Sex differences in biological aging with a focus on human studies. eLife 10:e63425.

26. Bou Sleiman, M., Roy, S., Gao, A. W., Sadler, M. C., von Alvensleben, G. V., Li, H., et al. (2022) Sex- and age-dependent genetics of longevity in a heterogeneous mouse population. Science 377(6614):eabo3191.

27. Vieira, C., Pasyukova, E. G., Zeng, Z. B., Hackett, J. B., Lyman, R. F., and Mackay, T. F. C. (2000) Genotype–environment interaction for quantitative trait loci affecting lifespan in *Drosophila melanogaster*. Genetics 154(1):213–227.

28. Fox, C. W., Bush, M. L., Roff, D. A., and Wallin, W. G. (2004) Evolutionary genetics of lifespan and mortality rates in two populations of the seed beetle, *Callosobruchus maculatus*. Heredity 92(3):170–181.

29. Archer, C. R., Zajitschek, F., Sakaluk, S. K., Royle, N. J., and Hunt, J. (2012) Sexual selection affects the evolution of lifespan and ageing in the decorated cricket *Gryllodes sigillatus*. Evolution 66(10):3088–3100.

30. Berg, E. C. and Maklakov, A. A. (2012) Sexes suffer from suboptimal lifespan because of genetic conflict in a seed beetle. Proc. R. Soc. B 279(1745):4296–4302.

31. Wright, K. M., Rand, K. A., Kermany, A., Noto, K., Curtis, D., Garrigan, D., et al. (2019) A prospective analysis of genetic variants associated with human lifespan. G3 9(9):2863–2878.

32. Leips, J. and Mackay, T. F. C. (2000) Quantitative trait loci for lifespan in *Drosophila melanogaster*: Interactions with genetic background and larval density. Genetics 155(4):1773–1788.

33. Mühlhäuser, C. and Blanckenhorn, W. (2004) The quantitative genetics of sexual selection in the dung fly *Sepsis cynipsea*. Behaviour 141(3):327–341.

34. Lehtovaara, A., Schielzeth, H., Flis, I., and Friberg, U. (2013) Heritability of lifespan is largely sex limited in *Drosophila*. Am. Nat. 182(5):653–665.

35. Rauser, C. L., Mueller, L. D., Travisano, M., and Rose, M. R. (2009) Evolution of aging and late life. In Rose, M. R. and Garland, T. J., eds. Experimental Evolution: Concepts, Methods, and Applications of Selection Experiments, pp. 551–584. University of California Press.

36. Blount, Z. D., Lenski, R. E., and Losos, J. B. (2018) Contingency and determinism in evolution: Replaying life’s tape. Science 362(6415):eaam5979.

37. McHugh, K. M. and Burke, M. K. (2022) From microbes to mammals: The experimental evolution of aging and longevity across species. Evolution 76(4):692–707.

38. Remolina, S. C., Chang, P. L., Leips, J., Nuzhdin, S. V., and Hughes, K. A. (2012) Genomic basis of aging and life-history evolution in *Drosophila melanogaster*. Evolution 66:3390–3403.

39. Carnes, M. U., Campbell, T., Huang, W., Butler, D. G., Carbone, M. A., Duncan, L. H., et al. (2015) The genomic basis of postponed senescence in *Drosophila melanogaster*. PLoS One 10:e0138569.

40. Fabian, D. K., Garschall, K., Klepsatel, P., Santos-Matos, G., Sucena, É., Kapun, M., et al. (2018) Evolution of longevity improves immunity in *Drosophila*. Evol. Lett. 2:567–579.

41. Tucić, N., Gliksman, I., Šešlija, D., Milanović, D., Mikuljanac, S., and Stojković, O. (1996) Laboratory evolution of longevity in the bean weevil (*Acanthoscelides obtectus*). J. Evol. Biol. 9(4):485–503.

42. Promislow, D. E., Flatt, T., and Bonduriansky, R. (2022) The biology of aging in insects: From *Drosophila* to other insects and back. Annu. Rev. Entomol. 67(1):83–103.

43. Tucić, N., Gliksman, I., Šešlija, D., Stojković, O., and Milanović, D. (1998) Laboratory evolution of life-history traits in the bean weevil (*Acanthoscelides obtectus*): the effects of selection on developmental time in populations with different previous history. Evolution 52(6):1713–1725.

44. Đorđević, M., Stojković, B., Savković, U., Immonen, E., Tucić, N., Lazarević, J., and Arnqvist, G. (2017) Sex-specific mitonuclear epistasis and the evolution of mitochondrial bioenergetics, ageing, and life history in seed beetles. Evolution 71(2):274–288.

45. Đorđević, M., Savković, U., Lazarević, J., Tucić, N., and Stojković, B. (2015) Intergenomic interactions in hybrids between short-lived and long-lived lines of a seed beetle: analyses of life history traits. Evol. Biol. 42(4):461–472.

46. Arnqvist, G., Stojković, B., Rönn, J. L., and Immonen, E. (2017) The pace-of-life: A sex-specific link between metabolic rate and life history in bean beetles. Funct. Ecol. 31(12):2299–2309.

47. Lazarević, J., Tucić, N., Šešlija Jovanović, D., Večeřa, J., and Kodrík, D. (2012) The effects of selection for early and late reproduction on metabolite pools in *Acanthoscelides obtectus* Say. Insect Sci. 19(3):303–314.

48. Lazarević, J., Đorđević, M., Stojković, B., and Tucić, N. (2013) Resistance to prooxidant agent paraquat in the short- and long-lived lines of the seed beetle (*Acanthoscelides obtectus*). Biogerontology 14(2):141–152.

49. Immonen, E., Sayadi, A., Stojković, B., Savković, U., Đorđević, M., Liljestrand-Rönn, J., et al. (2023) Experimental life history evolution results in sex-specific evolution of gene expression in seed beetles. Genome Biol. Evol. 15(1):evac177.

50. Šešlija, D., Lazarević, J., Janković, B., and Tucić, N. (2009) Mating behavior in the seed beetle *Acanthoscelides obtectus* selected for early and late reproduction. Behav. Ecol. 20(3):547–552.

51. Fromonteil, S., Rêgo, A., Immonen, E., Stojković, B., Savković, U., Đorđević, M., et al. (2026) The effects of sexual selection on functional and molecular reproductive divergence during experimental evolution in seed beetles. Evol. Lett. 10(1):91–103.

52. Tucic, N., Stojkovic, O., Gliksman, I., Milanovic, D., and Seslija, D. (1997) Laboratory evolution of life-history traits in the bean weevil (*Acanthoscelides obtectus*): The effects of density-dependent and age-specific selection. Evolution 51(6):1896–1910.

53. Marieshwari, B. N., Bhuvaragavan, S., Sruthi, K., Mullainadhan, P., and Janarthanan, S. (2023) Insect phenoloxidase and its diverse roles: Melanogenesis and beyond. J. Comp. Physiol. B 193(1):1–23.

54. Yiu, T. C., Tu, J., and Cheung, H. H. (2025) An overview of RecQ helicases and related diseases. Aging (Albany NY) 17(7):1881.

55. Cassidy, D., Epiney, D. G., Salameh, C., Zhou, L. T., Salomon, R. N., Schirmer, A. E., et al. (2019) Evidence for premature aging in a *Drosophila* model of Werner syndrome. Exp. Gerontol. 127:110733.

56. Anderson, R. M. and Sinclair, D. A. (2004) Yeast RecQ helicases: Clues to DNA repair, genome stability and aging. Mol. Mech. Werner’s Syndr. 78.

57. Hamazaki, J. and Murata, S. (2024) Relationships between protein degradation, cellular senescence, and organismal aging. J. Biochem. 175(5):473–480.

58. Noh, M. Y., Kramer, K. J., Muthukrishnan, S., Beeman, R. W., Kanost, M. R., and Arakane, Y. (2015) Loss of function of the *yellow-e* gene causes dehydration-induced mortality of adult *Tribolium castaneum*. Dev. Biol. 399(2):315–324.

59. Clancy, D. J., Gems, D., Harshman, L. G., Oldham, S., Stocker, H., Hafen, E., et al. (2001) Extension of life-span by loss of CHICO, a *Drosophila* insulin receptor substrate protein. Science 292(5514):104–106.

60. Mathew, R., Pal Bhadra, M., and Bhadra, U. (2017) Insulin/insulin-like growth factor-1 signalling (IIS)-based regulation of lifespan across species. Biogerontology 18(1):35–53.

61. Broman, K. W., Gatti, D. M., Simecek, P., Furlotte, N. A., Prins, P., Sen, Ś., Yandell, B. S., and Churchill, G. A. (2019) R/qtl2: Software for mapping quantitative trait loci with high-dimensional data and multi-parent populations. Genetics 211:495–502.

62. Stavrovskaya, E. D., Niranjan, T., Fertig, E. J., Wheelan, S. J., Favorov, A. V., and Mironov, A. A. (2017) StereoGene: Rapid estimation of genome-wide correlation of continuous or interval feature data. Bioinformatics 33(20):3158–3165.

63. Demuth, J. P. and Wade, M. J. (2006) Experimental methods for measuring gene interactions. Annu. Rev. Ecol. Evol. Syst. 37(1):289–316.

64. Šeslija, D. and Tucic, N. (2013) The genetic architecture of extended life span in the seed beetle *Acanthoscelides obtectus* (Coleoptera: Bruchidae). Eur. J. Entomol. 105(4):553–560.

65. Watanabe, K., Stringer, S., Frei, O., Umićević Mirkov, M., de Leeuw, C., Polderman, T. J. C., et al. (2019) A global overview of pleiotropy and genetic architecture in complex traits. Nat. Genet. 51(9):1339–1348.

66. Pettersson, M., Besnier, F., Siegel, P. B., and Carlborg, Ö. (2011) Replication and explorations of high-order epistasis using a large advanced intercross line pedigree. PLoS Genet. 7(7):e1002180.

67. Mackay, T. F. C. and Anholt, R. R. H. (2024) Pleiotropy, epistasis and the genetic architecture of quantitative traits. Nat. Rev. Genet. 25(9):639–657.

68. Lauer, S. and Gresham, D. (2019) An evolving view of copy number variants. Curr. Genet. 65(6):1287–1295.

69. Tomanek, I. and Guet, C. C. (2022) Adaptation dynamics between copy-number and point mutations. eLife 11:e82240.

70. Farslow, J. C., Lipinski, K. J., Packard, L. B., Edgley, M. L., Taylor, J., Flibotte, S., et al. (2015) Rapid increase in frequency of gene copy-number variants during experimental evolution in *Caenorhabditis elegans*. BMC Genomics 16(1):1044.

71. Gamazon, E. R. and Stranger, B. E. (2015) The impact of human copy number variation on gene expression. Brief. Funct. Genomics 14(5):352–357.

72. Stranger, B. E., Forrest, M. S., Dunning, M., Ingle, C. E., Beazley, C., Thorne, N., et al. (2007) Relative impact of nucleotide and copy number variation on gene expression phenotypes. Science 315(5813):848–853.

73. Springer, M., Weissman, J. S., and Kirschner, M. W. (2010) A general lack of compensation for gene dosage in yeast. Mol. Syst. Biol. 6(1):368.

74. Ma, C., Sangdehi, F. M., Kawaguchi, M., Sano, K., Dannenberg, S. V., Pettersson, M. E., Wallberg, A., Wort, J. L., Yan, Y., Moshkovskii, S., Berg, F., Folkvord, A., Lenz, C., Urlaub, H., Kaupp, U. B., Yasumasu, S., and Andersson, L. (2026) Sperm, egg and embryo proteins critical for genetic adaptation of herring to low salinity in the Baltic Sea. Proc. Natl. Acad. Sci. U.S.A., in press.

75. Lappalainen, T. and MacArthur, D. G. (2021) From variant to function in human disease genetics. Science 373(6562):1464–1468.

76. Quenez, O., Cassinari, K., Coutant, S., Lecoquierre, F., Le Guennec, K., Rousseau, S., et al. (2021) Detection of copy-number variations from NGS data using read depth information: A diagnostic performance evaluation. Eur. J. Hum. Genet. 29(1):99–109.

77. Lei, Y., Meng, Y., Guo, X., Ning, K., Bian, Y., Li, L., et al. (2022) Overview of structural variation calling: Simulation, identification, and visualization. Comput. Biol. Med. 145:105534.

78. Schrider, D. R., Begun, D. J., and Hahn, M. W. (2013) Detecting highly differentiated copy-number variants from pooled population sequencing. In Pac. Symp. Biocomput., p. 344.

79. Hamilton, W. D. (1966) The moulding of senescence by natural selection. J. Theor. Biol. 12:12–45.

80. Williams, G. C. (1957) Pleiotropy, natural selection and the evolution of senescence. Evolution 11:398–411.

81. Kirkwood, T. B. L. (1977) Evolution of aging. Nature 270:301–304.

82. Medawar, P. B. (1952) An Unsolved Problem of Biology. Lewis, London.

83. Moses, E., Bergman, M., Atlan, T., Duxbury, E. M., Franěk, R., Ben Dor, O., et al. (2026) An antagonistically pleiotropic gene regulates vertebrate growth, maturity, and lifespan. Nat. Commun. 17(1):4410.

84. Weber, A. L., Khan, G. F., Magwire, M. M., Tabor, C. L., Mackay, T. F. C., and Anholt, R. R. H. (2012) Genome-wide association analysis of oxidative stress resistance in *Drosophila melanogaster*. PLoS One 7(4):e34745.

85. Stojković, B. and Savković, U. (2011) Gender differences in longevity in early and late reproduced lines of the seed beetle. Arch. Biol. Sci. 63(1):129–136.

86. Tatar, M., Post, S., and Yu, K. (2014) Nutrient control of *Drosophila* longevity. Trends Endocrinol. Metab. 25(10):509–517.

87. Chippindale, A. K., Hoang, D. T., Service, P. M., and Rose, M. R. (1994) The evolution of development in *Drosophila melanogaster* selected for postponed senescence. Evolution 48(6):1880–1899.

88. Khazaeli, A. A., Van Voorhies, W., and Curtsinger, J. W. (2005) The relationship between lifespan and adult body size is highly strain-specific in *Drosophila melanogaster*. Exp. Gerontol. 40(5):377–385.

89. Barghi, N., Hermisson, J., and Schlötterer, C. (2020) Polygenic adaptation: A unifying framework to understand positive selection. Nat. Rev. Genet. 21(12):769–781.

90. Láruson, Á. J., Yeaman, S., and Lotterhos, K. E. (2020) The importance of genetic redundancy in evolution. Trends Ecol. Evol. 35(9):809–822.

91. Connallon, T. and Matthews, G. (2019) Cross-sex genetic correlations for fitness and fitness components: Connecting theoretical predictions to empirical patterns. Evol. Lett. 3(3):254–262.

92. Nuzhdin, E. G., Pasyukova, E. G., Dilda, C. L., Zeng, Z., and Mackay, T. F. C. (1997) Sex-specific quantitative trait loci affecting longevity in *Drosophila melanogaster*. Proc. Natl. Acad. Sci. U.S.A. 94(18):9734–9739.

93. Leips, J. and Mackay, T. F. C. (2002) The complex genetic architecture of *Drosophila* lifespan. Exp. Aging Res. 28(4):361–390.

94. Wilson, R. H., Morgan, T. J., and Mackay, T. F. C. (2006) High-resolution mapping of quantitative trait loci affecting increased lifespan in *Drosophila melanogaster*. Genetics 173(3):1455–1463.

95. Yeaman, S. (2013) Genomic rearrangements and the evolution of clusters of locally adaptive loci. Proc. Natl. Acad. Sci. U.S.A. 110(19):E1743–E1751.

96. Dijkers, P. F. and O’Farrell, P. H. (2007) *Drosophila* calcineurin promotes induction of innate immune responses. Curr. Biol. 17(23):2087–2093.

97. Bagchi, B., Corbel, Q., Khan, I., Payne, E., Banerji, D., Rönn, J. L., et al. (2021) Sexual conflict drives micro- and macroevolution of sexual dimorphism in immunity. BMC Biol. 19(1):1–19.

98. Kaneuchi, T., Sartain, C. V., Takeo, S., Horner, V. L., Buehner, N. A., Aigaki, T., and Wolfner, M. F. (2015) Calcium waves occur as *Drosophila* oocytes activate. Proc. Natl. Acad. Sci. U.S.A. 112(3):791–796.

99. Adler, M. I. and Bonduriansky, R. (2014) Sexual conflict, lifespan, and aging. Cold Spring Harb. Perspect. Biol. 6(8):a017566.

100. Xia, S., Arsala, D., Gschwend, A., Koval, W., Atlas, J., Han, S., et al. (2026) Analysis of the impact of gene evolution on reproductive effects reveals prevalent sexual and germline–soma conflicts. Nat. Ecol. Evol. 1–13.

101. Maklakov, A. A., Bonduriansky, R., and Brooks, R. C. (2009) Sex differences, sexual selection, and ageing: An experimental evolution approach. Evolution 63(10):2491–2503.

102. Harano, T., Yasui, Y., and Miyatake, T. (2006) Direct effects of polyandry on female fitness in *Callosobruchus chinensis*. Anim. Behav. 71(3):539–548.

103. Ricklefs, R. E. and Cadena, C. D. (2008) Heritability of longevity in captive populations of nondomesticated mammals and birds. J. Gerontol. A Biol. Sci. Med. Sci. 63(5):435–446.

104. Ruzicka, F., Zwoinska, M. K., Goedert, D., Kokko, H., Li Richter, X. Y., Moodie, I. R., et al. (2026) A century of theories of balancing selection. Biol. Rev. 101(2):804–825.

105. Zajitschek, F. and Connallon, T. (2018) Antagonistic pleiotropy in species with separate sexes, and the maintenance of genetic variation in life-history traits and fitness. Evolution 72(6):1306–1316.

106. Berger, D., Berg, E. C., Widegren, W., Arnqvist, G., and Maklakov, A. A. (2014) Multivariate intralocus sexual conflict in seed beetles. Evolution 68(12):3457–3469.

107. Berger, D., Martinossi-Allibert, I., Grieshop, K., Lind, M. I., Maklakov, A. A., and Arnqvist, G. (2016) Intralocus sexual conflict and the tragedy of the commons in seed beetles. Am. Nat. 188(4):E98–E112.

108. Parsons, D. M. and Credland, P. F. (2003) Determinants of oviposition in *Acanthoscelides obtectus*: A nonconformist bruchid. Physiol. Entomol. 28(3):221–231.

109. Boitard, S., Schlötterer, C., Nolte, V., Pandey, R. V., and Futschik, A. (2012) Detecting selective sweeps from pooled next-generation sequencing samples. Mol. Biol. Evol. 29:2177–2186.

110. Boitard, S., Kofler, R., Françoise, P., Robelin, D., Schlötterer, C., and Futschik, A. (2013) Pool-hmm: A Python program for estimating the allele frequency spectrum and detecting selective sweeps from next generation sequencing of pooled samples. Mol. Ecol. Resour. 13(2):337–340.

