## Supplemental Information for "Experimental evolution unveils a role for structural variants and sex-specificity in the genetic architecture of lifespan"

#### Supplementary Information

**Content:** **Page:**

| Content: | Page: |
|---|---|
| 1. Supplementary Materials and Methods | 2 |
| 2. Supplementary Results | 21 |
| 3. Supplementary References | 23 |

### 1. Supplementary Materials and Methods

**HiC library preparation, sequencing, scaffolding and annotation**

As our point of venture, we used an extant, well annotated, high quality genome assembly of *A. obtectus*, based on 80× long-read (PacBio) sequence data (Immonen et al. 2023) (GenBank accession GCA_933228535.1). While near complete (1.1 Gb), this assembly is fragmented and we thus first super-scaffolded the assembly, using a dedicated HiC sequencing effort, into a contiguous assembly containing 10 chromosome-length scaffolds (9A + X) that forms the reference for our current work (GenBank accession GCA_963669975.1).

We used a sample of a live male *A. obtectus* from an isogenic strain derived from the base population of our E and L lines for HiC sequencing. The beetle was killed by flash freezing in liquid nitrogen and was then ground into a flour-like powder using a plastic pestle in an Eppendorf tube on dry ice. The pestle and tube used were pre-cooled in liquid nitrogen prior to grounding and the tube contained some liquid nitrogen, to ensure that the material remained frozen through the entire grinding, preparation and transfer process. The sample was then stored at -80˚C.

The HiC library was prepared following the Arima Protocol "Arima-HiC_AnimalTissue_v00" (Document number A160126 v00). In short, 400 mg of ground beetle material was crosslinked in 2 % Formaldehyde for 20 min while rotating. The crosslinking was stopped by addition of the Arima-kit provided Stop Solution. Before continuation, larger debris was allowed to sink to the bottom and only material small enough to pipette was used in subsequent reactions. Lysis was performed on crosslinked tissue equivalent to approximately 2000 ng of chromatin. All steps of the HiC reaction was performed as described in the Arima protocol.

The chromatin was fragmented in an AFA Fiber Crimp-cap microTUBE using a COVARIS E220 with the following settings: Peak Incident Power 175 W, Acoustic Duty Factor 10 %, 200 Cycles Per Burst, 50 s treatment time. The fragmented chromatin was purified using AMPure XP beads and subjected to library preparation following the "Arima-HiC Kit, Library Preparation using Illumina TruSeq DNA PCR-Free Library Prep" Protocol (Document number A160111 v01) and the Arima-kit reagents were supplemented by reagents of an Illumina TruSeq PCR-Free library preparation kit. The library was amplified for 11 cycles using an Illumina TruSeq DNA CD Index. The final library was analysed for fragment length distribution using an Agilent Fragment Analyser with a high sensitivity NGS Fragment 1-6000bp kit and for concentration using the Qubit high sensitivity dsDNA kit.

Subsequently, the library was sequenced with a depth of approximately 690 million reads on a NovaSeq 6000 S4 flow cell with a read length of 2x150 bp.

The HiC reads were pre-processed using Juicer (git-rev. 84f6957) (Durand et al. 2016) and a script provided by Arima Genomics (https://github.com/ArimaGenomics/Scripts) to generate ligation site positions in the input assembly (GCA_933228535.1) using matching the sequence motifs 'GATCGATC', 'GANTGATC', 'GANTANTC' and 'GATCANTC'. The resulting list of valid HiC pairs was used as input for the scaffolder 3D-DNA (v. 180922). The draft scaffolded assembly was then manually error corrected and curated using the using the JBAT method provided by the authors of 3D-DNA (Dudchenko et al. 2017).

The resulting Hi-C genome assembly contained some 90% of all scaffolds within 10 chromosome-length scaffolds (Fig. S1). The new assembly was annotated by performing a coordinate conversion of the extant annotation (ENA project ID: PRJEB51445). We performed the liftover using Liftoff (Shumate and Salzberg 2021), using default parameters apart from a few that were set to more stringent criteria (coverage >80%, sequence identity >90%, distance scaling 5). The annotated genome assembly (Table S1) has been deposited and is available at ENA (accession GCA_963669975.1).

**Pool sequencing and differentiation of EE lines**

To assess genome-wide divergent evolution between lines, we sequenced pools of individuals (N=50 in each sample/pool) from all E and L lines (two biological replicate samples per population) from generation 329 (E) and 213 (L). Each sample was sequenced in four lanes (Illumina HiSeq 2500), resulting in ~50X coverage per sample (i.e., ~100X per line). Following assessment of read quality, reads were mapped to the reference genome and alignment statistics and quality metrics were evaluated. Variants were called with GATK v3.8 UnifiedGenotyper using default settings, and biallelic SNPs were extracted and filtered through successive quality control steps to retain high-confidence variants. Based on these calls, we then estimated pair-wise genome-wide $F_{ST}$ between samples (Hivert et al. 2018) and scanned the genome for differentiation (delta allele frequency; dAF) between super-pools of all E and L lines.

We first extracted high-quality DNA samples from pools of individuals, using a salt-ethanol precipitation protocol. Beetles were first gently macerated and placed in preparation buffer (100 mM NaCl, 10 mM Tris-HCl, pH = 8.0, 0.5% SDS) together with proteinase K, vortexed and incubated at 50°C overnight. Samples were then frozen overnight. To precipitate DNA, we added saturated NaCl several times before adding 95% ethanol, and then spun the DNA into a pellet. The DNA pellet was suspended in TE buffer (pH = 7.6). DNA quality and quantity was assessed using NanoDrop, Qubit and Bioanalyzer, followed by fragment length assessment on an agarose gel.

We prepared two independent samples from each of the 8 lines ($N_{tot}$ = 16 samples), each sample consisting of a pool of either N = 50 males or N = 50 females. Sequencing libraries were prepared from 1µg DNA using the TruSeq PCRfree DNA sample preparation kit (cat# FC-121-3001/3002, Illumina Inc.) targeting an insert size of 350bp. The library preparation was performed according to the manufacturers' instructions (guide#15036187). Libraries were then subjected to cluster generation and sequencing in 8 lanes using the Illumina HiSeq2500 system, paired-end 125bp read length and v4 sequencing chemistry. We sequenced on average some 300 million read pairs for each lane, resulting in an average coverage per sample of about 50X and an average cover per line of about 100X.

Read quality was assessed using FastQC v0.11.8 (Andrews et al. 2022) and aggregated FastQC reports were generated with MultiQC v1.9 (Ewels et al. 2016). Reads were mapped to the reference genome using BWA-MEM2 v2.2.1 (Vasimuddin et al. 2019) and post-processing steps, including duplicate marking and mate pair correction, were carried out using SAMtools v1.21 (Li H et al. 2009, Danecek et al. 2021). Alignment statistics and quality metrics were subsequently evaluated using Qualimap v2.3 (García-Alcalde et al. 2012, Okonechnikov et al. 2016).

Variant calling was carried out using GATK v3.8 (McKenna et al., 2010) with the UnifiedGenotyper default parameters. From the initial call set, we extracted biallelic SNPs and applied successive quality control filters to retain high confidence variants. First, we used hard filtering based on genome-wide distributions of GATK quality metrics, excluding SNPs with FS > 60.0, SOR > 3.0, MQ < 40.0, MQRankSum < −12.5, or ReadPosRankSum < −8.0 (details on these annotations are provided by the Broad Institute, 2019). We then kept SNPs with genotype quality greater than 20 (GQ > 20), permitted up to 20% missing data per locus, required a minor allele count (MAC) of at least 3 reads, and removed monomorphic sites using BCFtools v1.10 (Li et al., 2009). Finally, to reduce false positives associated with copy-number variants and repetitive regions that tend to exhibit unusually high coverage, we applied an additional depth-of-coverage filter. In R (R Core Development Team, 2021), we generated a depth of coverage distribution for each pool using per-SNP read depth (DP). Three alternative thresholds were assessed, ranging from most to least stringent: mean ± 1 standard deviation, mode ± half the mode, and the 5th–99th percentile range of coverage per sample (Fig. S2). For each approach, only SNPs that satisfied the coverage criteria across all pools were retained. We selected the 5–99% quantile filter because it preserved many loci while removing sites with exceptionally low or high depth. The final set of high-confidence SNPs was used for downstream analyses.

**Population genetic structure**

We evaluated population structure among the bean beetles' pools using pairwise $F_{ST}$ estimates. Pairwise pool-$F_{ST}$ ($\hat{F}_{\mathrm{ST}}^{\mathrm{pool}}$) values were calculated for all pool comparisons from raw per-SNP read counts with the R package poolfstat (Hivert et al. 2018). This estimator is comparable to the Weir and Cockerham (1984) $F_{ST}$ and explicitly accounts for the additional sampling variance introduced by pool-seq through random chromosome sampling. pool-$F_{ST}$ ranges from 0 to 1, with 0 indicating no genetic differentiation and 1 indicating complete differentiation among populations. We generated a heatmap plot of the pairwise pool-$F_{ST}$ values using ggplot2 (Wickham, 2016).

**Outlier genomic regions putatively under selection**

Prior to estimating allele frequencies for each pool, we converted raw read counts to *effective coverage* per SNP ($n_{\mathrm{eff}}$), which approximates the number of sampled chromosomes while accounting for sequencing depth. This adjustment corrects for stochastic differences in read coverage and chromosome sampling among pools during sequencing (Bergland et al., 2014; Feder et al., 2012; Kolaczkowski et al., 2011). We implemented the correction in Python using $n_{\mathrm{eff}} = \frac{(n \cdot RD)-1}{n+RD}$, where $RD$ is read depth and $n$ is the number of chromosomes in the pool (equal to $2N$ for a diploid species, with $N$ individuals per pool). Population allele frequencies were subsequently calculated from the $n_{\mathrm{eff}}$-adjusted read counts using a Python script.

To detect genomic regions showing higher differentiation than the genome-wide background, treated as putative targets of selection, we computed, for each SNP, the absolute allele-frequency difference (dAF) between paired contrasts of the superpools of the E and L lines (a 'superpool' referes to the data aggregation of all the pools of the E or the L line): *dAF = | meanAF(group1) − meanAF(group2) |*. To further examine broader signals supported by multiple neighboring markers and to reduce the influence of isolated SNPs potentially driven by Pool-seq sampling noise, we also calculated a rolling mean of dAF across windows of 100, 500, 1000, 5000, and 10 000 SNPs. All analyses were conducted in R, and figures were generated using ggplot2.

**Selective sweeps**

Briefly, following masking ("blacklisting") of potentially problematic regions, showing low mapping quality, abnormal coverage or being located close to assembly gaps, we employed Pool-HMM (Boitard et al. 2012, 2013) to identify autosomal regions of the genome subjected to selective sweeps. This method involves a likelihood-based scan of the genome for regions where the SNP allele frequency spectrum (AFS) in a sample is biased toward extreme allele frequencies and then implements a Hidden Markov Model for identifying and delineating regions showing signs of sweeps.

Read pileups were first generated for each Pool-seq sample using SAMtools v0.1.19. Evolutionary signatures were subsequently inferred for assembled scaffolds with Pool-HMM v1.4.4 (Boitard et al. 2012, 2013), using an initial theta value of 0.005, a chromosome count per sample set to 100, and a mutation rate parameter k=10^(-7). Chromosome X, even considering modified parameters for chromosomal numbers in male samples, exhibited minimal selective signatures, and hence was excluded. To reduce computational demands, only one in every ten allelic positions was analyzed. The workflow was managed using Nextflow v22.10.1 (Di Tommaso et al. 2017) and fully containerized with Apptainer (Kurtzer et al. 2021).

To assess similarity in the genome-wide signal of selective sweeps across replicate lines, selective sweep intervals identified by Pool-HMM were exported as BED files for all eight lines and analyzed at the interval level based on presence or absence of a sweep call (1 or 0). Pairwise overlap between EE lines was quantified using BEDTools v2.31 (Quinlan & Hall 2010, Dale et al. 2011) as the Jaccard distance matrix across replicate samples. This matrix was then used to ordinate all samples, using Principal Coordinates Analysis (PCoA) in scikit-bio v0.6.3 (Rideout et al. 2024), and EE centroids were compared between E and L regimes using PERMANOVA as implemented in the adonis2 function in vegan v2.7.0 (Oksanen et al. 2022) (5,000 permutations). In addition to these overlap-based analyses, we computed whole-genome empirical cumulative distribution function (ECDF) distance correlations of sweep status across all sample pairs using GenometriCorr v1.1.24 (Favorov et al. 2012), providing an alternative similarity matrix that accounts for genome-wide random association expectations.

**CNV calling**

Briefly, we used a depth-of-coverage (DOC) strategy to call CNVs (Lei *et al.* 2022), which is effective at capturing dosage changes spanning distant or repetitive genomic regions, such as transposable elements or large tandem arrays (Schrider *et al.* 2013). Given the high repeat content of the *A.*

*obtectus* genome (>60%), we considered a DOC-based method appropriate. We first assessed whether E and L lines showed consistent differences in DOC by subjecting data on average read depth per non-overlapping 100 bp bins in all samples to a factor analysis (Unsupervised Non-Negative Matrix Factorization). These analyses showed a very clear separation of E and L lines. Following a series of steps to remove noise in DOC data (excluding blacklisted regions and regions with aberrant mappability or GC content – see below), we derived a per-bin signal of DOC divergence based on the ratio of median DOC in L lines to that in E lines. This signal was de-noised using a stationary wavelet transform and CNVs were then called by a Circular Binary Segmentation (Olshen et al. 2011). To validate calls, we tested whether L and E lines differed in the summed per-bin median DOC values with a permutation test. Regions not significantly deviating (fold-change <1.5, P > 0.05) were removed. We note that our procedure is conservative in that it only calls CNVs where the configuration is consistently different between replicate L and E lines.

Prior to CNV calling, we evaluated whether genomic regions exhibited meaningful differences DOC between samples from E and L lines. Average read depth per 100 bp bins was calculated for all pool-seq WGS samples, with normalization to the sample mean performed separately for autosomes and sex chromosomes. Unsupervised Non-Negative Matrix Factorization (NMF) was conducted using scikit-learn v1.6.1 (Pedregosa *et al.* 2011), specifying four components based on a scree plot of reconstruction error, and employing the Kullback–Leibler beta divergence as the loss function. Assuming that only a limited number of loci contribute to the observed signal, sparsity was imposed on the H matrix through combined L1 and L2 regularization ($\alpha_H$ = 0.1, $L1_{ratio}$ = 0.5), promoting attribution of variance to localized genomic bins. This showed striking differences between E and L lines (see below).

Because *A. obtectus* is a non-model organism, the genome lacks a curated list of problematic genomic regions. Existing tools for *de novo* blacklist creation, such as the ENCODE Blacklist (Amemiya *et al.* 2019), are primarily designed for functional genomic datasets and did not perform well with our data. We used the following procedure to identify regions with aberrant characteristics, including abnormal coverage, high proportions of low-quality reads, proximity to assembly gaps, and unassembled contigs.

For coverage-based filtering, a pipeline analogous to the ENCODE Blacklist approach was implemented. Coverage was calculated using mosdepth v0.3.3 (Pedersen & Quinlan 2018) in 1000 bp windows with a 100 bp step size (i.e. 900 bp overlap). Regions falling below the 7th percentile or above the 99th percentile of coverage in each sample were flagged. A consensus list of anomalous regions was constructed using BEDtools multiinter v2.31.1 (Quinlan & Hall 2010, Dale *et al.* 2011). A region was retained if at least 3 out of 8 samples agreed on its classification. To account for potential regime-specific effects, only regions identified as anomalous in both EE regimes were included in the final blacklist. When appropriate, GNU Parallel (Tange 2023) was used to parallelize the workflow.

A similar procedure was used to identify regions with low mapping quality. In this case, the metric was the read count drop-off fraction, defined as the proportion of reads with MAPQ ≥10 falling below a threshold of 0.4. Additionally, regions located within 150 bp of assembly gaps and contigs smaller than 20 kb were excluded.

To further support downstream analyses, a custom mappability map was also generated. Simulated paired-end reads were produced from the reference genome using dicey chop v0.3.3 (Rausch *et al.* 2020, 2025a) with an insert size of 400 bp, reflecting the empirical library characteristics. These reads were then mapped back to the reference using BWA MEM2 v2.2.1 (Vasimuddin *et al.* 2019). Mean coverage was computed for non-overlapping bins of various sizes using mosdepth, and regional mappability was defined as the ratio of bin-level coverage to the maximum genome-wide depth, producing values normalized to the [0,1] interval. To ensure compatibility with downstream analyses, read filtering was similarly applied during mappability estimation.

Initially, we applied CNVpytor v1.3.1 (an updated implementation of CNVnator) (Abyzov *et al.* 2011, Suvakov *et al.* 2021) on a per-sample basis. However, its performance on our dataset was suboptimal, as evidenced by manual inspection of per-bin coverage signals, reflecting regions affected by GC-content and mappability biases that distorted read depth signals (Benjamini & Speed 2012a). Other tools either require a reference sample, as in the case of matched tumor / normal tissue samples (Rausch *et al.* 2012, Fromer & Purcell 2014, Babadi *et al.* 2023), or rely on ploidy assumptions that are violated in Pool-seq data (Boeva *et al.* 2011, Boeva *et al.* 2012, Klambauer *et al.* 2012, Talevich *et al.* 2016, Karunarathne *et al.* 2023). Given the specific challenges associated with Pool-seq data, conventional CNV calling approaches must thus be adapted. In our case, the experimental design offers an opportunity to use the two EE regimes as mutual controls. We chose to use this strategy in order to focus on detecting consistent evolved differences in DOC per region between E and L lines. A similar pipeline was employed by Schrider et al. (2013).

Coverage for non-overlapping bins of 100bp was calculated for each sample with mosdepth, filtering any reads with MAPQ < 20, and with the flag 3852. Bins with one or more of the following characteristics were removed: (i) present in the aberrant coverage blacklist described above, (ii) GC content in the 5$^{th}$ or 99$^{th}$ percentile or (iii) mappability < 0.10. Bin-level read depths were then GC-normalized by fitting a one degree LOESS regression of mean depth against rounded GC content (1% bins), weighted by the number of windows per GC class using scikit-misc v0.1.4. Corrected values were obtained by scaling observed depths to the genome-wide mean using the predicted GC bias (Benjamini & Speed 2012b). After merging and median normalization of biological replicates, the same approach was undertaken for mappability bias. The correlation between populations was calculated with NumPy. Finally, to satisfy the control aspect of the pipeline, the matrix of observed DOC values was transformed into a log-of-ratios 1D signal as

$$signal_i = \log_2\left(\frac{\mathrm{median}(\boldsymbol{L}_{*,i})}{\mathrm{median}(\boldsymbol{E}_{*,i})}\right)$$

where ***L*** and ***E*** are coordinate-ordered coverage matrices for EE regimes (rows: lines, columns: genomic bins), and *i* indexes the genomic bin. The $\log_2$-ratio was computed from the median values across lines, to avoid impact of outliers.

While the preceding steps controlled for several sources of bias, we found that the signal still contained residual noise, as expected due to the complexity of DOC data. As is customary, we thus employed a final denoising step prior to CNV calling. Simple moving average denoising can result in ”smearing” of spike edges leading to suboptimal performance. We thus chose to implement a stationary wavelet transform (MODWT) (Percival & Walden 2000) with a Haar basis to the log2-ratio signal, at five levels. Noise thresholds for each detail level were determined using a SureShrink dynamic threshold (Stein 1981, Donoho & Johnstone 1995) and applied via soft-thresholding to avoid artifacts.

To call CNVs, segmentation of denoised log-ratio data was performed using a Circular Binary Segmentation algorithm via the PSCBS R package v0.68.0 (Olshen *et al.* 2011), with parameters aligned to the CNVkit pipeline except for an undo threshold of 4 SDs to reduce over-segmentation. Segments with length less than 1kb or with more than 60% prevalence of nulls were filtered. Finally, to assess whether a segment truly differs between the two EE regimes, a permutation test was performed. In short, the labels for the original normalized population-level data were permuted, with the utilized test metric being the difference of means of the summed per bin median coverage values. This was used to mimic the data processing in previous steps. We note that since there are 8 replicate lines, the minimum two-sided P-value is 0.029 and multiple testing correction is thus not possible. Segments not significantly deviating between EE regimes (fold-change <1.5, P-value > 0.05) were removed from further processing. CNV state (gain or loss) was called for the median of each population in relation to the segments of the genome identified as "normal" by the contrasting EE regime.

**Correlation between CNVs and selective sweep signatures**

Spatial associations between CNV calls and selective sweeps calls were quantified using GenometriCorr v1.1.24 (Favorov et al. 2012). CNV tracks were defined separately for deletions and gains identified in the Early and Late regimes, and selective sweep signatures were represented as BED intervals corresponding to consistently selected regions within each regime, as defined above. Prior to analysis, all tracks were harmonized by merging adjacent intervals separated by ≤10 kb to minimize the effects of fragmentation on relative midpoint distance estimates. Chromosome X was excluded from all analyses to preserve a valid null distribution for permutation testing. For each CNV track, GenometriCorrelation was run against each consistently selected interval track using whole-genome summarization (awhole.only=TRUE). Genomic interval association was quantified using the empirical cumulative distribution function (ECDF) area-based relative-distance correlation, with statistical significance assessed by a two-sided interval permutation test (N=1000). Jaccard similarity was additionally calculated as a complementary measure of interval co-localization.

**QTL mapping**

To provide an experimental assessment of effects of genomic regions on the key phenotypes (lifespan and body size), we conducted a pedigree-based QTL mapping experiment based on a large panel of recombinant F2s (i.e., a genotype – phenotype association study). Four newly-hatched virgin males and four virgin females from E and L lines (E2 and L5) were placed in sex-reciprocal E/L pairs to serve as grandparental F0 pairs (N = 4; A - D). These pairs were placed in 8 cm ∅ petri dishes provided with ad libitum *P. vulgaris* beans for oviposition and an Eppendorf vial with 20% saccharose solution, kept at 29$^{o}$ C and 70% RH at 12L:12D. After 12 days, F0s were frozen in -70$^{o}$ C and beans containing F1 larvae were placed individually in virgin chambers (well culturing plates). Emerging virgin F1 beetles were used to form in total 46 F1 pairs. Each pair was also placed in an 8 cm ∅ petri dish provided with ad libitum *P. vulgaris* beans for oviposition and an Eppendorf vial with 20% saccharose solution. Beans containing F2 larvae were then placed individually in virgin chambers and

N = 1334 emerging adult virgin F2 beetles (on average 29 F2 individuals per full-sib F2 family) were used for phenotyping and subsequent sequencing (Fig. S3).

Newly emerged and virgin F2 individuals were sexed and weighed to the nearest 0.01 mg using an electronic microbalance (Sartorius Genius ME 235P-OCE). They were then placed individually in 3.5 ml ([LxØ]: 55 x 12 mm) PS tubes provided with holes in the cap, at 29° C and 70% RH at 12L:12D. All tubes were inspective twice per day to determine lifespan. Beetles were frozen in -70° C upon their death.

For all F0s and F2s, DNA was extracted using a 96-well plate format, using the Qiagen QIAamp DNA Micro kit (Twort et al. 2021). For F0 individuals, sequencing libraries were prepared using the TruSeq PCRfree DNA sample preparation kit (cat# FC-121-3001/3002, Illumina Inc.) targeting an insert size of 350bp. The library preparation was performed according to the manufacturers' instructions (guide#15036187) and these were then sequenced as individual samples using Illumina paired-end sequencing (2 × 150 bp) on a Novaseq 6000 S4 to a depth of ~50X. For all F2s, short read linked-read sequencing libraries were constructed by mixing genomic DNA with a pool of haplotagging beads essentially as described in Meier et al. (2021) and Pal et al. (2024) but with the following modifications. Briefly, each haplotagging bead is carrying on its surface Tn5 transposase assembled with both the i5 and i7 sequencing adapters and four 6-bp segmental barcodes; where one 6-bp barcode is well specific barcode (sample barcode). Two of the four barcode segments were placed in standard i5-index position of a complete Tn5ME-A-adapter sequence accompanied by P5 sequence which attaches the adapter to the bead through streptavidin-biotin bond. The other two 6-bp barcode segments were placed in the beginning of read 2 downstream of a standard i7-TruSeq adapter sequence. The two in-line barcodes were followed by a mutated Tn5-17A/18G-MEmut sequence—ACTTGTGTATAAGAGACAG (Steiniger-White et al. 2002). The mutated Tn5-MEmut sequence allows tagmentation but does not otherwise interfere with Illumina sequencing. An additional standard 8-bp i7 Illumina index barcode was added during the PCR amplification to introduce a fifth barcode segment (i7-index, N701-N714). This additional barcode segment, combined with the original C-segments, made it possible to multiplex more than 96 samples in a single run. Amplified libraries were cleaned up and size-selected using Ampure magnetic beads (Beckman Coulter), Qubit quantified and adjusted with 10 mM Tris, pH 8, 0.1 mM EDTA to 2.5 nM concentration for sequencing. Libraries of F2s were sequenced aiming for 0.5 x coverage with Illumina paired-end sequencing (2 × 150 bp) on one lane of Novaseq 6000 S4, 750 GB sequencing output. Sequences were then processed by trimming away the Tn5-MEmut sequence and demultiplexing the four 6-bp barcode segments as previously described in Meier et al. (2021). For both F0 and F2 individuals, read quality before and after processing was assessed using FastQC v0.11.8 (Andrews et al. 2022). Aggregated FastQC reports were generated and visualized with MultiQC v1.9 (Ewels et al. 2016).

For the F0 individuals, reads were trimmed using Trimmomatic v0.36 (Bolger et al., 2014) to remove low-quality bases (Phred <15), Illumina adapters, and reads <36 bp (ILLUMINACLIP:adapters.fa:2:40:15:8:true; SLIDINGWINDOW:4:15; LEADING:15; TRAILING:15; MINLEN:36). Filtered reads were aligned to the reference genome with BWA-MEM2 v2.2.1 (Vasimuddin et al., 2019), then processed with SAMtools v1.21 (Li et al., 2009; Danecek et al., 2021) for duplicate marking and mate-pair correction. Alignment quality and mapping statistics were assessed using Qualimap v2.3 (García-Alcalde et al., 2012; Okonechnikov et al., 2016). Variant calling was performed using the HaplotypeCaller and GVCFtyper implemented in the sentieon bioinformatics pipeline (Freed et al. 2017). We extracted biallelic SNPs and applied sequential quality control filters to retain only high quality variants. SNPs were hard-filtered using GATK annotations (FS > 60, SOR > 3, MQ < 40, MQRankSum < −12.5, ReadPosRankSum < −8, QD < 2, DP < 3), then polymorphic SNPs with GQ > 20, ≤20% missingness, and MAC ≥ 3 were retained using BCFtools v1.10 (Li et al., 2009). To mitigate depth-related false positives, we applied a per-pool DP filter in R (R Core Team, 2021), retaining only SNPs within the 5th–99th percentile range across all pools. The final SNP set was used for downstream analyses.

For the F2 individuals, we processed the low-coverage haplotagging sequence data following the recommendations, code and barcode files described in the GitHub repositories https://github.com/evolgenomics/haplotagging and https://github.com/evolgenomics/HeliconiusHaplotagging/tree/main. In brief, after read demultiplexing, paired-end reads were trimmed using cutadapt (Marcel, 2011) to remove low-quality bases (--nextseq-trim=20), Illumina adapters (≥15 bp overlap, two rounds), and 5′ primer/anchor sequences (≥19 bp overlap, four rounds), retaining read pairs ≥30 bp and writing shorter reads to separate "too-short" files. For read mapping, we followed a two-step process. First, reads from each of the 14 pooled libraries were mapped to the reference genome with BWA v2.23.4 (Vasimuddin et al., 2019), using the -C option to retain the BX tag, and a single BAM file was generated per pool. Reads were sorted with SAMtools v1.14 (Li et al., 2009; Danecek et al., 2021), and duplicate reads were marked with Picard v2.23.4 using BX-aware options. Second, pooled BAM files were split into individual-level BAM files using an awk script that extracted reads based on the

BX tag. Finally, mapping statistics were generated for each BAM file using Qualimap v.2.3 (García-Alcalde et al. 2012, Okonechnikov et al. 2016) and summarized into a single report with MultiQC v1.12 (Ewels et al. 2016).

We found that non-overlapping QTL bins of 100 kb allowed reasonable genotype assignment in F2s over most of the genome while also providing sufficiently high-resolution mapping. Genotypes in F2s (i.e., EE, EL or LL) was first assigned for QTLs bins based on grandparental genotypes. This was done by comparing each F2 individual with its set of founders, identifying the SNPs that were fixed for opposite alleles in the founders belonging to the L and E lines, respectively. The individual was then scored for genotype concordance with either founder line, and scored for each bin as either LL, LE or EE. At least five informative SNPs needed to be present in the bin to record a call, and at least 75% of the detected SNPs needed to fully support a homozygous call. Custom scripts detailing this procedure are deposited at https://github.com/Goran-Arnqvist/UUBeetleLab/tree/main/Lifespan_selection. As is typical for ultra-low coverage data and outbred founders, however, genotype assignments were noisy and many QTLs showed a significant segregation distortion relative to the expected Mendelian 1:2:1 ratio. Final QTL genotype calls were thus imputed using offline segmentation analysis, by smoothing over data. Here, each genomic bin in an F2 individual is assigned to either EE: 0, EL: 1 or LL: 2. F2 chromosomes was thus modelled as noisy one-dimensional signals composed of long, internally consistent segments punctuated by transitions, with each valid changepoint corresponding to a recombination event during meiosis in F1s. Offline segmentation analysis has been widely applied in high-throughput sequencing contexts, including CNV detection (Olshen *et al.* 2011, Talevich *et al.* 2016). Conceptually, segmentation is an optimization problem in which both the number and location of changepoints are treated as free parameters, as formalized below. Although binary segmentation has historically been the default approach in genomics, the Pruned Exact Linear Time (PELT) (Killick *et al.* 2012) algorithm offers a dynamic programming-based alternative that guarantees an optimal solution while maintaining computational efficiency even on whole-genome sequencing data. The objective function for offline changepoint detection across *T* observations is as

$$\arg \min_{K \in \{1,\dots,T-1\},\{t_k\}} \left\{ \sum_{k=1}^{K} c_{L1}(y_{t_{k-1}:t_k}) + \beta K \right\}$$

Here, the PELT algorithm selects the number (K) and position ($t_k$) of changepoints that result in the global mimimum of the cost function ($C_{L1}$). A penalty term β×K is introduced to prevent oversegmentation. The Segment cost function based on the $L_1$ norm as

$$c_{L1}(y_{a:b}) = \sum_{t=a}^{b} |y_t - median(y_{a:b})|$$

defined as the sum of absolute deviations from the median within each segment, this cost function is used to reduce sensitivity to noise, particularly in heterozygous regions.

Segmentation was performed independently for each chromosome of every F2 individual, using the PELT algorithm as implemented in the ruptures v1.1.9 Python package (Truong *et al.* 2020). The $L_1$ cost model was selected due to its ability to handle noisy heterozygous states in an unbiased manner, and all genomic bins were considered (jump argument in ruptures equal to one). Biologically informed heuristics, including constraints on obligatory recombination, invalid genotype transitions, and crossover interference, were used to filter implausible segments and to guide the tuning of the $\beta$ penalty parameter in the segmentation. A representative example of this approach applied to a single chromosome is shown in Fig. S4.

Following segmentation, imputation was carried out by assigning each segment to its modal genotype state. Chromosome X was treated similarly, with heterozygous transitions in male individuals ignored. This method was extended to all scaffolds larger than 1 Mb, while smaller scaffolds were imputed using heuristic rules. Genotype imputation parameters, particularly the penalty term, were optimized through a finetuning process. Our offline segmentation analysis was very successful and post-imputation genotype assignments entirely resolved segregation distortion (Fig. S5). The resulting imputed dataset (N = 3639 QTL markers genome-wide) was then used for the downstream QTL analyses.

Assuming each detected changepoint corresponds to a single recombination event, this information was utilized to construct fine-scale genetic maps. Genetic distances between loci were estimated using Kosambi's mapping function (Kosambi 1943), which transforms the observed fraction of recombinant gametes into centimorgan units. Cumulative distances along each chromosome were then generated, with linear interpolation applied between adjacent points to estimate intermediate positions. To address the peak-dominated nature of the resulting data, window-based smoothing was applied. The accuracy and structure of the resulting genetic maps were further assessed by comparison with patterns of linkage disequilibrium, calculated using scikit-allel v1.3.13 (Miles *et al.* 2024) and NumPy v2.1.3 (Harris *et al.* 2020). The resulting recombination maps for all assembled autosomes (Fig. S6) is available at https://arnqvist.org/myQTL/recombination_mapping_rerun.html. Average genome-wide recombination rate was estimated to be 1.07 cM/Mb. The recombination map for the X chromosome (i..e, chromosome 10) (https://arnqvist.org/myQTL/chromosome_X_recombination_rate.html) confirmed that the X is acrocentric (Garaud and Lecher 1982, Rozek et al. 1999) and suggested that the assembled scaffold only covers the longer of the two arms (Fig. S6).

QTL mapping for both lifespan and body size was done using both standard interval mapping (IM) and multiple QTL mapping (MQM) models in R/qtl (Broman et al. 2019). The genetic map for QTL mapping was further refined from the previously estimated recombination maps (see above) with the `est.map` function in R/qtl (with an error probability of 0.01). Duplicate markers, in which observed genotypes at one marker matched those at an adjacent location, were removed as non-informative, for a final set of 2,839 informative markers. Genome scans of single QTL models (IM) were conducted on lifespan and body size separately, both on each sex individually as well with full data and sex added as a covariate to the model. A LOD score threshold ($\alpha$=0.05) for candidate QTL on autosomes was calculated per trait by permutation tests (N = 1,000), and separately for the X-chromosome with specific permutations. Multiple QTL mapping was conducted with the R package `qtlTools` (Delaneau et al. 2017) on each of the two traits separately using the function `stepwiseqtl.int`. This function acts similarly to `stepwiseqtl` in R/qtl, but allows for interactions between putative QTL and covariates (i.e., sex) during model construction. Penalties on main effects and interactions for model construction were calculated from permutation tests (n = 1,000) of genome scans of a two-QTL model including sex as an additive and an interactive covariate. Effects of QTL genotype on traits were estimated for all markers in IM models, and for all QTLs identified during modelling in MQM models.

**Intersecting QTLs with signals of evolutionary differentiation**

We performed genome-wide correlations between the LOD score from our global IM model on one hand, and the sweep and CNV call intervals on the other using StereoGene (Stavrovskaya et al. 2017), compiled from source at GitHub (commit b2d817a, corresponding to version 2.50). We used standard parameter settings. Window size was set to 1 Mbp and we used a Kernel width of 100 kbp. Bin size was set to 1000bp, while maxNA and maxZero filtering was set to 100, meaning that only windows that contain no or zero values in at least one track were dismissed. This method correlates genome-wide tracks with discrete features (e.g. sweep or CNV calls) or continuous profiles (e.g. LOD scores) while allowing for control of potential genome-wide confounders by partial correlation (e.g. mappability). The method is based on Gaussian kernel correlations, which provides an estimate of the spatially smoothed correlation of the two features and the statistical significance of correlations is evaluated by a permutation test, comparing the empirical distribution of correlations across the genome with a random distribution composed by spatial permutation of all valid windows.

**Functional annotation and enrichment analysis**

In brief, Functional Gene Ontology (GO) annotations were compiled by combining two complimentary sources, excluding all protein-coding genes with ambiguous or missing chromosomal assignments and retaining only those with valid GO term evidence codes and non-missing annotations. To characterize the functional characteristics of regions, candidate genomic intervals were intersected with coding sequences from the 10 chromosomes. For GO functional analyses, only genes detected in sweeps in at least six replicate samples per EE regime (≥6 out 8) were included, thus placing emphasis on genes consistently targeted by selection. Enrichment analyses followed standard procedures. An identical procedure was applied to the CNVs identified in each EE regime.

Functional Gene Ontology (GO) annotations were compiled by combining two complementary sources. First, InterPro domain assignments were extracted from the NCBI GFF annotation and mapped to GO terms using the interpro2go database (accessed 2025-04-22) (Camon *et al.* 2005). As a second orthogonal source, EggNOG-mapper v2.1.12 (Huerta-Cepas *et al.* 2019, Cantalapiedra *et al.* 2021) was used with the diamond aligner (Buchfink *et al.* 2021) in ultra-sensitive mode to provide supplemental functional annotations. The final GO annotation set (two references to GO) was derived from the union of both sources. Obsolete or deprecated GO terms were systematically removed or replaced using GOATools v1.4.12 (Klopfenstein *et al.* 2018), referencing the go-basic.obo ontology file (https://purl.obolibrary.org/obo/go/go-basic.obo, downloaded on 22-04-2025). Protein-coding genes with ambiguous or missing chromosomal assignments were excluded from the final annotation set.

GO term metadata—including definitions and ontology classifications—were obtained from the GO.db package v3.20.0 (Carlson 2017) and merged with the gene-level annotations to generate a comprehensive annotation table. Chromosomal coordinates were assigned to each gene, and quality control flags such as low coverage, low mapping quality, and proximity to assembly gaps based on the blacklist described above were appended. Only protein-coding genes with valid GO evidence codes and non-missing annotations were retained. The final annotation was compiled into an organism-specific package with AnnotationForge (Marc Carlson 2017, 2025) to enable reproducible downstream enrichment analyses.

To characterize the functional implications of selective sweeps, candidate genomic intervals were intersected with coding sequences (CDS) from the *A. obtectus* genome to identify potentially selected genes. Only genes located within the ten major assembled chromosomal scaffolds were included in the analysis. Any gene overlapping regions in the blacklist described above was excluded from both the candidate gene list and the background universe, in accordance with best practices for enrichment analysis (Yon Rhee *et al.* 2008, Tipney & Hunter 2010). Candidate gene lists for GO functional analysis were compiled by selecting genes that overlapped selective sweep regions within the ten major assembled scaffolds and were detected in at least six replicate samples (≥6 occurrences) per regime, thus focussing on consistently selected loci. Over Representation Analysis (ORA) was performed using the enrichGO function from ClusterProfiler v4.14.6 (Yu *et al.* 2012, Wu *et al.* 2021) to identify enriched functional categories potentially targeted by selection. Multiple testing correction was applied using the Benjamini-Hochberg procedure with $\alpha = 0.05$, and results were filtered to retain only terms with $P < 0.01$ and $Q < 0.05$. Redundant GO terms were pruned using the simplify function (cut-off = 0.7; Wang semantic similarity) from GOSemSim v2.32.0 (Yu *et al.* 2010). Visualization of results was conducted using clusterProfiler and enrichplot v1.26.6 (Guangchuang Yu 2018).

A procedure identical to the one described above was applied to the CNVs identified in each selective regime, with the exception that a gene was considered a candidate only if at least 50% of its length overlapped with a CNV. Deletions and copy number gains were analysed separately for the E and the L EE regimes.

**MultiGo analyses**

To identify convergent signals across our multi-omics datasets, we performed an integrated gene-level analysis within the Early (E) and Late (L) regimes. Candidate loci were defined from consistently detected selective sweeps and from CNVs intervals, called as described above. *Drosophila melanogaster* represents the closest well-studied and widely curated insect model for lifespan genetics to *A. obtectus*. To examine and contextualize our results, we also considered (i) all annotated longevity-associated genes in GenAge for *D. melanogaster* (https://genomics.senescence.info/genes/search.php?organism=melanogaster – retrieved 2025-10-30), and (ii) candidate gene sets from experimental evolution studies with a design qualitatively similar to ours (Remolina et al. 2012, Carnes et al. 2015, Fabian et al. 2018 ). Orthologs were assigned using InParanoid-DIAMOND v5.1 (Remm et al. 2001, Buchfink et al. 2021, Persson and Sonnhammer 2022, retaining only the top-scoring ortholog(s) per orthologous group (ties retained). Finally , we incorporated differentially expressed genes (DEGs) from previous work comparing gene expression between E and L both in the abdomen and in the head/thorax, separately in the two sexes (Immonen et al. 2023).

To ensure comparability across datasets, all evidence layers were mapped to a unified set of protein-coding genomic features from the final Hi-C scaffolded assembly. Transcript-derived features from Immonen et al. (2023) were assigned to protein-coding genes by local alignment against the full set of coding sequences (CDS) using MegaBLAST from the BLAST+ suite v 2.17.0 (Altschul et al. 1990, Zhang et al. 2000, Camacho et al. 2009), requiring ≥98% sequence identity and ≥80% CDS coverage. We note that the transcriptome in Immonen et al. (2023) was generated using an earlier, non-scaffolded version of the genome assembly.

Because each evidence layer has a distinct source, we defined a background universe for each category to avoid biased overlap estimates. Universes were specified as follows: (i) Selective sweeps; all protein-coding features within the nine major autosomal scaffolds, (ii) CNVs; all non-blacklisted protein-coding features within the ten chromosomal scaffolds, (iii) DEGs; all genomic features with an assigned transcript, and (iv) *Drosophila* orthologs; all genomic features with an assigned ortholog.

We then focused on the L (long-lived) regime and visualized evidence overlap across gene sets using the UpSetPlot Python library [Lex] v 0.9.0. For each evidence category, gene identifiers were merged into regime-specific sets, forming an incidence matrix. To quantify whether overlaps exceeded random expectation, we calculated a $\log_2$ enrichment ratio for each intersection relative to a null model assuming random assortment, conditional on their shared background universe. For any combination of $n$ evidence sets, we defined the common universe ($U_c$) as the intersection of their respective background universes:

$$U_c = \bigcap_{i=1}^{n} U_i$$

The expected fraction of overlapping genes under the null hypothesis was computed as the product of the individual set frequencies within $U_c$:

$$E_{fraction} = \prod_{i=1}^{n} \frac{|S_i \cap U_c|}{|U_c|}$$

The enrichment ratio was then defined as the ratio of the observed intersection fraction to the expected fraction:

$$EnrichmentRatio = \frac{\frac{|(\bigcap_{i=1}^{n} S_i) \cap U_c|}{|U_c|}}{E_{fraction}}$$

Values of $\log_2 Enrichment\ ratio\ > 0$ indicate overlap beyond stochastic expectation. However, $E_{fraction}$ is unstable for small intersection sizes and should therefore be interpreted with caution.

**Multi-evidence functional gene set ranking**

Several results from the functional analyses of our candidate gene sets shared enriched biological process GO terms with previous work on gene expression (Immonen et al. 2023).

To systematically explore whether functional properties of genes in selective sweep and CNV calls overlap with those of DEGs, we implemented a custom multi-evidence Gene Ontology (GO) term ranking framework. Evidence was organized into two categories: selection and expression. Selection evidence included two regime-specific layers (candidate gene sets) per EE regime: (i) genes consistently overlapping candidate selective sweep intervals and (ii) genes overlapping differentiated copy-number variants (both deletions and gains), as defined above. Expression evidence was comprised of four sex × tissue DEG sets derived from E vs L contrasts as reported by Immonen et al. (2023), with DEG transcripts mapped to genomic features as described above. Because expression evidence is defined from direct E–L contrasts, the Expression category is the same when scoring Early versus Late regimes.

For each layer, genes/features in the layer-specific background universe (post-filtering) were represented as a binary vector (hit = 1, non-hit = 0). To make the layers comparable, features absent from a given layer universe were treated as missing values (“structural zeros”).

For each GO term $t$ and layer $l$, Z-scores were calculated under a hypergeometric model. Accounting for structural zeros, let $N_l$ be the number of genes in the universe, $n_l$ the number of hits, $K_{t,l}$ the gene set size of $t$, and $x_{t,l}$ the number of hits annotated to $t$. The expected hits and variance were defined as:

$$\mathrm{m}_{t,l} = n_l \cdot \frac{K_{t,l}}{N_l}, \qquad \mathrm{s}_{\mathrm{t,l}}^2 = \mathrm{n_l} \cdot \frac{\mathrm{K_{t,l}}}{\mathrm{N_l}}\left(1 - \frac{\mathrm{K_{t,l}}}{\mathrm{N_l}}\right) \cdot \frac{\mathrm{N_l} - \mathrm{n_l}}{\mathrm{N_l} - 1}$$

To ensure smooth downstream integration, and since underrepresentation signals can be influenced by technical biases (low expression, low-quality regions etc.), the negative scores were capped to non-negative values:

$$Z_{t,l} = \max\left(0, \frac{x_{t,l} - \mathrm{m}_{t,l}}{\sqrt{\mathrm{s}_{t,l}^2}}\right)$$

Within each evidence category $G$, Z-scores were aggregated using a normalized p-norm, with $p = 3.0$:

$$S_t^{(G)} = \left(\frac{1}{|G|}\sum_{l \in G}\left(Z_{t,l}\right)^p\right)^{1/p}, \; G \in \{\text{Selection, Expression}\}$$

Regime-specific final scores were computed as the sum of the Selection score for the corresponding regime and the shared Expression score:

$$S_t^{Early} = S_t^{Selection\ Early} + S_t^{Expression}, \qquad S_t^{Late} = S_t^{Selection\ Late} + S_t^{Expression}$$

Statistical significance per regime was assessed with a one-sided test from an empirical distribution for $S_t$ through permutations of the Gene-GO term annotation labels. GO terms annotated to fewer than five genes or more than 500 genes were excluded. Multiple testing correction was performed using the Benjamini–Hochberg FDR procedure with $a = 0.05$ (statsmodels v0.14.5). Terms significant in both regimes were designated “shared”. All significant GO terms (Early, Late, and shared) were clustered based on gene set overlap in Cytoscape v3.10.4 using clustermaker2 v2.3.4, with the Markov Cluster Algorithm (MCL) and inflation parameter 2.2, after pruning network edges for overlap ≥ 0.5.

**LLM assistance**

Large Language Models (LLMs) were used during the development of some analysis pipelines and scripts. This included assistance with debugging and clarification of documentation. The models used were GPT-4o (OpenAI 2024a), GPT-o1 (OpenAI 2024b), GPT-o3-mini (OpenAI 2025a), GPT-o3 (OpenAI 2025b), and GPT-o4-mini (OpenAI 2025c). GitHub Copilot (GitHub, Inc. and OpenAI 2021) was also used for code completion and boilerplate suggestions. All generated code was reviewed, edited, and adapted as necessary to fit the specific context of the analyses.

##### Genome browser

An interactive genome browser (JBrowse2) containing selected genomic tracks from the Early and Late selection lines is publicly available at:
https://arnqvist.org/obtectus_genome-browser/index.html?config=config.json
Tracks are organized by evidence category, with brief descriptions provided for each. All source data visualized in the displayed tracks are deposited at:
https://doi.org/10.17044/scilifelab.34059444

#### 2. Supplementary Results

##### Genetic differentiation

A genome scan of the genetic differentiation between E and L lines, across all chromosomes, is given in Fig. S7.

##### Selective sweeps

The ECDF correlation matrix, based on the Jaccard spatial similarity matrix of the genome-wide signal of selective sweeps, is illustrated in Fig. S8. We note that samples from the E regime cluster closer together than those from the L regime.

##### Copy-number-variants

The degree of read-depth differentiation between samples is illustrated in Figs. S9 and S10. The pipe-line used to call CNVs is illustrated in Fig. S11. The size distribution of called CNVs is illustrated in Fig. S12.

##### QTL mapping

Please see our main article for the overall results of the QTL mapping. Supplementary results are illustrated in Figs. S13-S27. Parameter estimates of multiple QTL mapping models are given in Table S2 and Table S3. The sex-specific interval mapping models (Figure 5) showed the following significant QTL peaks; females weight: (chromosome:window_start–window_end) 2:88,331,884–88,431,884; male weight : 2:31,799,142–31,899,142; 3:37,443,485–37,543,484; 4:103,344,924–103,444,923;

9:28,995,489–29,052,433; females for lifespan: 2:88,331,884–88,431,883; 4:37,340,105–37,440,105; 7:2,906,321–3,006,320; males for lifespan: 2:14,146,184–14,246,183; 4:6,201,107–6,301,106; 7:53,922,540–54,022,539. The multiple QTL mapping models revealed significant QTLs at: (chromosome:window_start–window_end) for body weight: 2:15,346,184–15,446,183; 4:12,748,327–12,848,326; 4:14,405,940–14,505,939; 6:6,301,250–6,401,249; 9:69,103,912–69,203,911; 9:72,703,912–72,803,911; and for lifespan: 2:88,331,884–88,431,883; 4:17,105,940–17,205,939; 5:61,394,889–61,494,888; 6:85,522,833–85,622,832; 7:3,306,321–3,406,320; 9:58,286,282–58,386,281.

**Enrichment and multi-evidence functional gene set ranking**

The results of the GO term enrichment analyses of selective sweeps and CNVs are given in Figs. 28-34. Our multi-evidence functional gene set ranking resulted in a rich set of functional modules which were identified as being associated with the observed evolution of lifespan. These were all enriched in more than one data layer and involved a large number of functional categories canonically associated with lifespan, as illustrated in Figs. S35-37. These modules included, for example, immune function and wound healing, oocyte growth and development, mitochondrial function, mitotic spindle organization, formation and differentiation of blood cells, morphogenesis of male genitalia, chemosensation and behavior, brain development, regulation of circadian rhythm, response to stress (oxidative, thermal and osmotic), transcription and RNA catabolism/metabolism, regulation of carbohydrate metabolism, toll signaling, muscle and neural tissue development, DNA repair and replication, cell cycle regulation, various metabolic and catabolic processes, cGMP and kinase signaling, apoptosis, protein disassembly and phagocytosis.

**Intersection of QTLs with signals of evolutionary differentiation**

The weak genomic association between CNV calls and selective sweep intervals is illustrated in Fig. S38. The mean genome-wide spatial correlations between LOD scores and sweep/CNV calls were generally sizeable (see main text), for both lifespan and body size. The empirical foreground distributions of correlations were very highly significantly different from the random expectation in all cases (Fig. S39). Inspection of diagnostic cross-correlation plots showed that these were smooth and symmetrical with a clear peak around 0, as expected for co-localize features in the correlated tracks.

**3. Supplementary References**

Altschul SF, Gish W, Miller W, Myers EW, Lipman DJ (1990) Basic local alignment search tool. J Mol Biol 215:403–410.

Andrews S, Krueger F, Segonds-Pichon A, Biggins L, Krueger C, Wingett S (2022) FastQC: a quality control tool for high throughput sequence data. [https://www.bioinformatics.babraham.ac.uk/projects/fastqc/].

Bergland AO, Behrman EL, O'Brien KR, Schmidt PS, Petrov DA (2014) Genomic evidence of rapid and stable adaptive oscillations over seasonal time scales in Drosophila. PLoS Genet 10:e1004775.

Bolger AM, Lohse M, Usadel B (2014) Trimmomatic: a flexible trimmer for Illumina sequence data. Bioinformatics 30:2114–2120.

Broad Institute (2019) Hard-filtering germline short variants. Available at: https://gatk.broadinstitute.org/hc/en-us/articles/360035890471-Hard-filtering-germline-short-variants.

Buchfink B, Reuter K, Drost HG (2021) Sensitive protein alignments at tree-of-life scale using DIAMOND. Nat Methods 18:366–368.

Camacho C, Coulouris G, Avagyan V, Ma N, Papadopoulos J, Bealer K, Madden TL (2009) BLAST+: Architecture and applications. BMC Bioinformatics 10:421.

Danecek P, Bonfield JK, Liddle J, Marshall J, Ohan V, Pollard MO, Whitwham A, Keane T, McCarthy SA, Davies RM, Li H (2021) Twelve years of SAMtools and BCFtools. GigaScience 10:giab008.

Delaneau O, Ongen H, Brown AA, et al. (2017) A complete tool set for molecular QTL discovery and analysis. Nat Commun 8:15452.

Dudchenko O, Batra SS, Omer AD, Nyquist SK, Hoeger M, Durand NC, et al. (2017) De novo assembly of the Aedes aegypti genome using Hi-C yields chromosome-length scaffolds. Science 356:92–95.

Durand NC, Shamim MS, Machol I, Rao SS, Huntley MH, Lander ES, Aiden EL (2016) Juicer provides a one-click system for analyzing loop-resolution Hi-C experiments. Cell Syst 3:95–98.

Ewels P, Magnusson M, Lundin S, Käller M (2016) MultiQC: summarize analysis results for multiple tools and samples in a single report. Bioinformatics 32:3047–3048.

Favorov A, Mularoni L, Cope LM, Medvedeva Y, Mironov AA, Makeev VJ, Wheelan SJ (2012) Exploring massive genome scale datasets with the GenometriCorr package. PLoS Comput Biol 8:e1002529.

Feder AF, Petrov DA, Bergland AO (2012) LDx: Estimation of linkage disequilibrium from high-throughput pooled resequencing data. PLoS ONE 7:e48588.

Freed D, Aldana R, Weber JA, Edwards JS (2017) The Sentieon genomics tools – a fast and accurate solution to variant calling from next-generation sequence data. bioRxiv:115717.

Garaud P, Lecher P (1982) Etude biométrique du caryotype de la bruche du haricot (Acanthoscelides obtectus, Coléoptéres, Bruchidae). Can J Genet Cytol 24:687–692.

García-Alcalde F, Okonechnikov K, Carbonell J, Cruz LM, Götz S, Tarazona S, Dopazo J, Meyer TF, Conesa A (2012) Qualimap: evaluating next-generation sequencing alignment data. Bioinformatics 28:2678–2679.

Hivert V, Leblois R, Petit EJ, Gautier M, Vitalis R (2018) Measuring genetic differentiation from pool-seq data. Genetics 210:300900.

Iskow RC, Gokcumen O, & Lee C (2012) Exploring the role of copy number variants in human adaptation. Trends in Genetics, 28(6), 245-257.

Kolaczkowski B, Kern AD, Holloway AK, Begun DJ (2011) Genomic differentiation between temperate and tropical Australian populations of Drosophila melanogaster. Genetics 187:245–260.

Li H, Handsaker B, Wysoker A, Fennell T, Ruan J, Homer N, et al. (2009) The sequence alignment/map format and SAMtools. Bioinformatics 25:2078–2079.

Martin M (2011) Cutadapt removes adapter sequences from high-throughput sequencing reads. EMBnet J 17:10–12.

McKenna A, Hanna M, Banks E, Sivachenko A, Cibulskis K, Kernytsky A, et al. (2010) The genome analysis toolkit: a MapReduce framework for analyzing next-generation DNA sequencing data. Genome Res 20:1297–1303.

Meier JI, Salazar PA, Kučka M, et al. (2021) Haplotype tagging reveals parallel formation of hybrid races in two butterfly species. Proc Natl Acad Sci USA 118:e2015005118.

Okonechnikov K, Conesa A, García-Alcalde F (2016) Qualimap 2: advanced multi-sample quality control for high-throughput sequencing data. Bioinformatics 32:292–294.

Oksanen J, Simpson GL, Blanchet FG, Kindt R, Legendre P, Minchin PR, ... & Weedon J. (2022) vegan: Community ecology package (2.6-4). CRAN.

Olshen AB, Bengtsson H, Neuvial P, Spellman PT, Olshen RA, & Seshan VE (2011) Parent-specific copy number in paired tumor–normal studies using circular binary segmentation. Bioinformatics, 27(15), 2038-2046.

Pal A, Shipilina D, Le Moan A, McNairn AJ, Grenier JK, Kucka M, Coop G, Chan YF, Barton NH, Field DL, Stankowski S (2025) Genealogical analysis of replicate flower colour hybrid zones in Antirrhinum. Mol Ecol 34:e70067.

Persson E, Sonnhammer ELL (2022) InParanoid-DIAMOND: Faster orthology analysis with the InParanoid algorithm. Bioinformatics 38:2918–2919.

R Core Development Team (2021) R: A language and environment for statistical computing. R Foundation for Statistical Computing, Vienna, Austria.

Remm M, Storm CEV, Sonnhammer ELL (2001) Automatic clustering of orthologs and in-paralogs from pairwise species comparisons. J Mol Biol 314:1041–1052.

Rideout JR, Caporaso G, Bolyen E, McDonald D, Baeza YV, Alastuey JC, ... & Radinger J (2024) scikit-bio/scikit-bio: scikit-bio 0.6.1. Zenodo.

Rozek M, Holecová M, Lachowska D (1999) Karyological notes on Acanthoscelides obtectus (Say 1859) (Coleoptera, Bruchidae). Cytologia 64:367–369.

Twort VG, Minet J, Wheat CW, Wahlberg N (2021) Museomics of a rare taxon: placing Whalleyanidae in the Lepidoptera tree of life. Syst Entomol 46:926–937.

Weir BS, Cockerham CC (1984) Estimating F-statistics for the analysis of population structure. Evolution 38:1358.

Wickham H (2016) ggplot2: Elegant graphics for data analysis. Springer, New York.

Zhang Z, Schwartz S, Wagner L, Miller W (2000) A greedy algorithm for aligning DNA sequences. J Comput Biol 7:203–214.

**Table S1.** Assembly and annotation statistics for the new genome assembly of *A. obtectus* . Completeness was assessed with BUSCO v5.8.2, using the insecta_odb10 reference gene

| | New HiC assembly (GCA_963669975.1) |
|---|---|
| Genome size | 1.1 Gb |
| Total ungapped length | 1.1 Gb |
| Number of scaffolds | 3,796 |
| Scaffold N50 | 108.7 Mb |
| Scaffold L50 | 5 |
| Number of contigs | 7,275 |
| Contig N50 | 772.9 kb |
| Contig L50 | 391 |
| GC % | 37.5 |
| Genome coverage | 80x |
| Total genes | 37,346 |
| Protein-coding genes | 36,598 |
| BUSCO assessment (n=1367): | |
| Complete BUSCOs ( C ) | 98.6% (n=1348) |
| Single-copy BUSCOs (S) | 86.6% (n=1184) |
| Duplicated BUSCOs ( D ) | 12.0% (n=164) |
| Fragmented BUSCOs ( F ) | 0.4% (n=5) |

**Table S2**. Parameter estimates of the multiple QTL model for weight. The additive (denoted with 'a') and dominance (denoted with 'd') estimates are presented for each QTL in the model. Significant parameters are denoted with an asterisk.

| **Parameter** | **Estimate (Weight, mg)** | **Standard Error** | ***t*** |
|---|---|---|---|
| Intercept | 5.608 | 0.027 | 204.725* |
| sex | -0.872 | 0.037 | -23.500* |
| 2@22.9a | 0.146 | 0.026 | 5.588* |
| 2@22.9d | 0.028 | 0.037 | 0.748 |
| 9@90.5a | 0.405 | 0.126 | 3.217* |
| 9@90.5d | 0.426 | 0.146 | 2.924* |
| 4@28.3a | 0.487 | 0.076 | 6.414* |
| 4@28.3d | 0.070 | 0.081 | 0.860 |
| 4@25.3a | -0.383 | 0.077 | -4.979* |
| 4@25.3d | -0.106 | 0.081 | -1.304* |
| 9@89.4a | -0.284 | 0.126 | -2.255* |
| 9@89.4d | -0.444 | 0.145 | -3.052* |
| 6@12.6a | 0.098 | 0.026 | 3.734* |
| 6@12.6d | 0.082 | 0.037 | 2.202* |

**Table S3**. Parameter estimates of the multiple QTL model for lifespan. The additive (denoted with 'a') and dominance (denoted with 'd') estimates are presented for each QTL in the model. Significant parameters are denoted with an asterisk. Only significant additive marker:sex interactions are shown.

| **Parameter** | **Estimate (Lifespan, days)** | **Standard Error** | ***t*** |
|---|---|---|---|
| Intercept | 23.60 | 0.21 | 111.52* |
| sex | -6.15 | 0.29 | -21.36* |
| 4@33.1a | 1.31 | 0.19 | 6.79* |
| 4@33.1d | -0.23 | 0.28 | -0.81 |
| 2@82.0a | 1.33 | 0.28 | 4.74* |
| 2@82.0d | -1.62 | 0.42 | -3.88* |
| 7@5.8a | 1.49 | 0.30 | 5.04* |
| 7@5.8d | 0.99 | 0.42 | 2.38* |
| 9@80.3a | 0.80 | 0.27 | 3.03* |
| 9@80.3d | -1.68 | 0.42 | -3.96* |
| 5@56.1a | 1.55 | 0.29 | 5.30* |
| 5@56.1d | 0.22 | 0.42 | 0.52 |
| 6@125.3a | 0.41 | 0.19 | 2.08* |
| 6@125.3d | -1.05 | 0.28 | -3.70* |
| 2@82.0a:sex | -0.61 | 0.39 | -1.57 |
| 2@82.0d:sex | 1.62 | 0.57 | 2.86* |
| 7@5.8a:sex | -1.15 | 0.40 | -2.89* |
| 7@5.8d:sex | -1.20 | 0.56 | -2.13* |
| 9@80.3a:sex | -0.29 | 0.36 | -0.81 |
| 9@80.3d:sex | 1.84 | 0.58 | 3.20* |
| 5@56.1a:sex | -1.34 | 0.40 | -3.40* |
| 5@56.1d:sex | -0.30 | 0.57 | -0.52 |

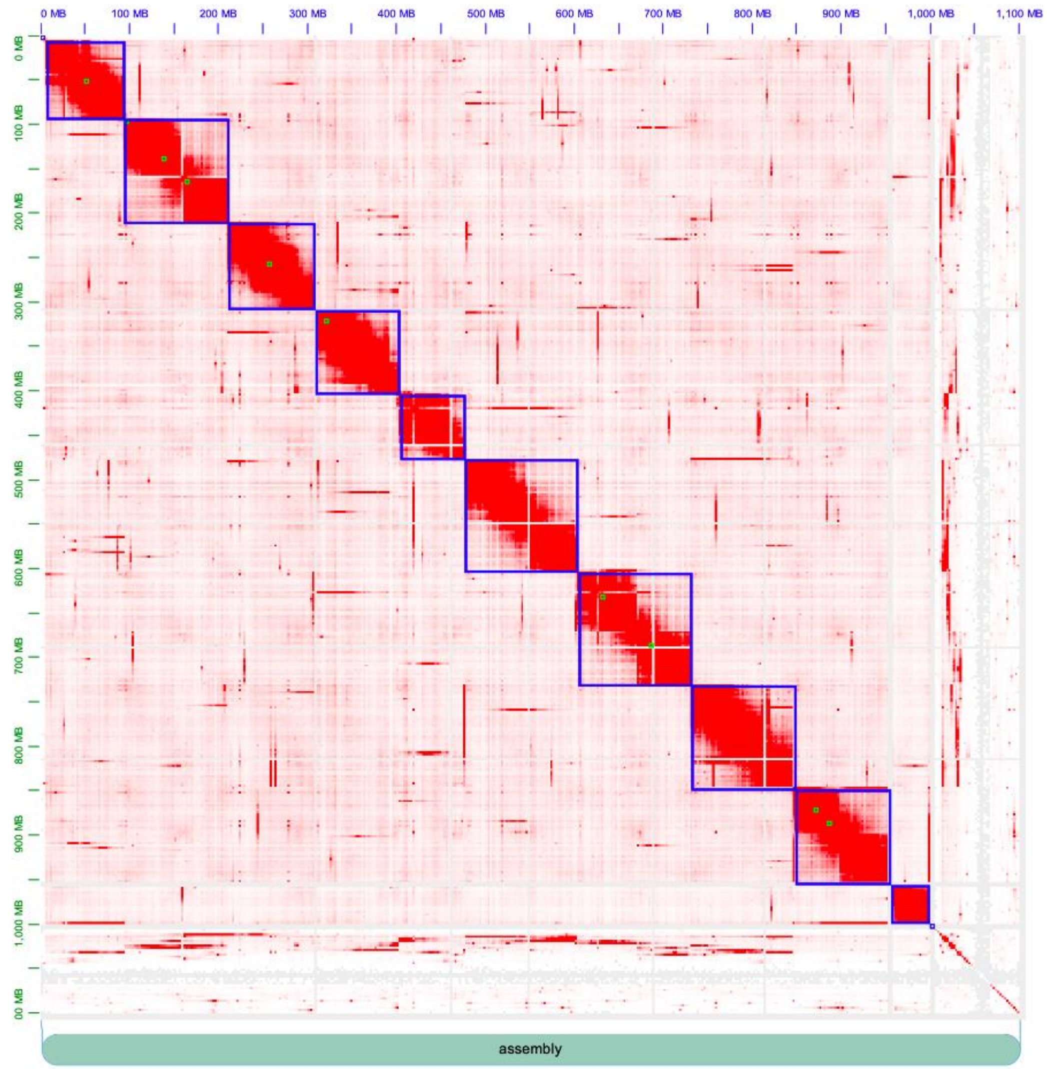

**Figure S1**. Hi-C contact map on the scaffolded A. obtectus assembly, where >90% of the input assembly was arranged into 10 chromosome-length scaffolds.

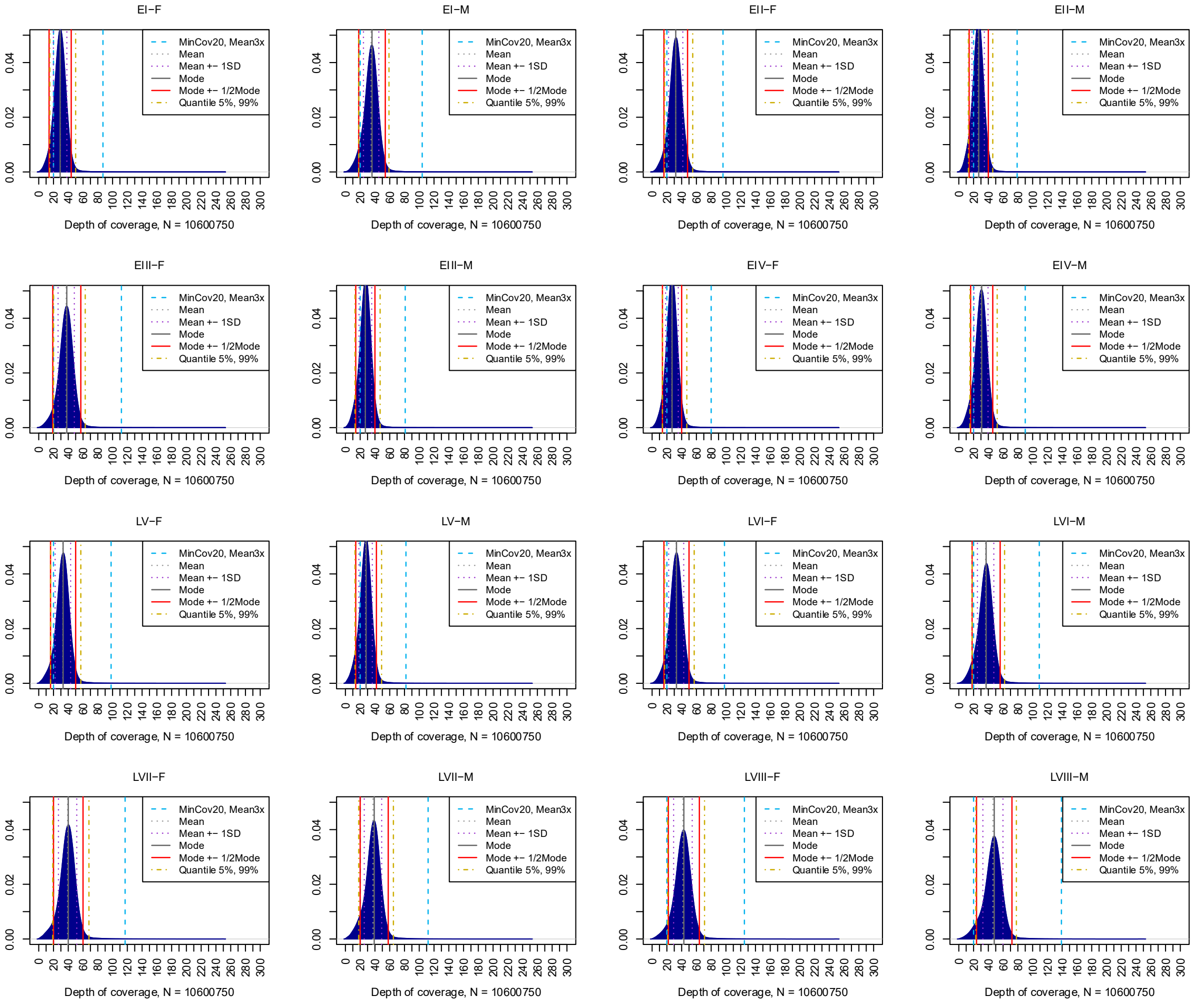

Fig. S2. Distribution of sequencing depth across *A. obtectus* sequencing pools based on about 10.6 million SNPs that passed quality filters. Vertical lines indicate the different lower and upper depth-of-coverage thresholds evaluated.

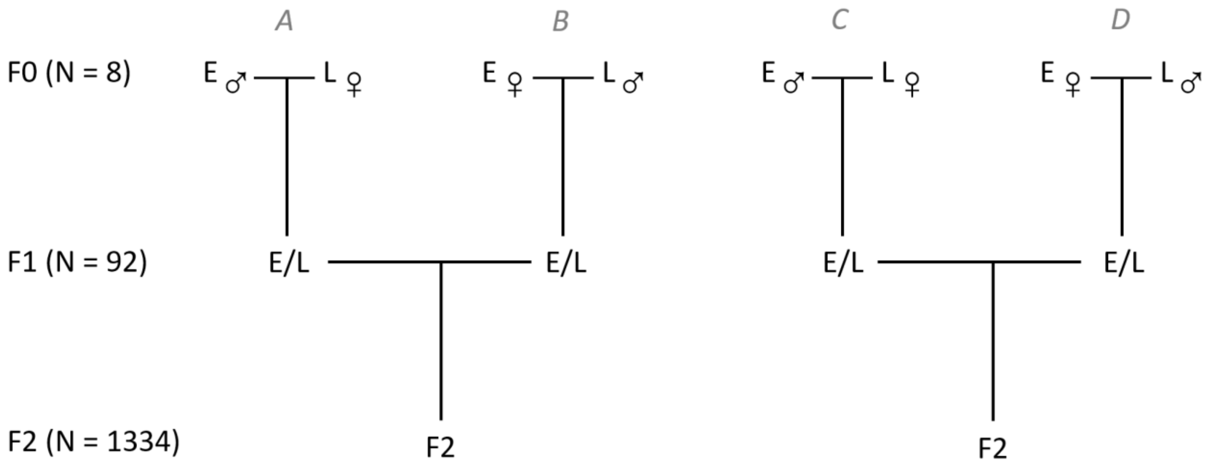

**Fig. S3.** Schematic representation of the QTL pedigree, where four F0 families were used to create 46 pairs of heterozygote F1s which, in turn, generated 1334 focal recombinant F2 individuals.

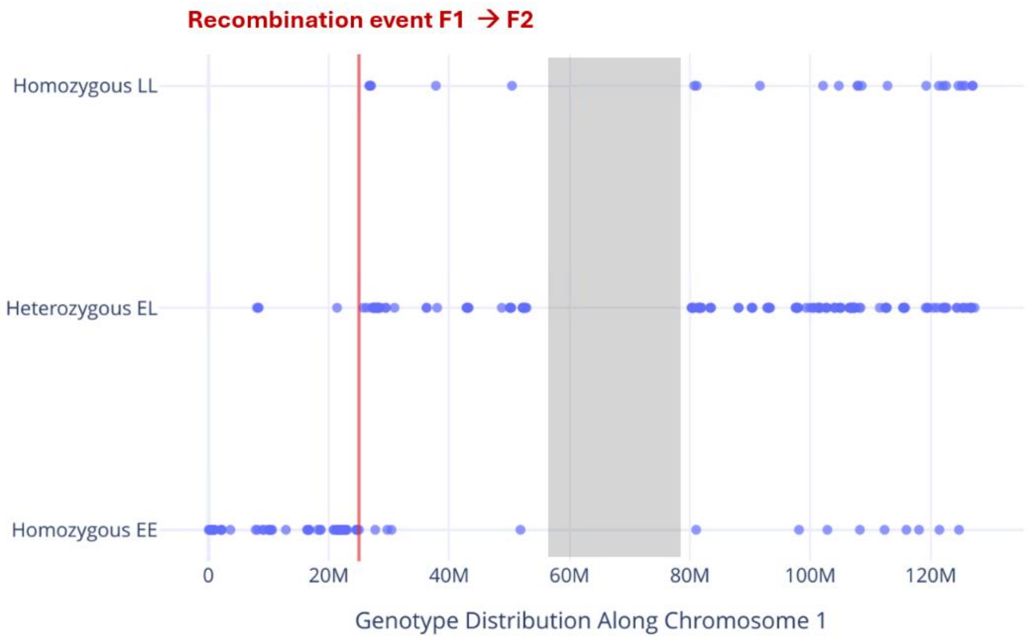

**Fig. S4.** An example of offline segmentation of chromosome 1 in a single problematic F2 individual. One recombination event is detected, transitioning from a homozygous EE to a heterozygous EL genotype. The greyed-out region was excluded in this example due to lack of segregating information, corresponding to presumed location of the centromere.

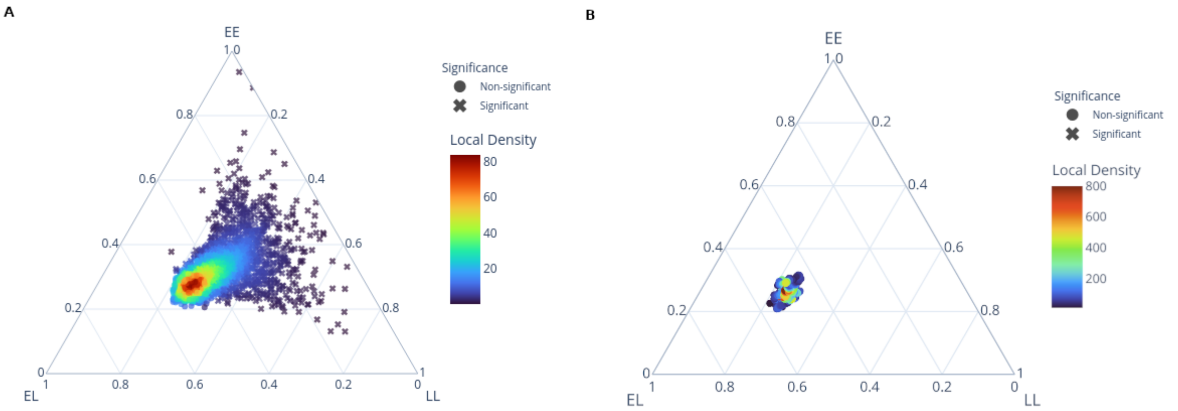

**Fig. S5.** De Finetti-style diagram showing segregation distortion in grandparental origin of F2 genotype calls. Panels show QTL genotype calls before (A) and after (B) imputation. Point density is visualized using a Gaussian kernel estimate. Significant deviations from expected Mendelian segregation were identified before imputation using a chi-square test (α = 0.05, Benjamini–Hochberg corrected).

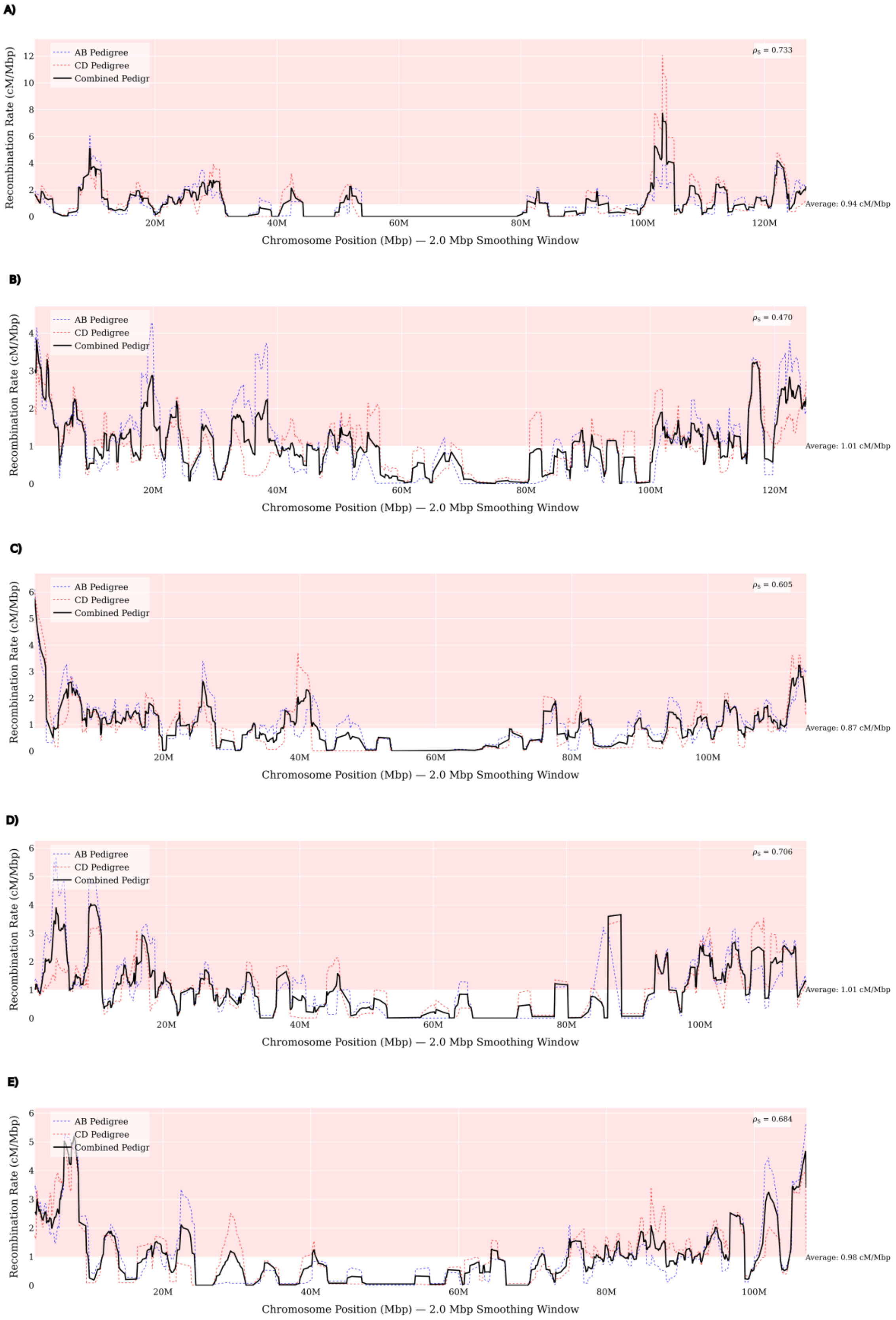
A)
AB Pedigree
CD Pedigree
Combined Pedigr
$\rho_S$ = 0.733
Average: 0.94 cM/Mbp
Recombination Rate (cM/Mbp)
Chromosome Position (Mbp) — 2.0 Mbp Smoothing Window
B)
AB Pedigree
CD Pedigree
Combined Pedigr
$\rho_S$ = 0.470
Average: 1.01 cM/Mbp
Recombination Rate (cM/Mbp)
Chromosome Position (Mbp) — 2.0 Mbp Smoothing Window
C)
AB Pedigree
CD Pedigree
Combined Pedigr
$\rho_S$ = 0.605
Average: 0.87 cM/Mbp
Recombination Rate (cM/Mbp)
Chromosome Position (Mbp) — 2.0 Mbp Smoothing Window
D)
AB Pedigree
CD Pedigree
Combined Pedigr
$\rho_S$ = 0.706
Average: 1.01 cM/Mbp
Recombination Rate (cM/Mbp)
Chromosome Position (Mbp) — 2.0 Mbp Smoothing Window
E)
AB Pedigree
CD Pedigree
Combined Pedigr
$\rho_S$ = 0.684
Average: 0.98 cM/Mbp
Recombination Rate (cM/Mbp)
Chromosome Position (Mbp) — 2.0 Mbp Smoothing Window

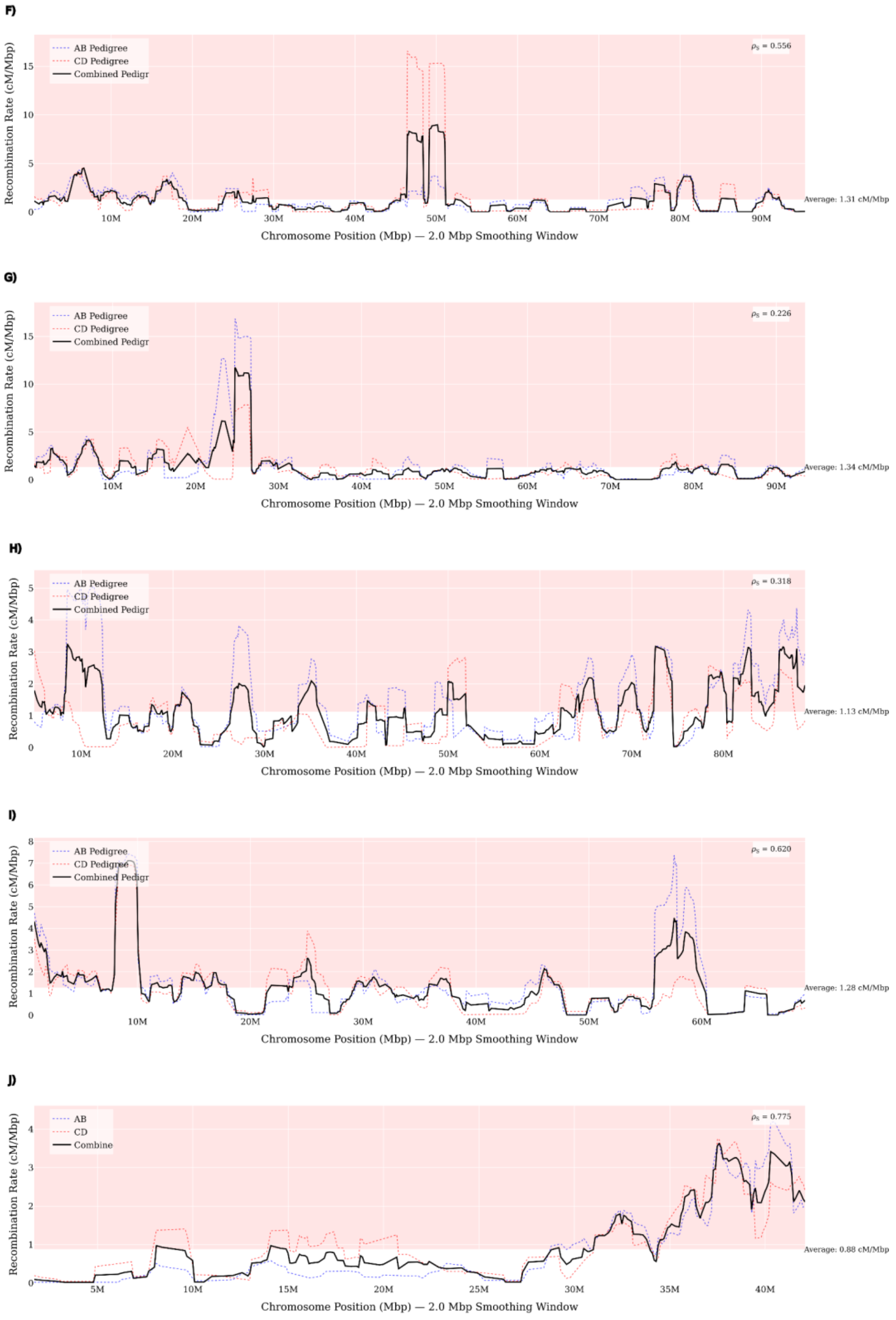

**Fig. S6.** Recombination rate (cM/Mb) across the nine autosomes (A–I) and the X (J). Smoothed values for each half of the pedigree (AB and CD) are shown as well as Spearman rank correlations ($\rho_s$) between the two halves is indicated in the legend. A more detailed genetic map is available at: https://arnqvist.org/myQTL/recombination_mapping_rerun.html or upon request from the corresponding author.

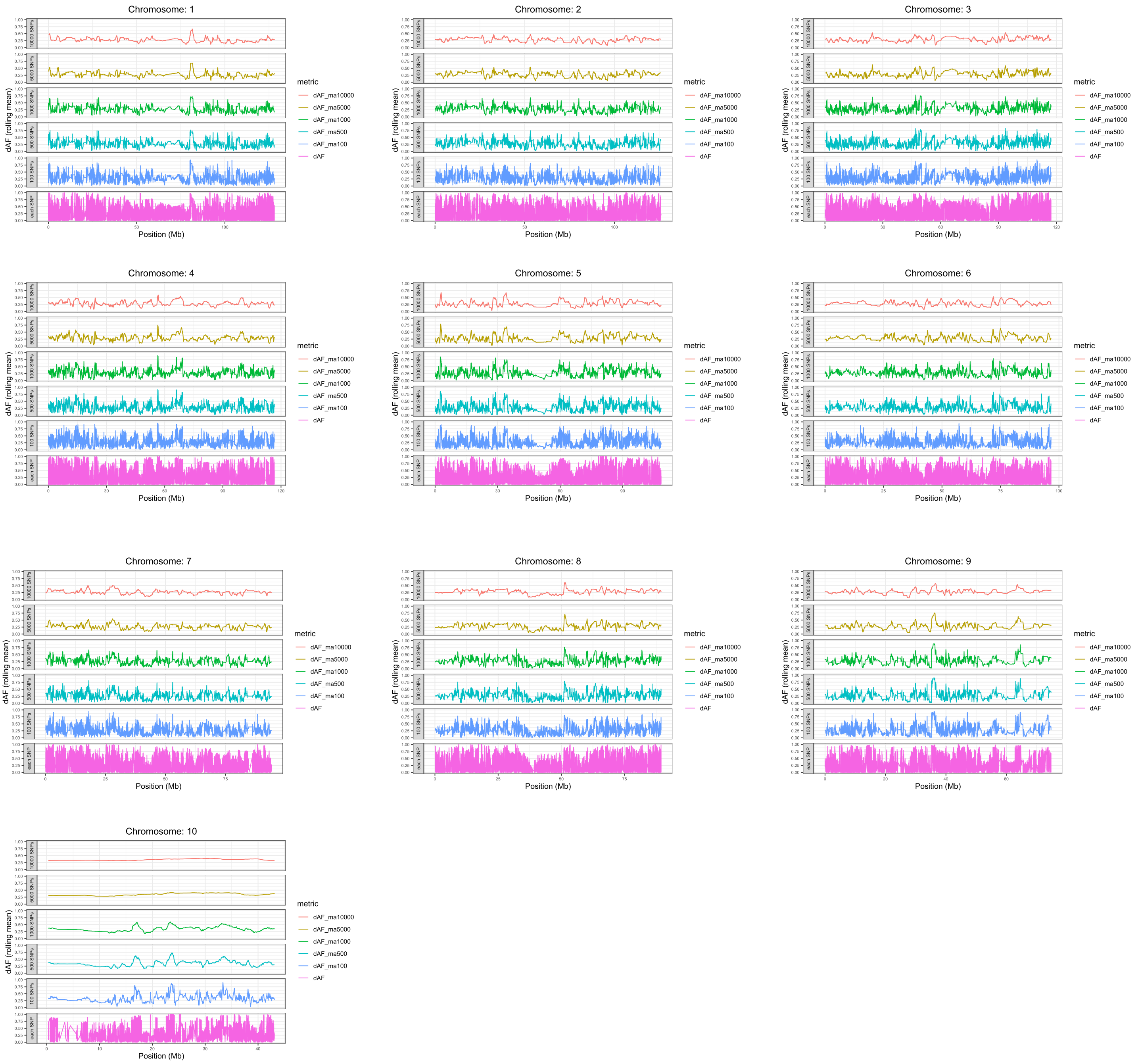

**Fig. S7.** Genome scans illustrating the genomewide differences in SNP allelefrequencies ( dAF; range 0 - 1) between superpools of E and L lines across the genome. Graphs show show differentiation along each chromosome as a rolling average of various window sizes (window sizes bottom up: 1, 100, 500, 1000, 5000 and 10000 bp). Chromosome 10 represents the X chromosome.

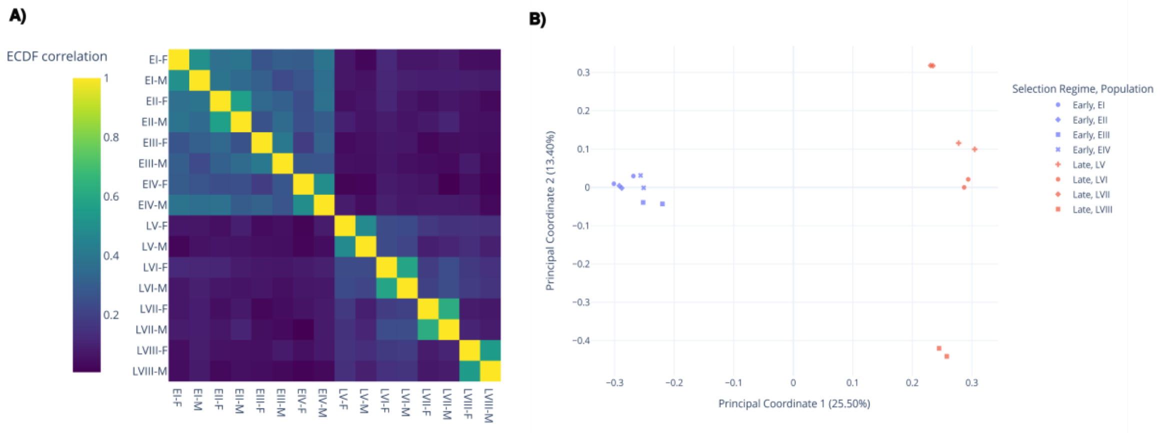

**Fig. S8.** Similarity of selective sweep signatures across samples. (A) Heatmap of ECDF correlations, computed with GenometriCorr. Values near zero indicate random association; values near one indicate perfect correspondence. (B) Principal Coordinate Analysis (PCoA) of Jaccard distances among all samples, illustrating clear clustering by EE regime along PCo1.

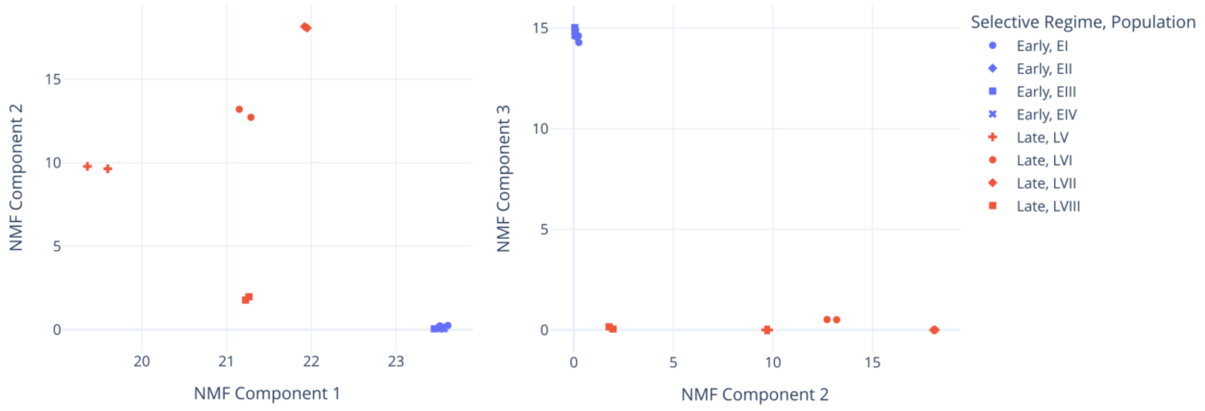

**Fig. S9.** Non-Negative Matrix Factorization (NMF) ordination of WGS read-depth profiles across samples and populations. Ordination shows component loadings from the NMF of normalized read depth data. Populations clearly cluster by EE regime along all NMF components, particularly component 1 and 3, illustrating reproducible differences in genome-wide coverage patterns across lines from the same EE regime and very high agreement between replicates from the same EE lines.

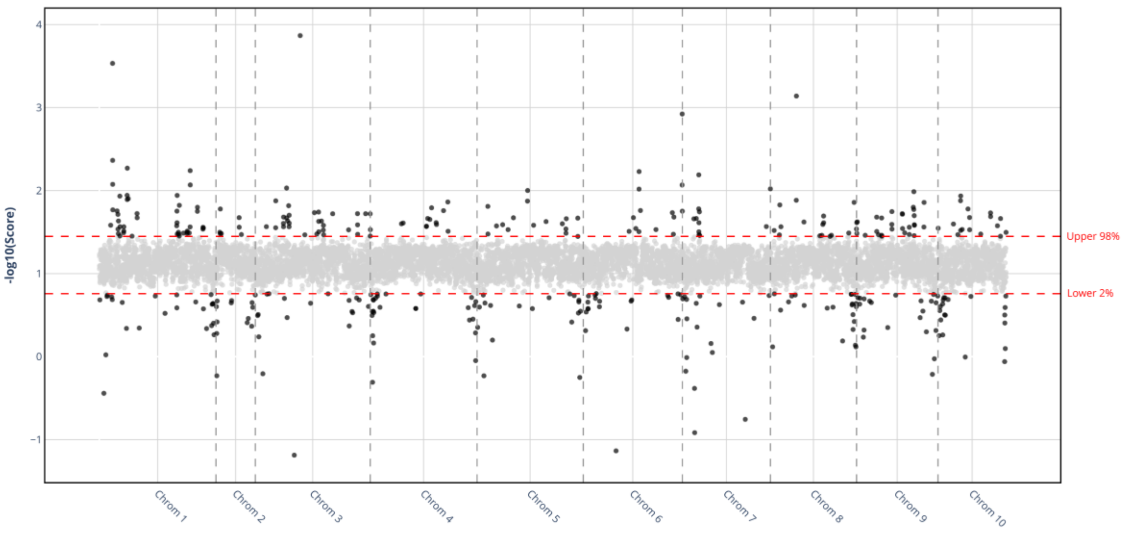

**Fig. S10.** Genomic divergence between E and L lines, based on NMF component 3 weights. Manhattan plot shows the score per 100 kb genomic bin where higher absolute scores indicate greater contribution to regime-specific divergence.

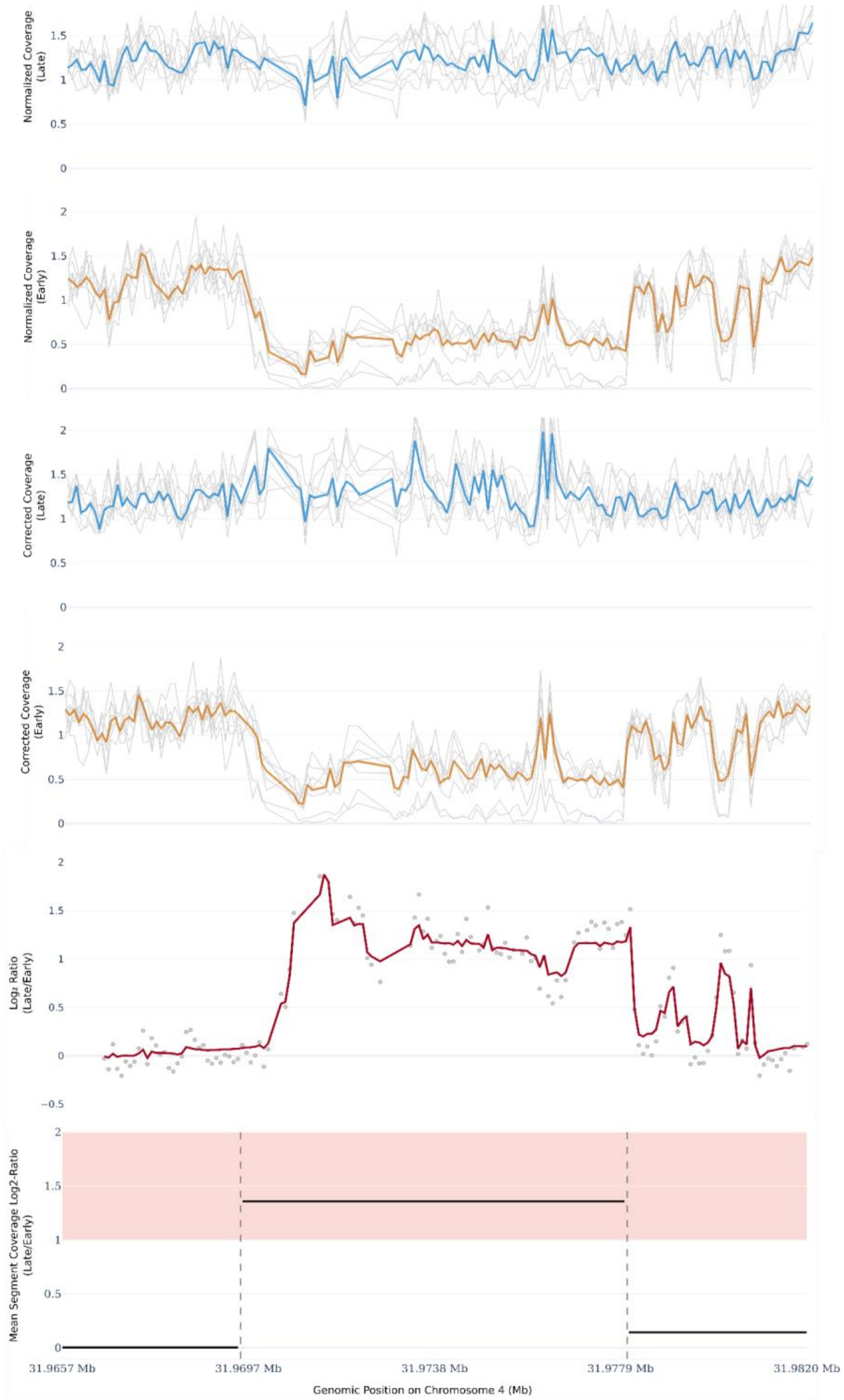

**Fig. S11.** An example of the stepwise CNV calling pipeline. Top panels show per-bin coverage profiles for all replicates in each EE regime (light grey), with regime-specific medians (blue: E; orange: L). Subsequent panels display GC- and mappability-normalized coverage signals, followed by the $log_2$-ratio signal (light grey) and its denoised version via MODWT (red line). The bottom panel shows the final CNV call at the population level, with segments exceeding a $log_2$ fold-change of ±1 highlighted.

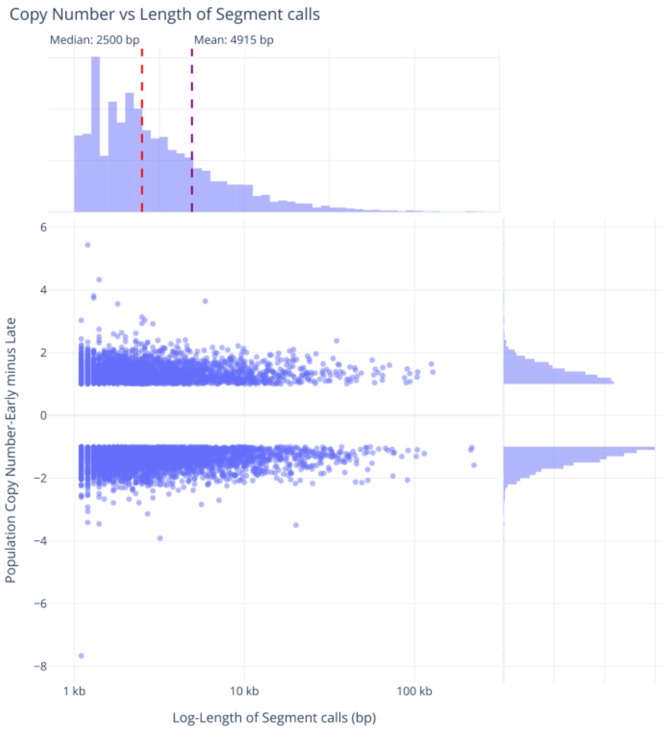

**Fig. S12**. Copy number state differentiation versus segment length, after filtering. Scatter plot show EE regime-level copy number differences as a function of log-transformed segment length. Marginal histograms illustrate the distributions of each variable.

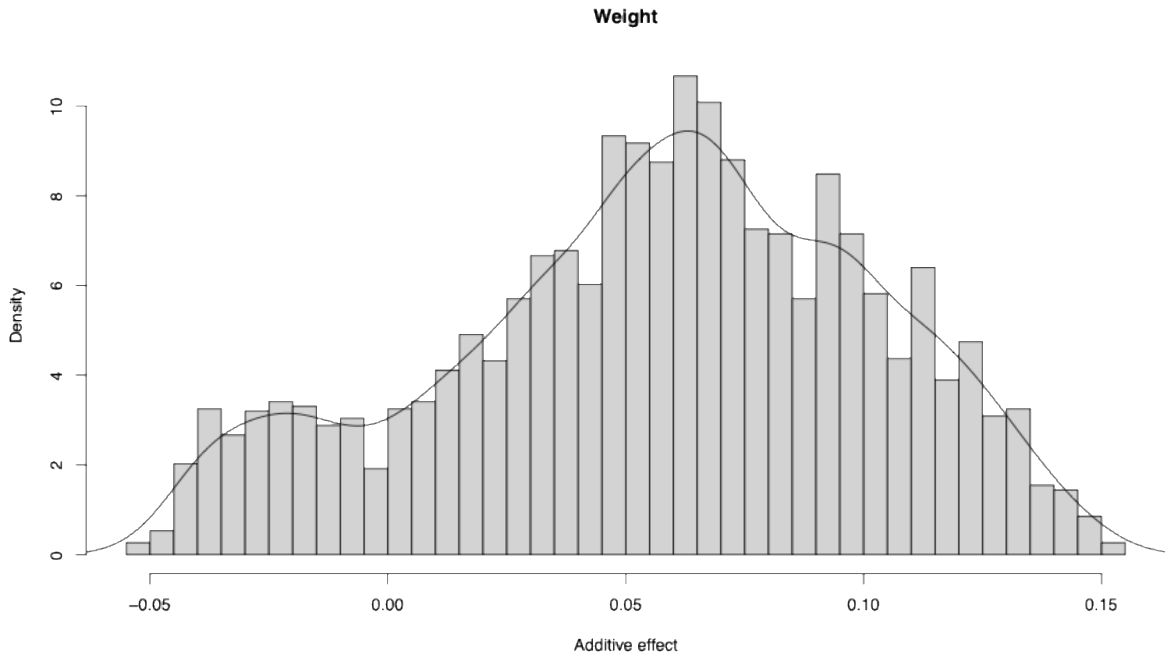

***Fig. S13.*** *Distribution (histogram and density curve) of the sex-averaged additive effect of the L allele for bodyweight, across all QTL markers.*

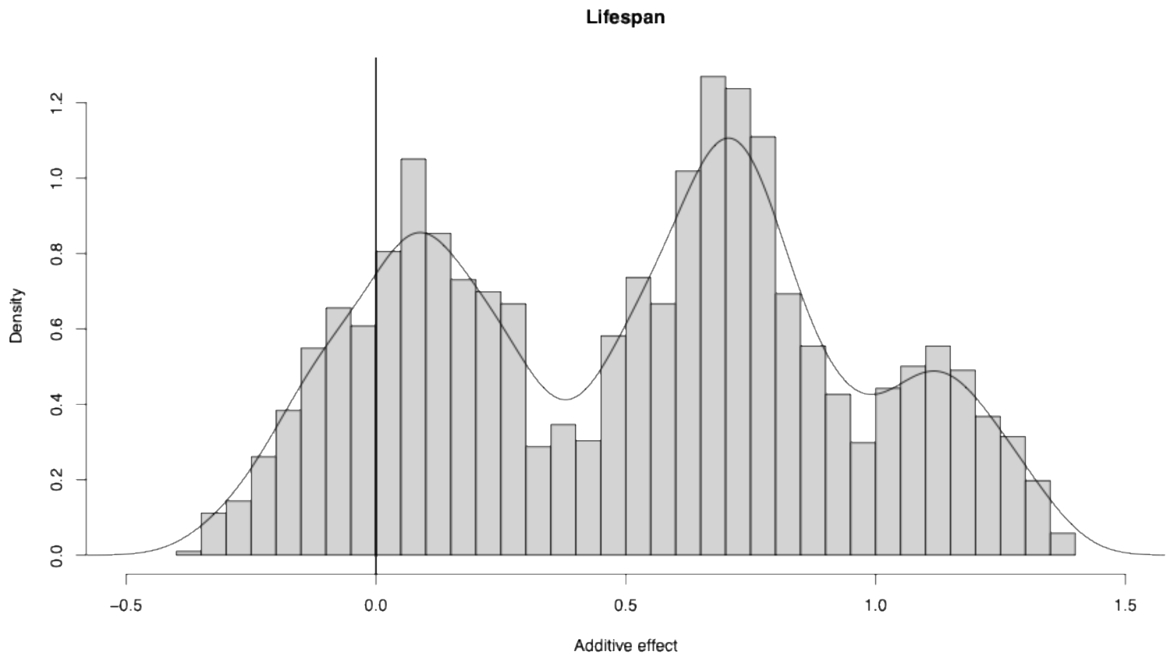

***Fig. S14****. Distribution (histogram and density curve) of the sex-averaged additive effect of the L allele for lifespan, across all QTL markers.*

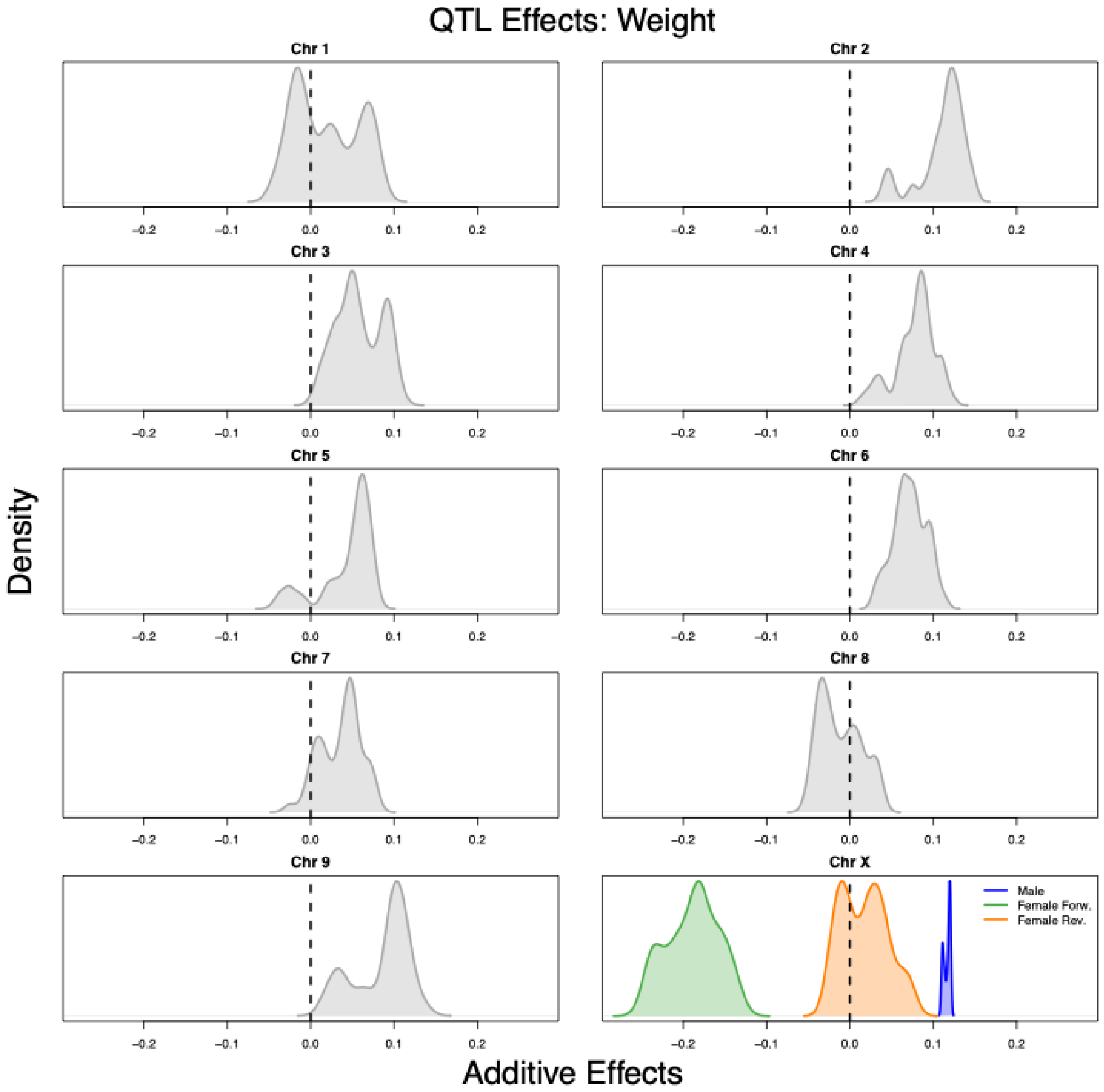

***Fig. S15***. *Distributions of per-chromosome QTL effects of the L allele for bodyweight (mg) (main effect of sex accounted for). Vertical dashed lines denote no additive effect. The X chromosome is split into estimates for males, estimates for females who received both* $X_E$ *chromosomes (Forward), and estimates for females who received both* $X_L$ *chromosomes (Reverse).*

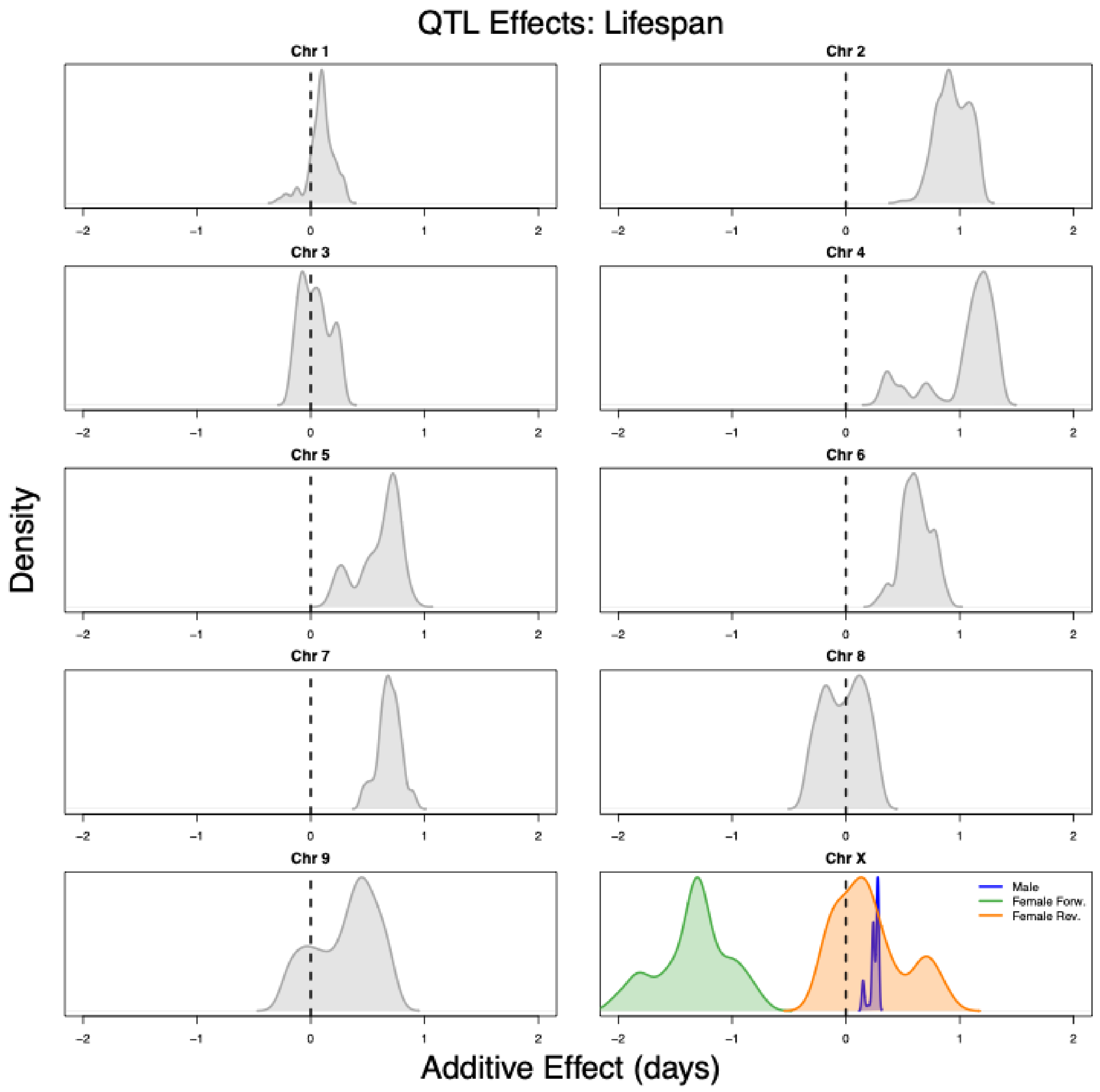

***Fig. S16.*** *Distributions of per-chromosome QTL effects of the L allele for lifespan (days) (main effect of sex accounted for). Vertical dashed lines denote no additive effect. The X chromosome is split into estimates for males, estimates for females who received both* $X_E$ *chromosomes (Forward), and estimates for females who received both* $X_L$ *chromosomes (Reverse).*

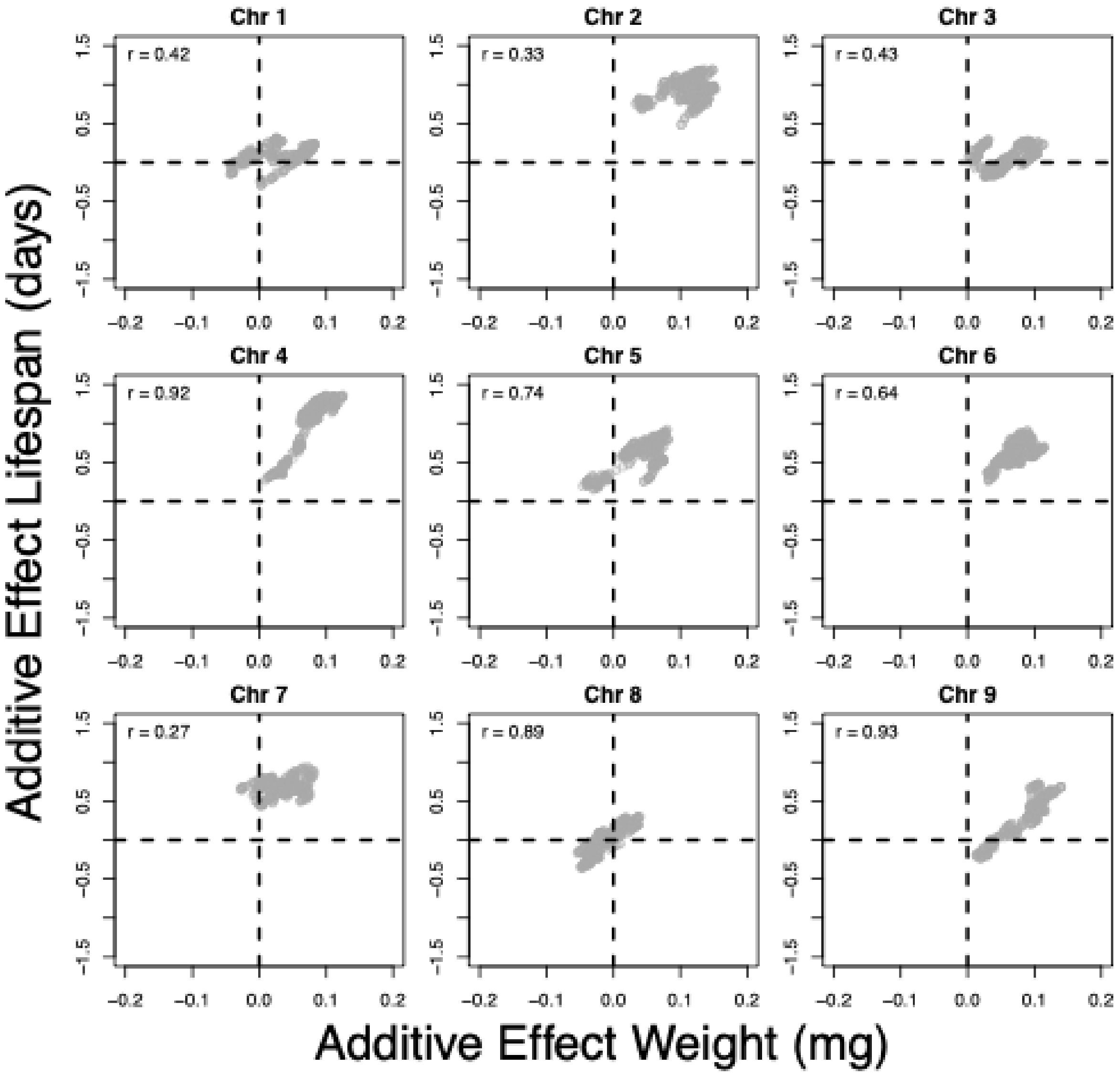

***Fig. S17****. Correlations of additive effects between weight and lifespan from sex-averaged single-QTL interval mapping.*

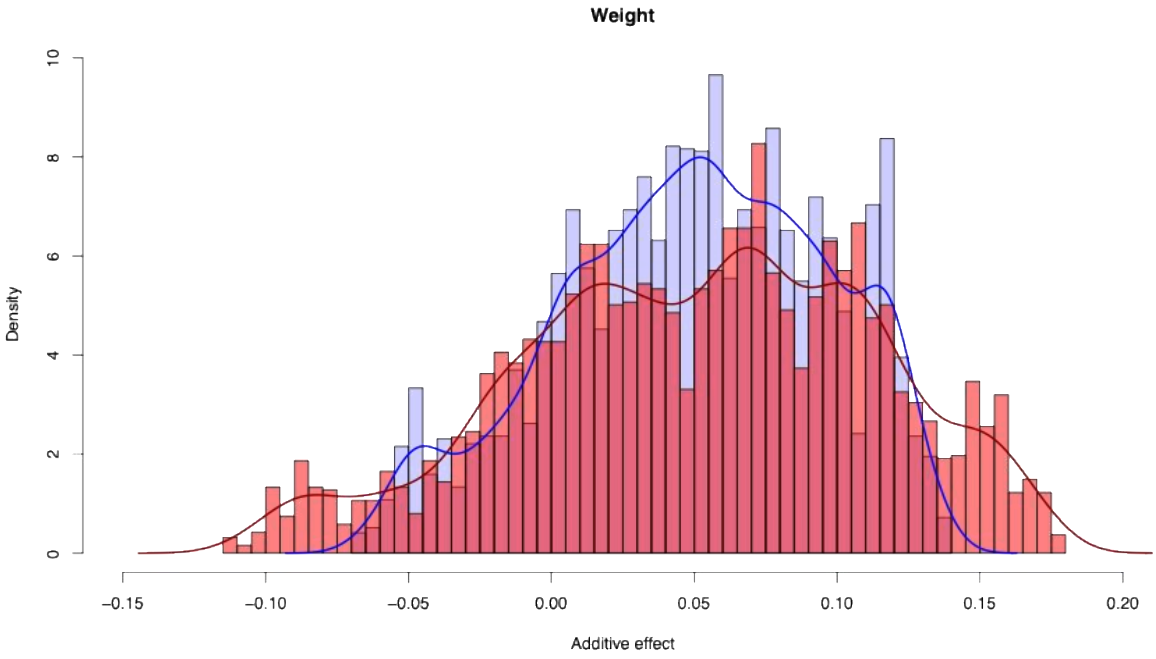

***Fig. S18**: Frequency distributions of the additive effects of the L allele for bodyweight (mg) in males (blue) and in females (red) for all QTL markers, estimated separately in the sexes in interval mapping models.*

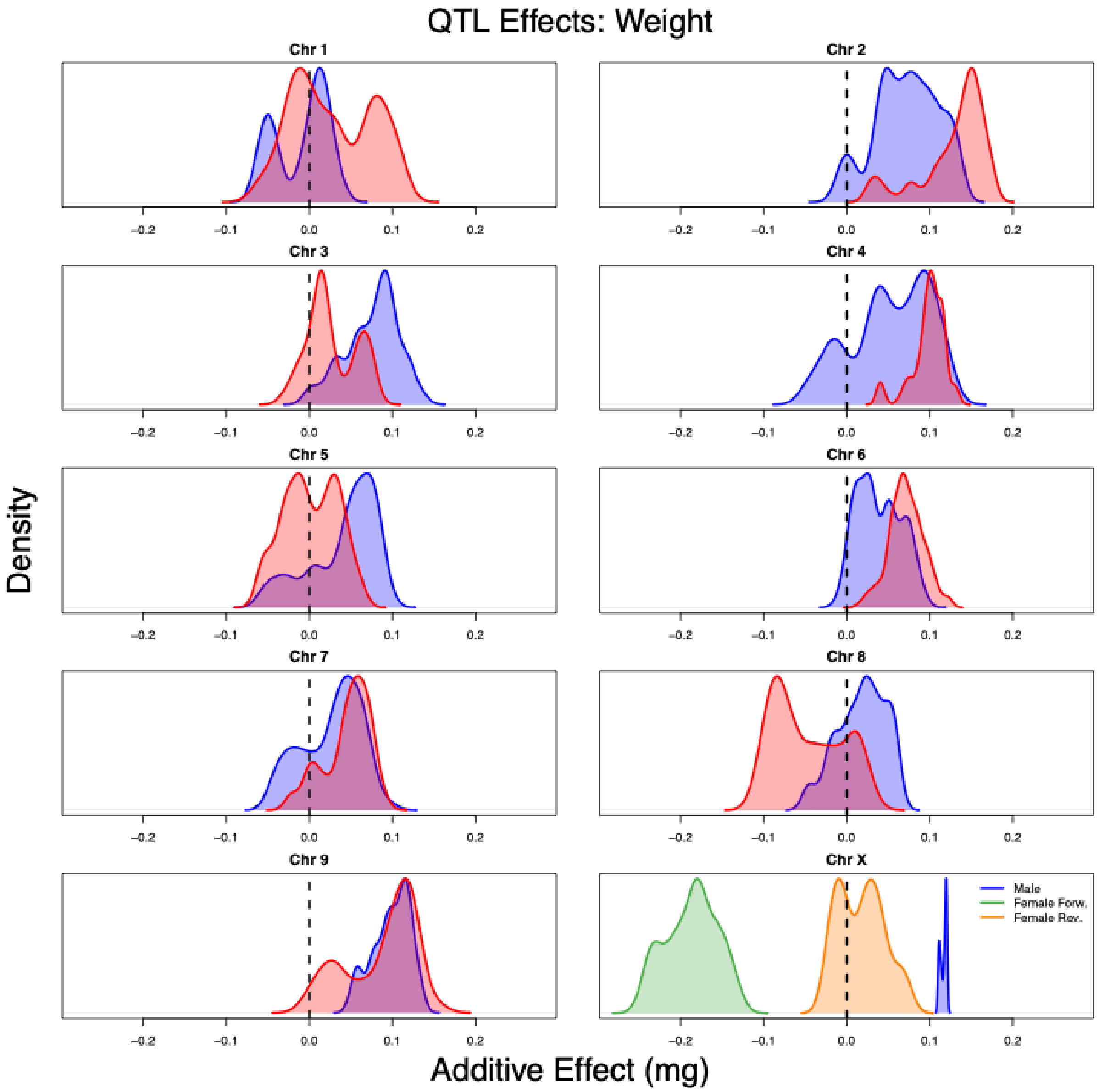

***Fig. S19**. Distributions of per-chromosome QTL effects of the L allele for weight (mg) for males (blue) and females (red) separately. Vertical dashed lines denote no additive effect. The X chromosome is split into estimates for males, estimates for females who received both $X_E$ chromosomes (Forward), and estimates for females who received both $X_L$ chromosomes (Reverse).*

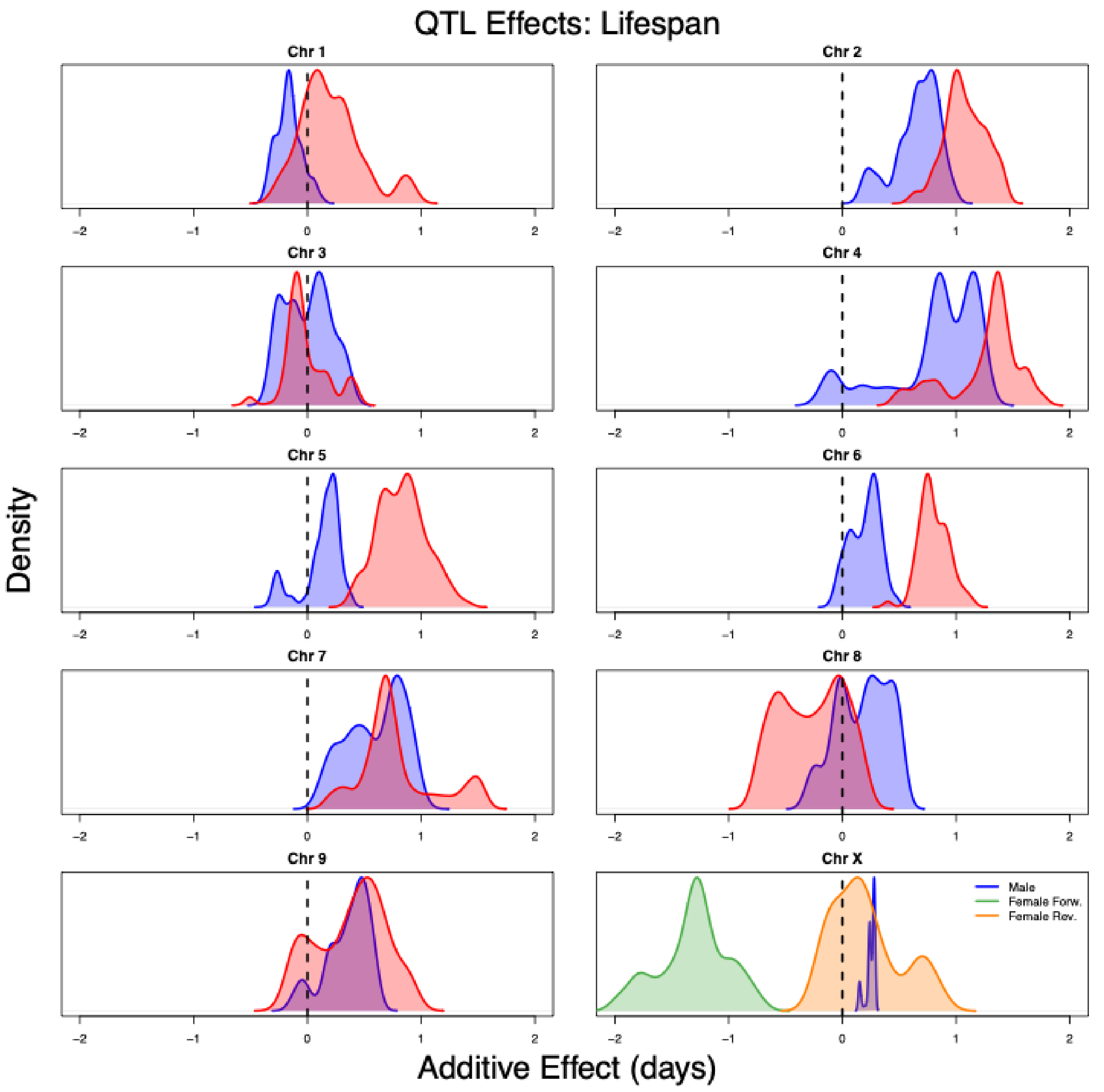

***Fig. S20***. *Distributions of per-chromosome QTL effects of the L allele for lifespan (days) for males (blue) and females (red) separately. Vertical dashed lines denote no additive effect. The X chromosome is split into estimates for males, estimates for females who received both* $X_E$ *chromosomes (Forward), and estimates for females who received both* $X_L$ *chromosomes (Reverse).*

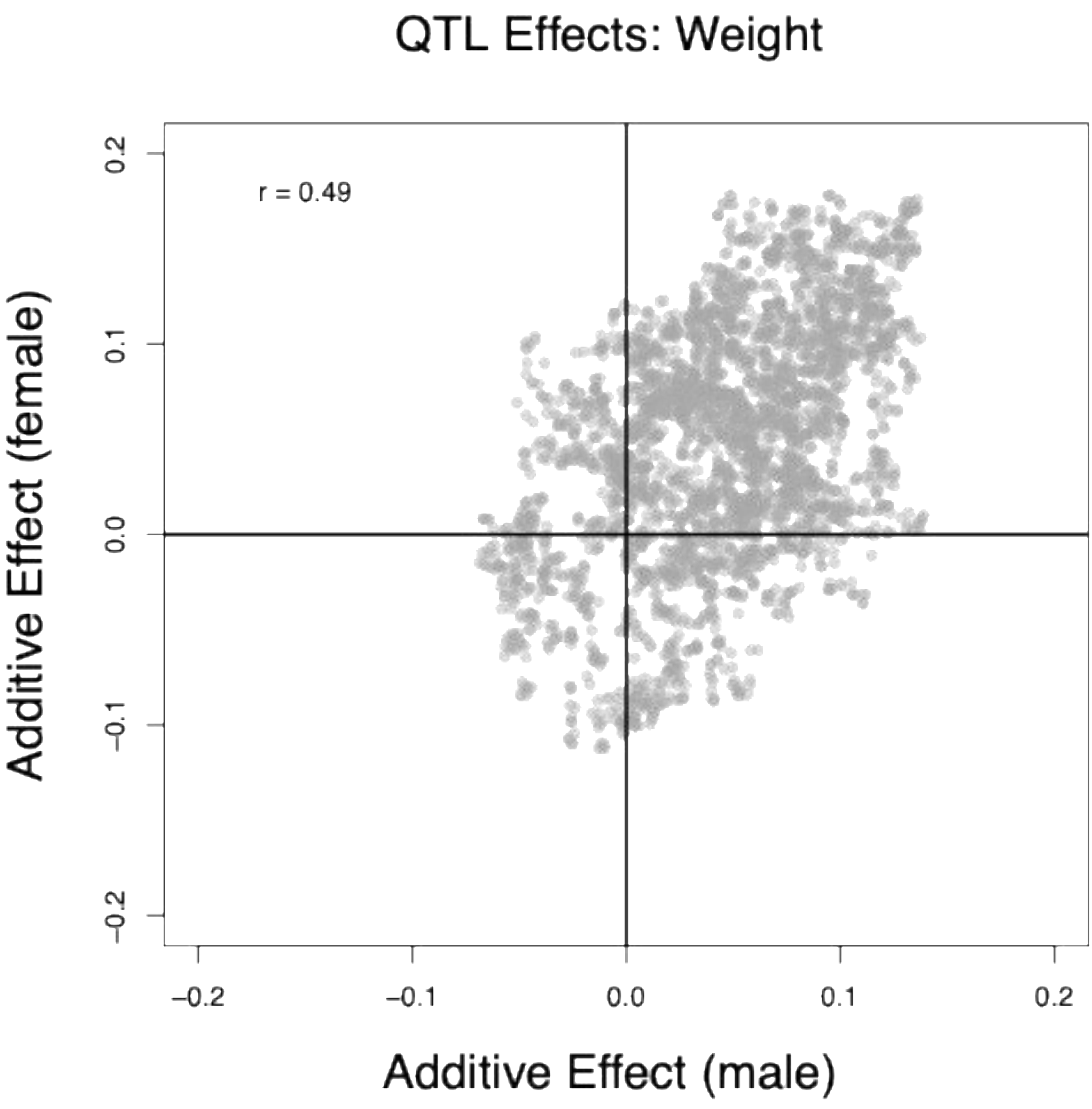

***Fig. S21***. *Genome-wide correlation between additive effects of all markers, estimates in males and females separately for bodyweight.*

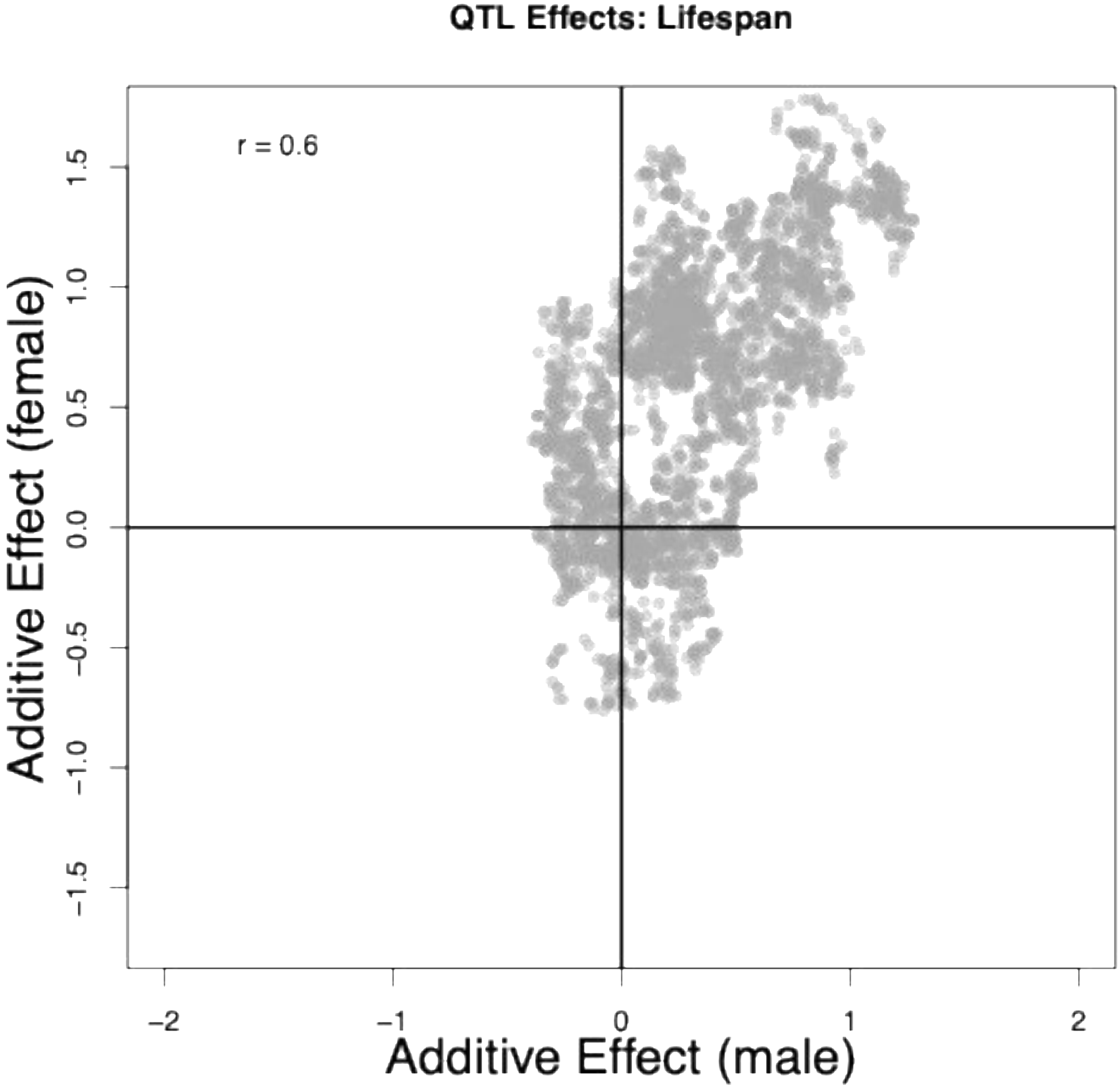

***Fig. S22***. *Genome-wide correlation between additive effects of all markers, estimates in males and females separately for lifespan.*

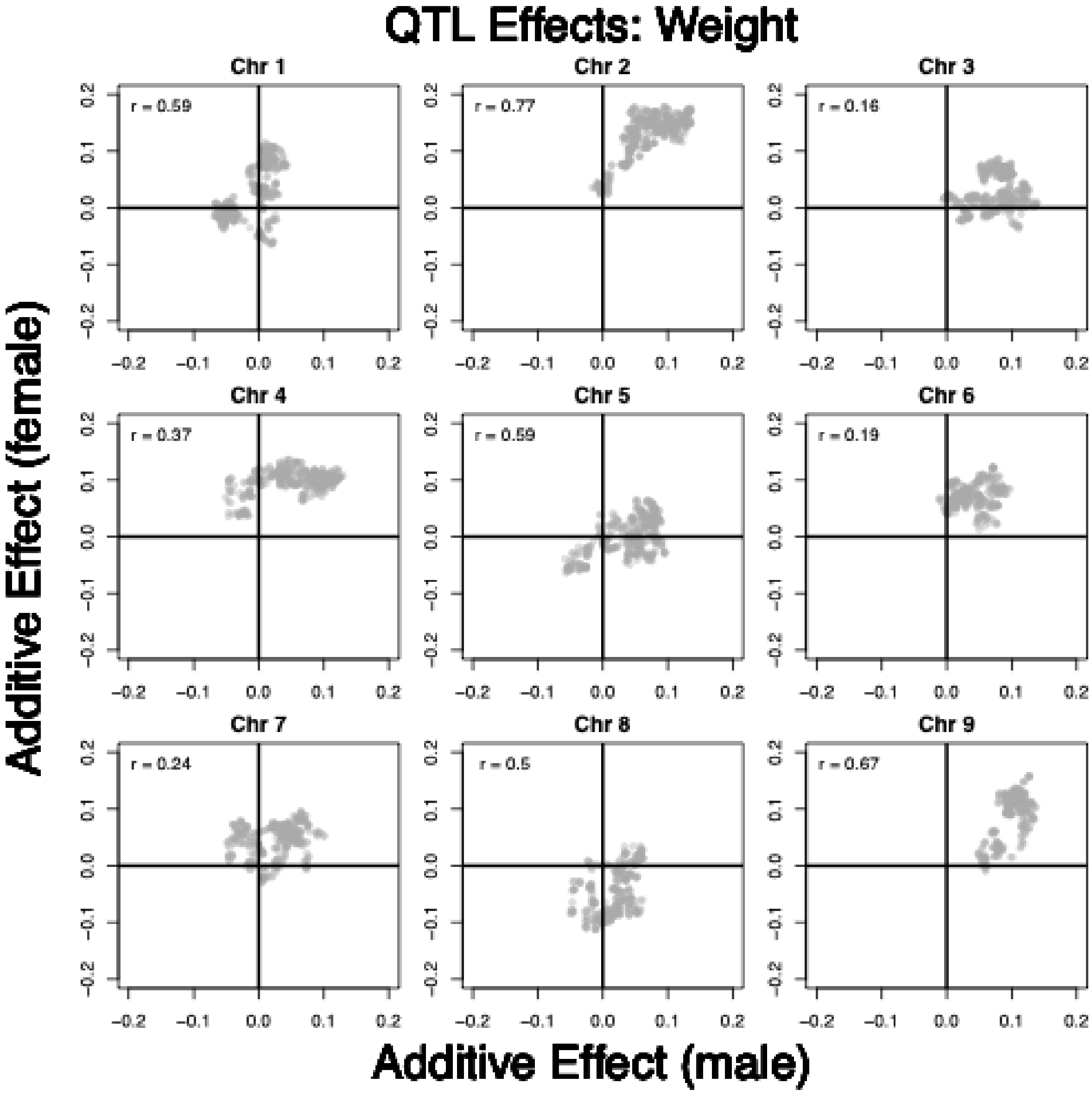

***Fig. S23***. *Per-chromosome correlation between additive effect estimates in males and females separately for bodyweight.*

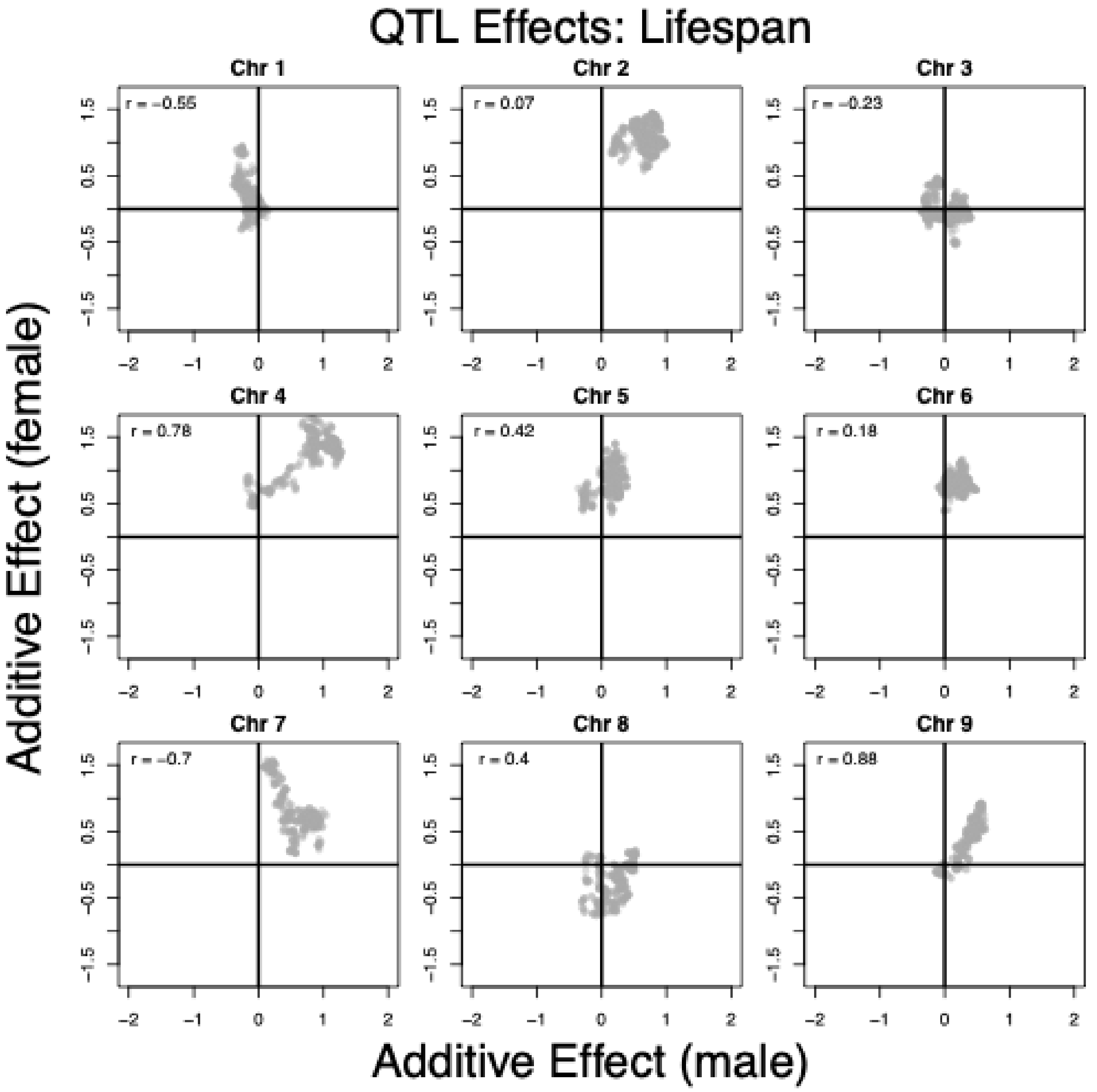

***Fig. S24****. Per-chromosome correlation between additive effect estimates in males and females separately for lifespan.*

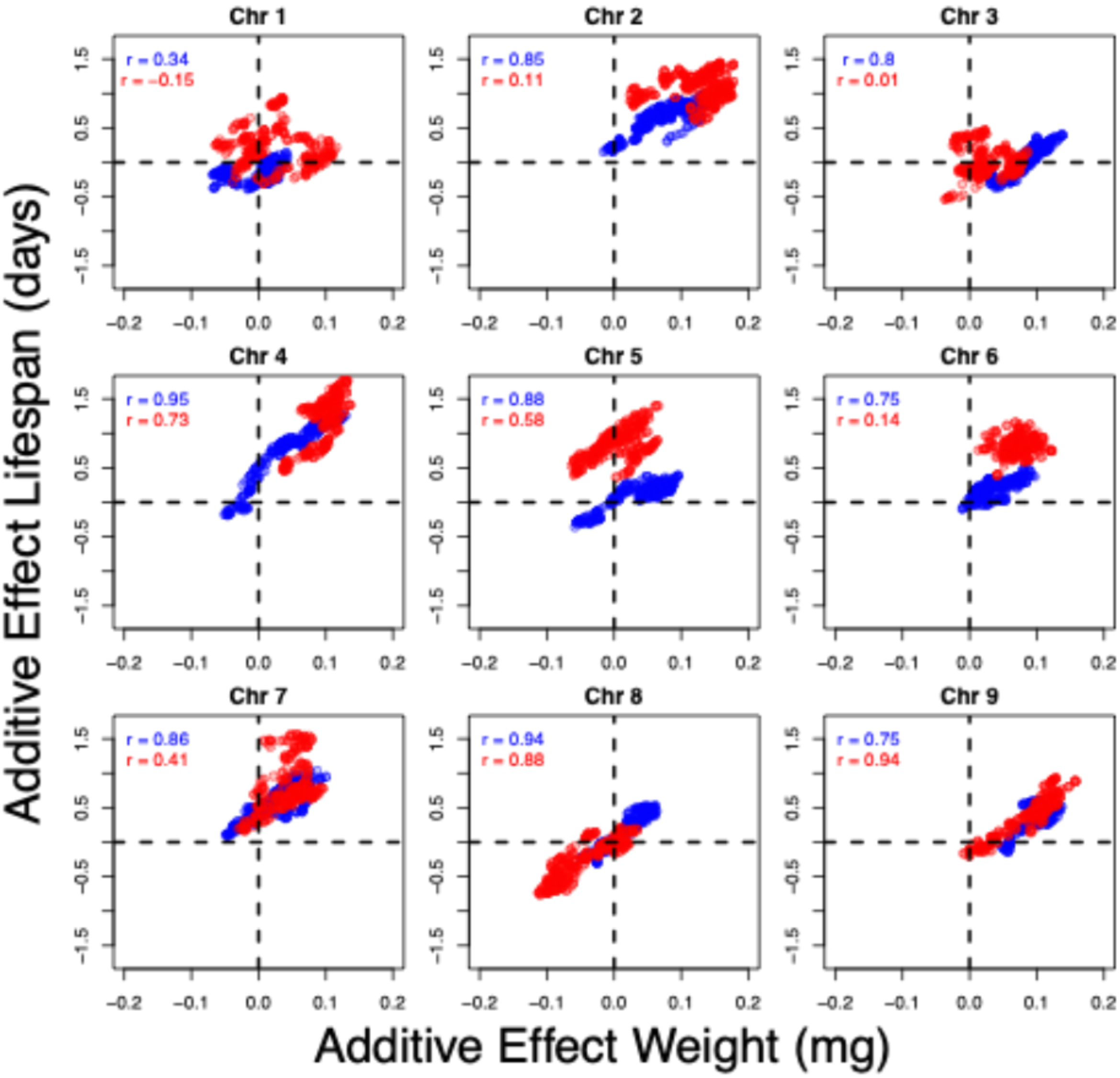

***Fig. S25****. Per-chromosome correlations between the additive effects of weight and lifespan for males (blue) and females (red) separately across markers.*

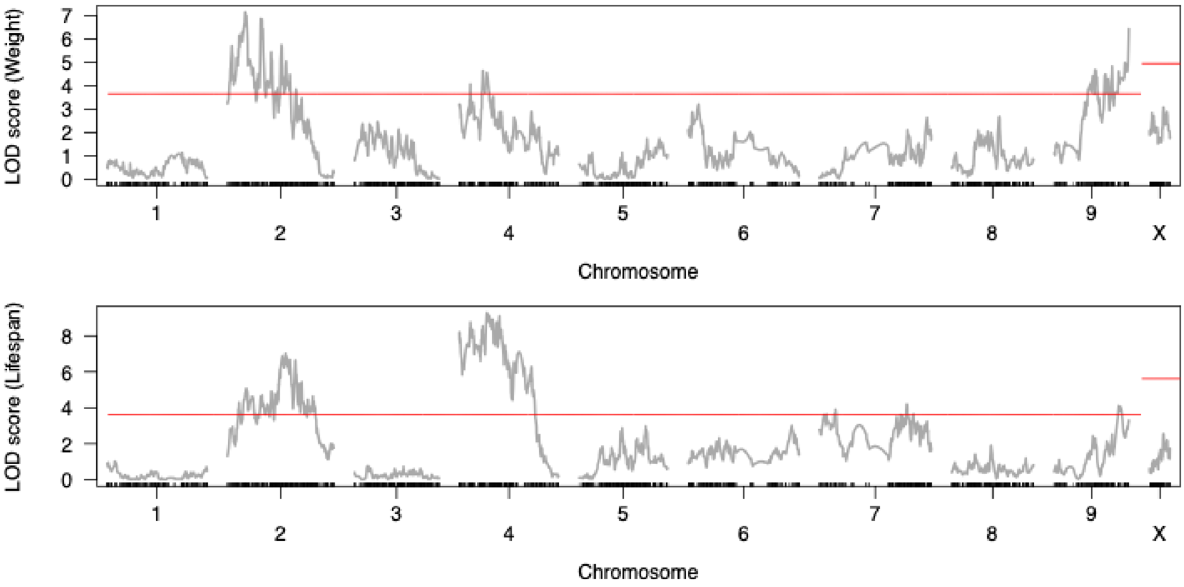

***Fig. S26***. *LOD scores for bodyweight and lifespan for each chromosome (sex-averaged). The red line denotes the LOD threshold determined by permutation tests, with thresholds for the autosomes and the X-chromosome estimated independently. For body weight, the greatest LOD peaks per chromosome (chromosome:window_start–window_end) occurred at 2:13,846,184–13,946,184, 4:14,405,940–14,505,940, and 9:73,803,912–73,903,912. For lifespan, the largest peaks were located at 2:56,724,798–56,824,798; 4:17,105,940–17,205,940; 9:57,686,282–57,786,282, but also at 7:52,022,540–52,122,540.*

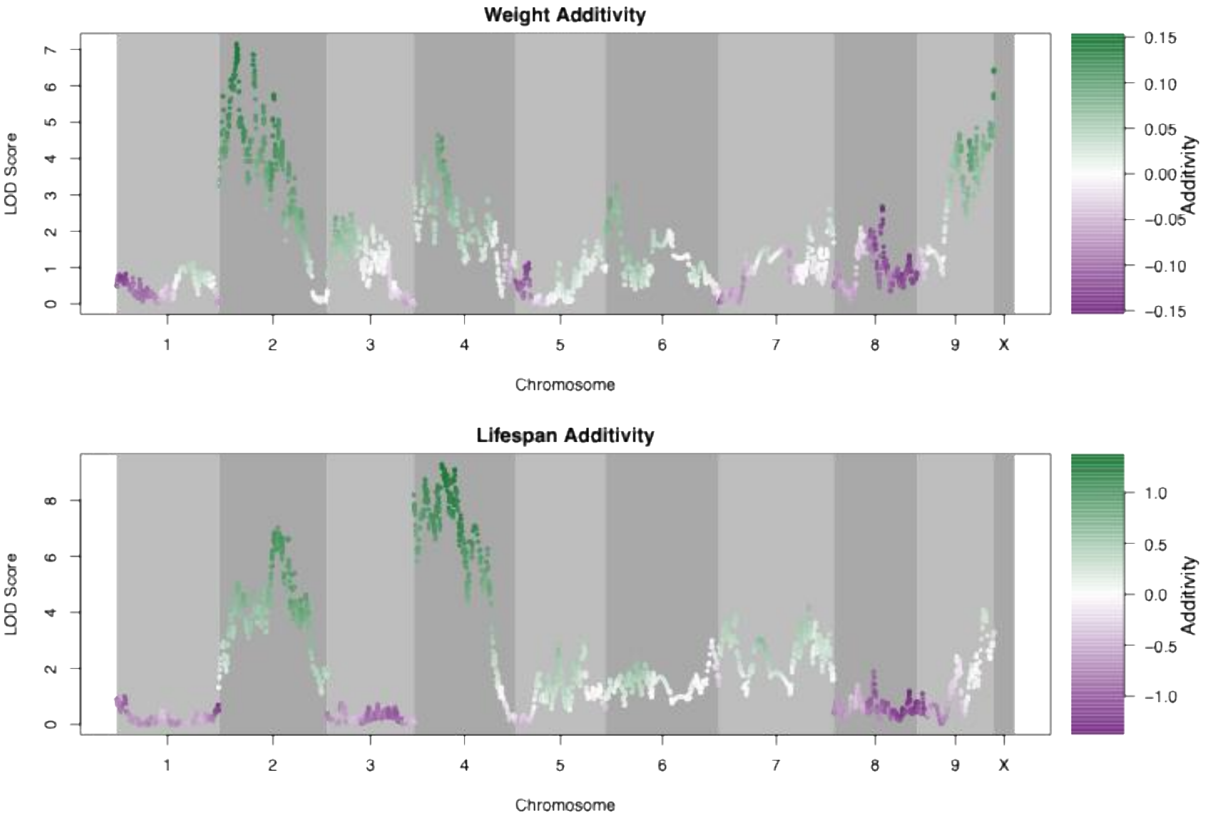

***Fig. S27***. *LOD scores for bodyweight and lifespan for each chromosome colored by the additive effect of the L allele for bodyweight (mg) and lifespan (days).*

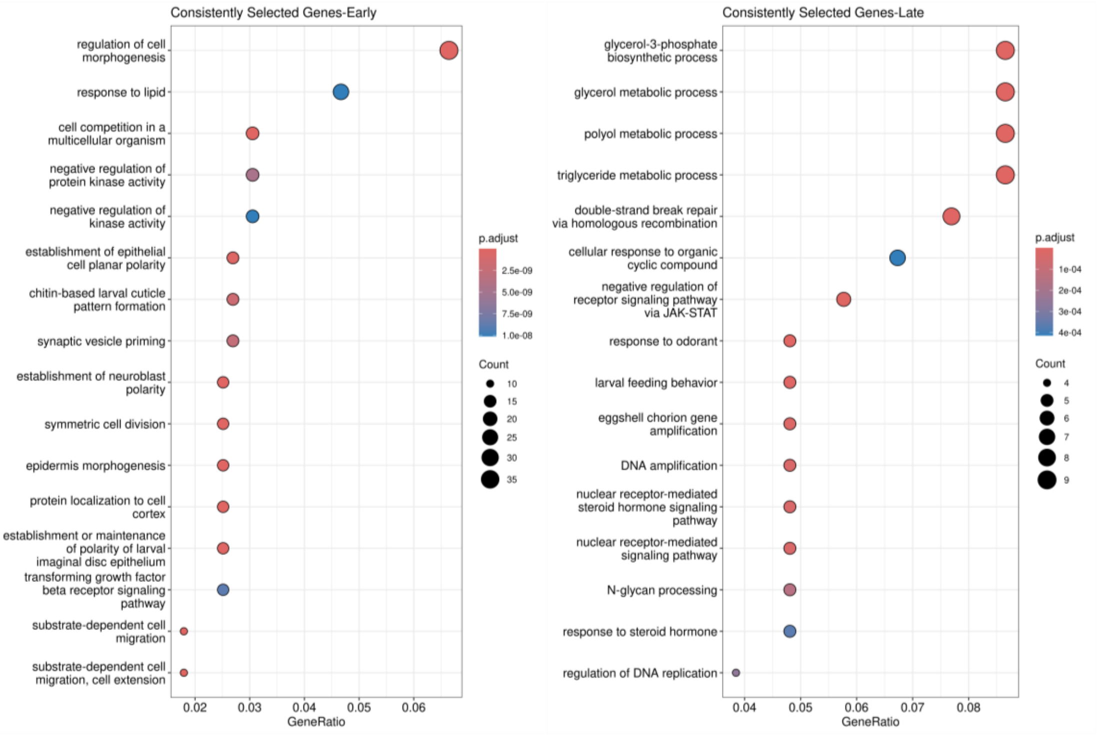

**Fig. S28.** Top-ranked enriched Gene Ontology terms (Biological Process) for genes consistently showing selective sweeps in E lines (left) and L lines (right). Terms are ranked by the proportion of unique input genes annotated with each GO term (GeneRatio).

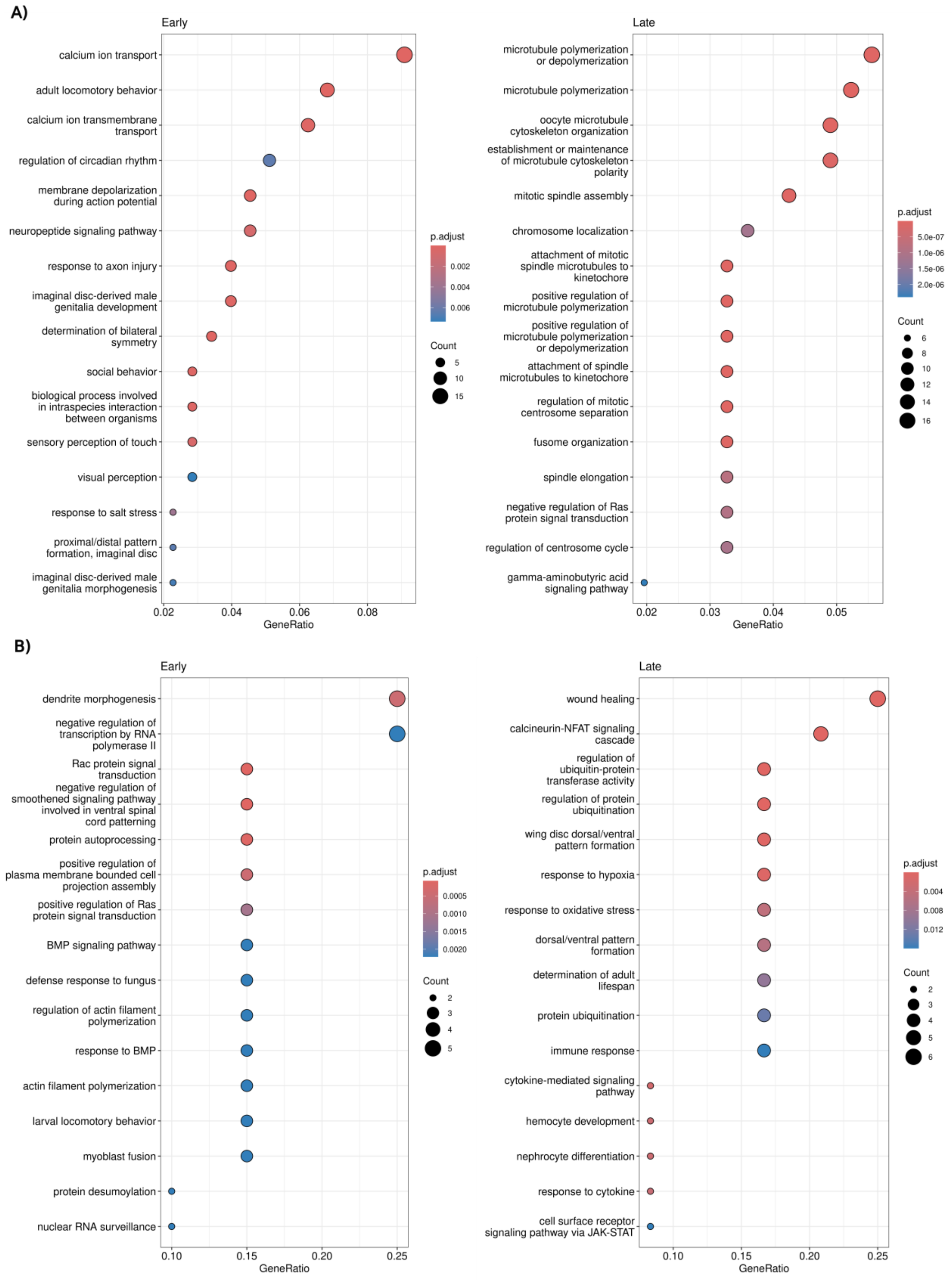

**Fig. S29**. Top-ranked enriched Gene Ontology terms (Biological Process) for genes enclosed within CNV regions. Shows are terms for (A) deletions and (B) gains in EE lines (left) and L lines (right), based on overrepresentation analyses of the final CNV call set. Terms are ranked by the proportion of unique input genes annotated with each GO term (GeneRatio).

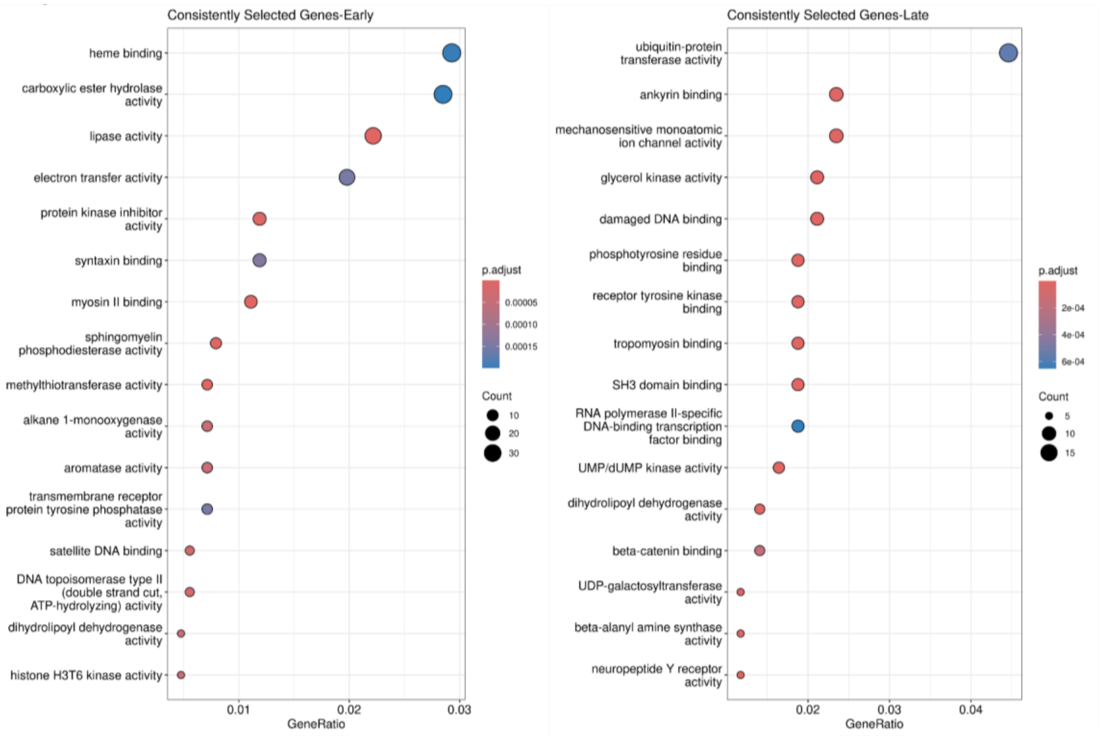

**Fig. S30.** Top-ranked Gene Ontology terms (Molecular Function) for genes consistently showing selective sweeps in E lines (left) and L lines (right). Terms are ranked by the proportion of unique input genes annotated with each GO term (GeneRatio).

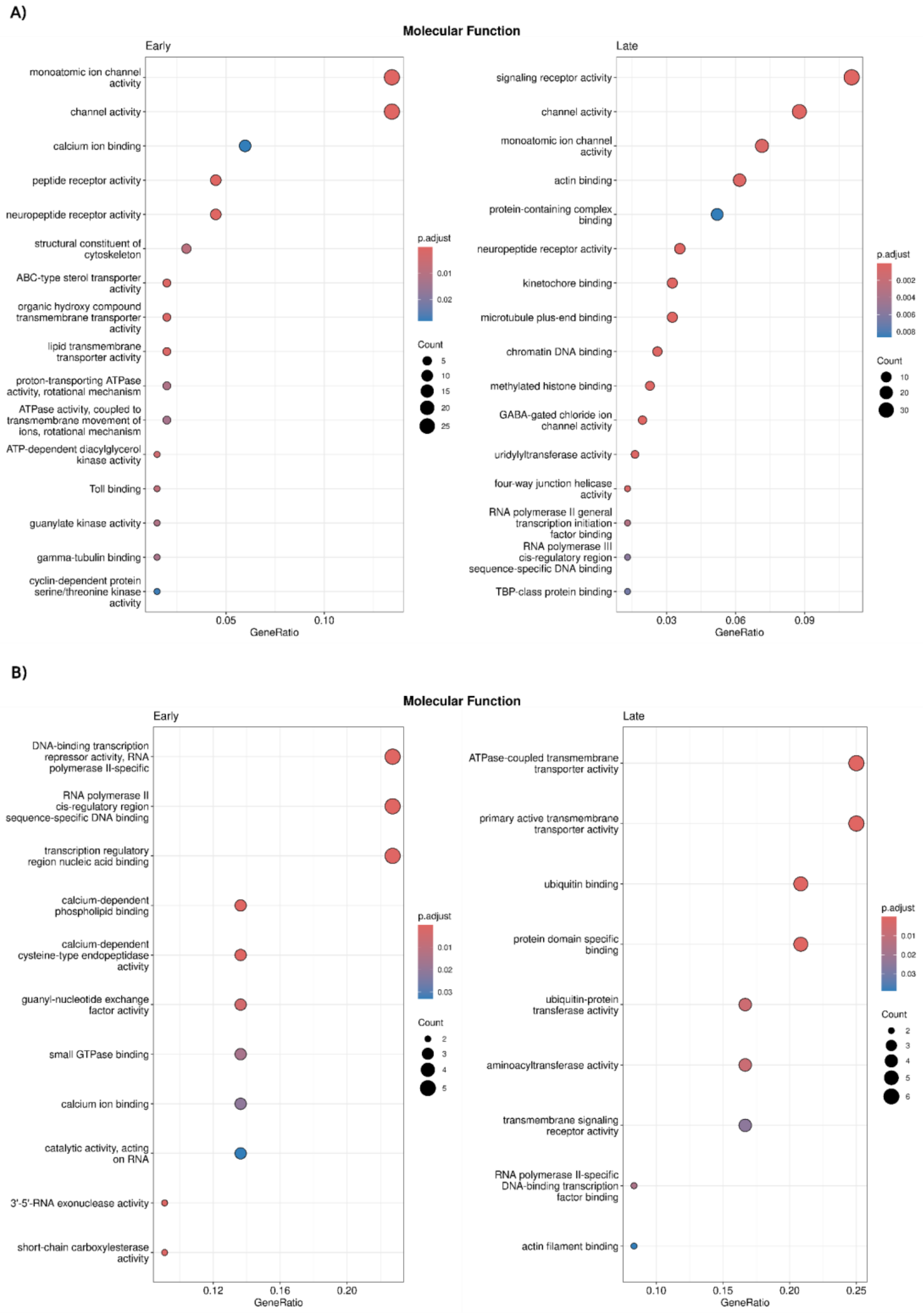

**Fig. S31**. Top-ranked enriched Gene Ontology terms (Molecular Function) for genes enclosed within CNV regions. Shows are terms for (A) deletions and (B) gains in E lines (left) and L lines (right), based on overrepresentation analyses of the final CNV call set. Terms are ranked by the proportion of unique input genes annotated with each GO term (GeneRatio).

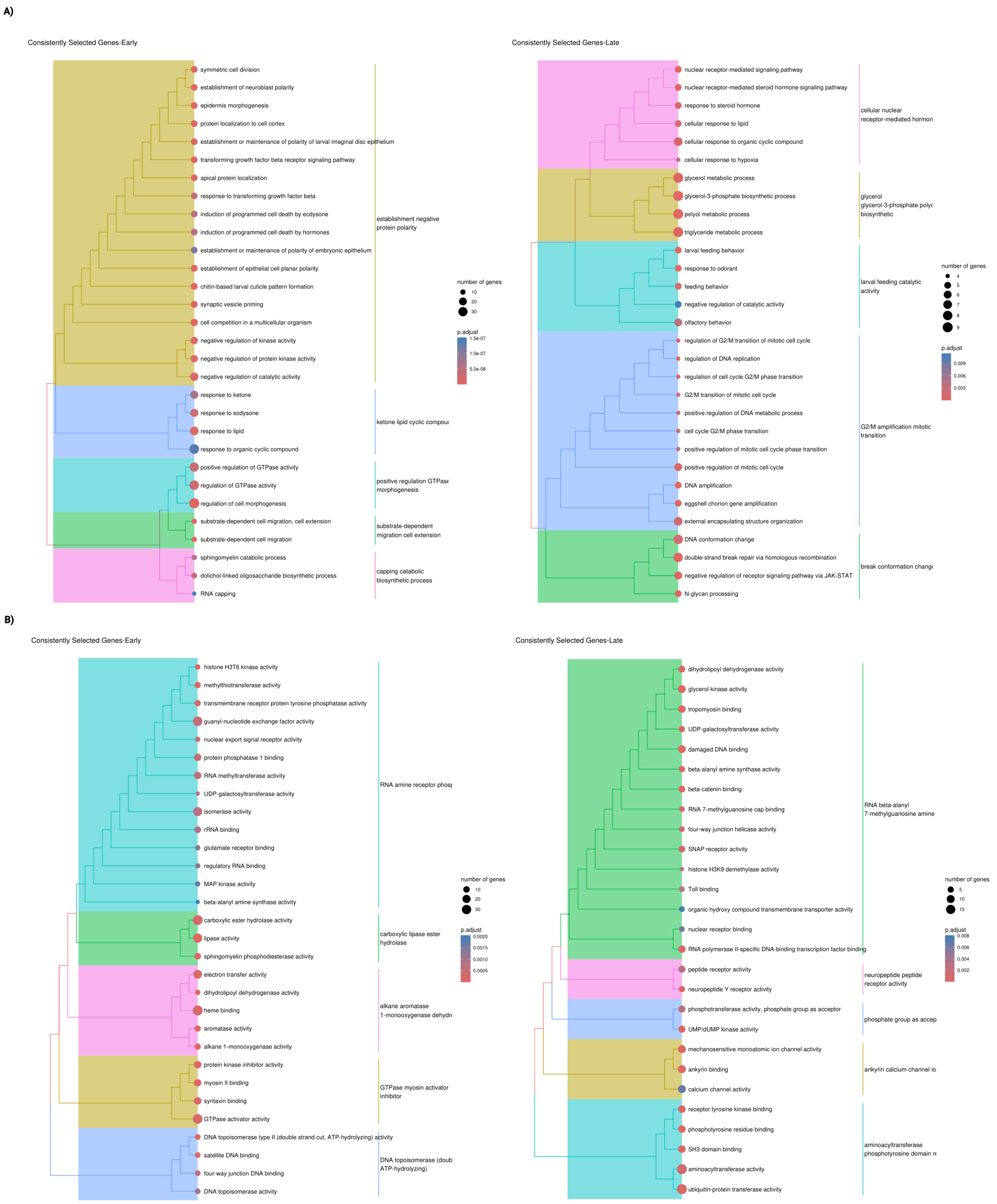

**Fig. S32**. Tree plot of enriched GO terms among genes consistently showing signs of selective sweeps for (A) Biological Process and (B) Molecular Function terms, in E lines (left) and L lines (right). Clustering by semantic similarity using the Wang method.

**A)**

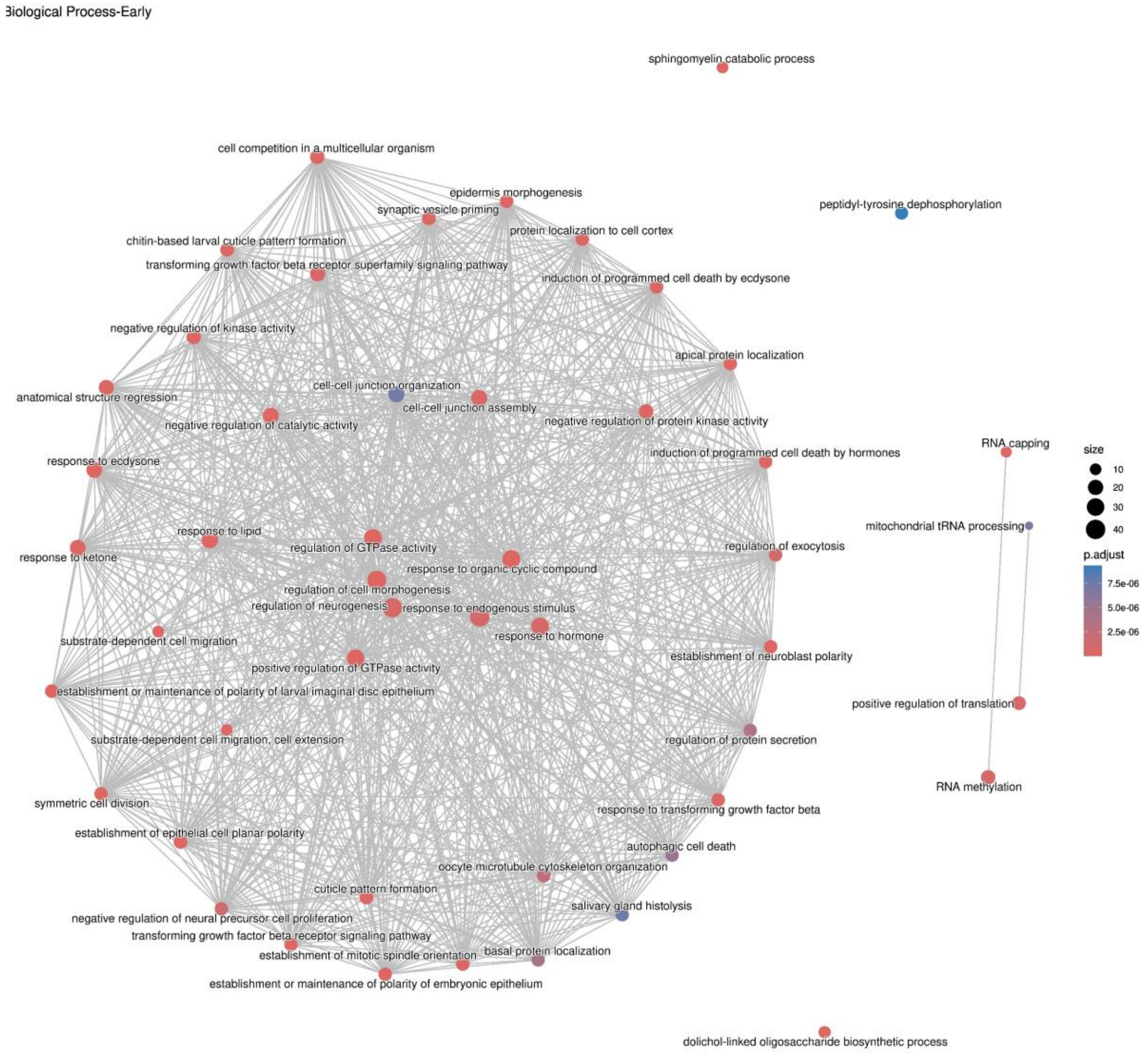

Biological Process-Early
sphingomyelin catabolic process
cell competition in a multicellular organism
epidermis morphogenesis
synaptic vesicle priming
peptidyl-tyrosine dephosphorylation
protein localization to cell cortex
chitin-based larval cuticle pattern formation
transforming growth factor beta receptor superfamily signaling pathway
induction of programmed cell death by ecdysone
negative regulation of kinase activity
apical protein localization
cell-cell junction organization
anatomical structure regression
cell-cell junction assembly
negative regulation of catalytic activity
negative regulation of protein kinase activity
RNA capping
response to ecdysone
induction of programmed cell death by hormones
size
10
20
30
40
mitochondrial tRNA processing
response to lipid
regulation of GTPase activity
regulation of exocytosis
p.adjust
response to ketone
response to organic cyclic compound
7.5e-06
5.0e-06
regulation of cell morphogenesis
regulation of neurogenesis
response to endogenous stimulus
2.5e-06
response to hormone
substrate-dependent cell migration
establishment of neuroblast polarity
positive regulation of GTPase activity
establishment or maintenance of polarity of larval imaginal disc epithelium
positive regulation of translation
substrate-dependent cell migration, cell extension
regulation of protein secretion
RNA methylation
symmetric cell division
response to transforming growth factor beta
establishment of epithelial cell planar polarity
autophagic cell death
oocyte microtubule cytoskeleton organization
cuticle pattern formation
salivary gland histolysis
negative regulation of neural precursor cell proliferation
transforming growth factor beta receptor signaling pathway
basal protein localization
establishment of mitotic spindle orientation
establishment or maintenance of polarity of embryonic epithelium
dolichol-linked oligosaccharide biosynthetic process

B)

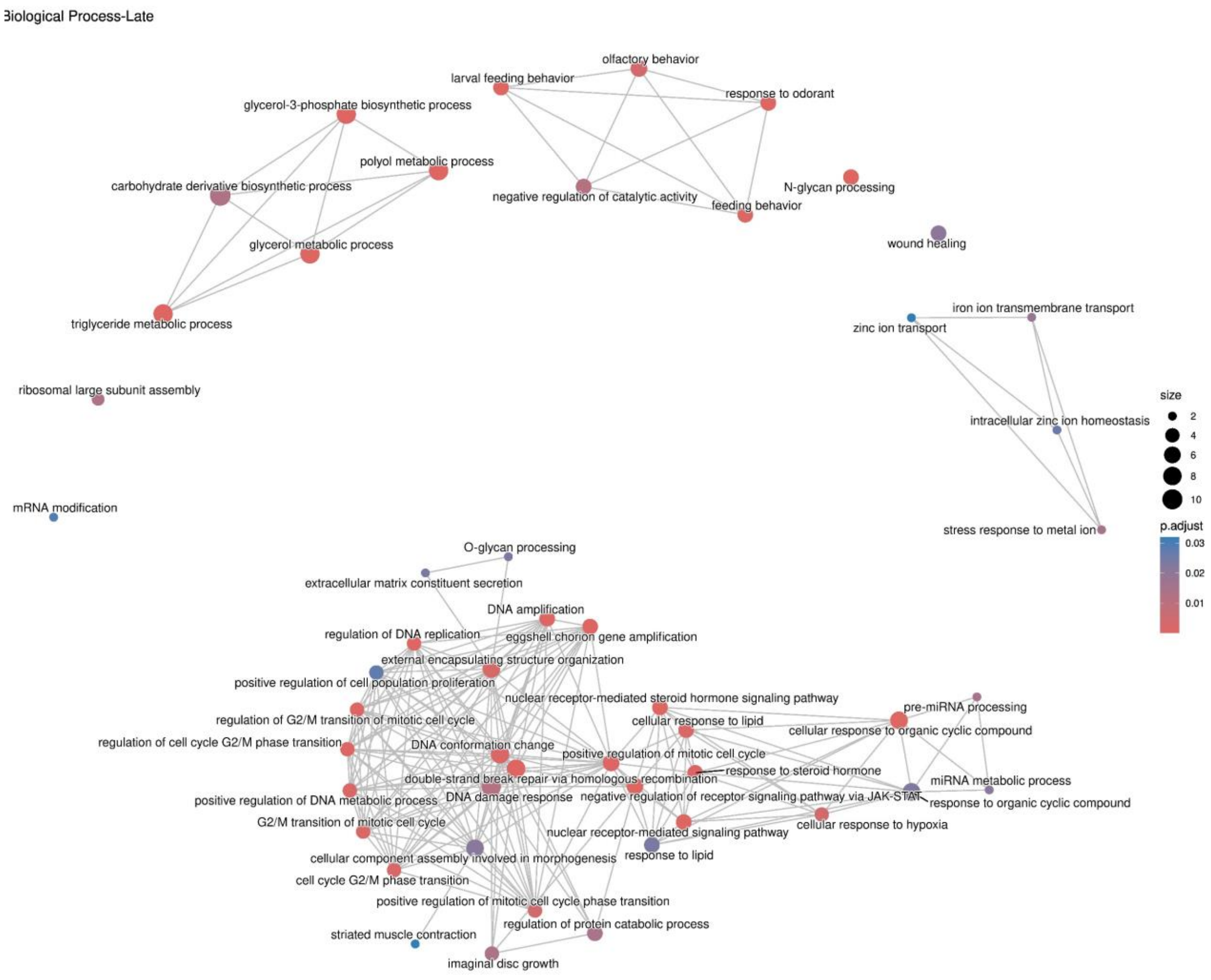

**Fig. S33**. Enrichment map of overrepresented GO Biological Process terms among genes located within selective sweep regions. Top 50 significantly enriched terms for (A) Early and (B) Late lines. Edges indicate ≥20% overlap in associated gene sets between terms.

A)

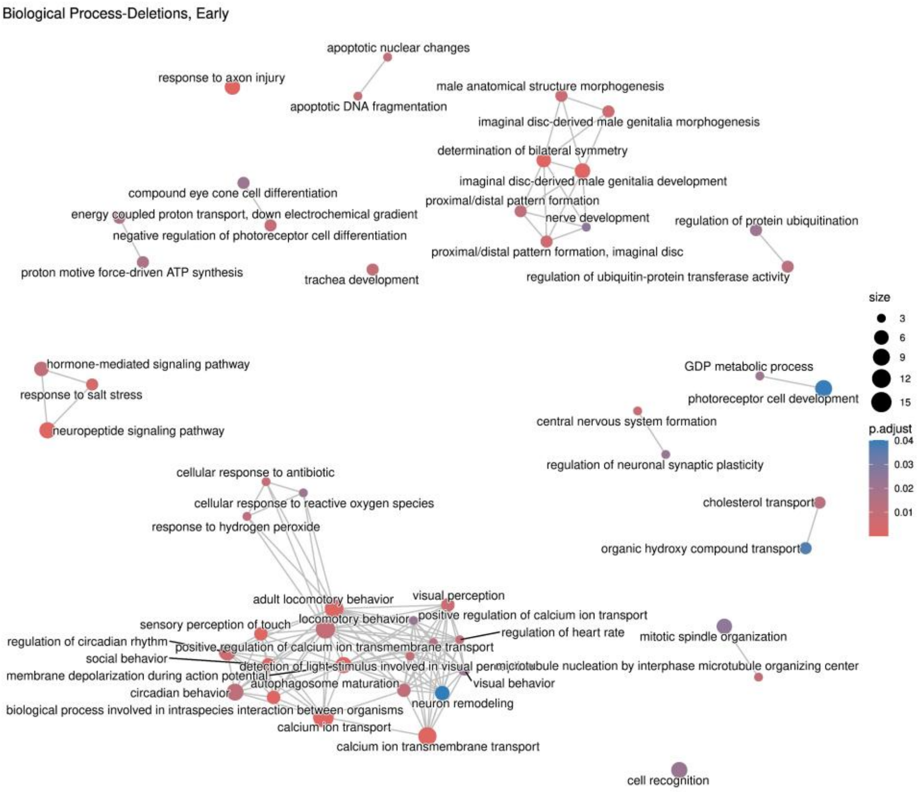
Biological Process-Deletions, Early
apoptotic nuclear changes
response to axon injury
apoptotic DNA fragmentation
male anatomical structure morphogenesis
imaginal disc-derived male genitalia morphogenesis
determination of bilateral symmetry
imaginal disc-derived male genitalia development
compound eye cone cell differentiation
energy coupled proton transport, down electrochemical gradient
negative regulation of photoreceptor cell differentiation
proximal/distal pattern formation
nerve development
regulation of protein ubiquitination
proximal/distal pattern formation, imaginal disc
proton motive force-driven ATP synthesis
trachea development
regulation of ubiquitin-protein transferase activity
size
3
6
9
12
15
hormone-mediated signaling pathway
response to salt stress
neuropeptide signaling pathway
GDP metabolic process
photoreceptor cell development
central nervous system formation
p.adjust
0.04
0.03
0.02
0.01
regulation of neuronal synaptic plasticity
cellular response to antibiotic
cellular response to reactive oxygen species
response to hydrogen peroxide
cholesterol transport
organic hydroxy compound transport
adult locomotory behavior
visual perception
sensory perception of touch
locomotory behavior
positive regulation of calcium ion transport
regulation of heart rate
regulation of circadian rhythm
positive regulation of calcium ion transmembrane transport
mitotic spindle organization
social behavior
detection of light stimulus involved in visual perception
microtubule nucleation by interphase microtubule organizing center
membrane depolarization during action potential
autophagosome maturation
visual behavior
circadian behavior
neuron remodeling
biological process involved in intraspecies interaction between organisms
calcium ion transport
calcium ion transmembrane transport
cell recognition

B)

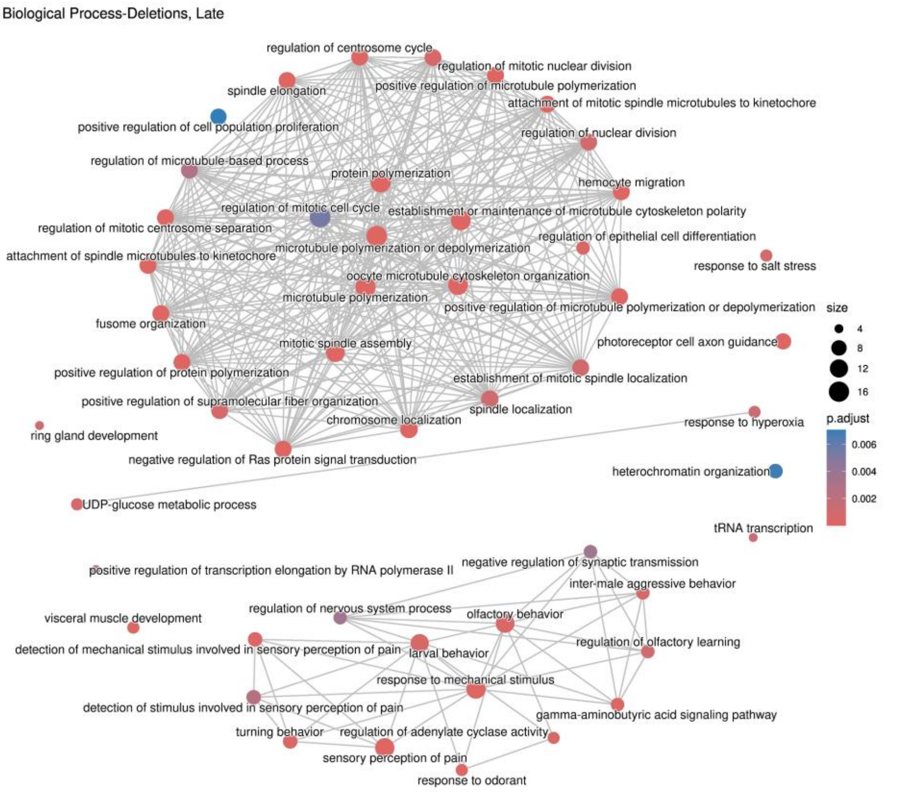
Biological Process-Deletions, Late
regulation of centrosome cycle
regulation of mitotic nuclear division
spindle elongation
positive regulation of microtubule polymerization
attachment of mitotic spindle microtubules to kinetochore
positive regulation of cell population proliferation
regulation of nuclear division
regulation of microtubule-based process
protein polymerization
hemocyte migration
regulation of mitotic cell cycle
establishment or maintenance of microtubule cytoskeleton polarity
regulation of mitotic centrosome separation
regulation of epithelial cell differentiation
attachment of spindle microtubules to kinetochore
microtubule polymerization or depolymerization
response to salt stress
oocyte microtubule cytoskeleton organization
microtubule polymerization
positive regulation of microtubule polymerization or depolymerization
size
fusome organization
4
8
12
16
photoreceptor cell axon guidance
mitotic spindle assembly
positive regulation of protein polymerization
establishment of mitotic spindle localization
positive regulation of supramolecular fiber organization
spindle localization
chromosome localization
response to hyperoxia
ring gland development
p.adjust
negative regulation of Ras protein signal transduction
0.006
heterochromatin organization
0.004
0.002
UDP-glucose metabolic process
tRNA transcription
positive regulation of transcription elongation by RNA polymerase II
negative regulation of synaptic transmission
inter-male aggressive behavior
visceral muscle development
regulation of nervous system process
olfactory behavior
regulation of olfactory learning
detection of mechanical stimulus involved in sensory perception of pain
larval behavior
response to mechanical stimulus
detection of stimulus involved in sensory perception of pain
gamma-aminobutyric acid signaling pathway
turning behavior
regulation of adenylate cyclase activity
sensory perception of pain
response to odorant

C)

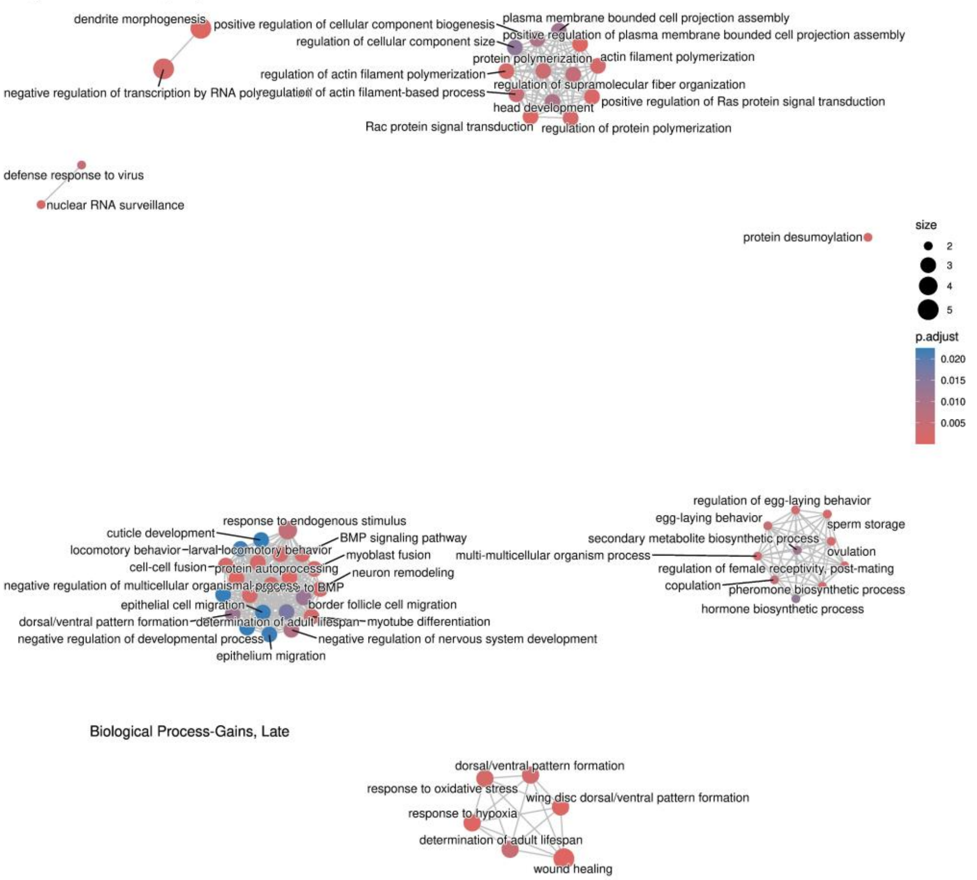

D)

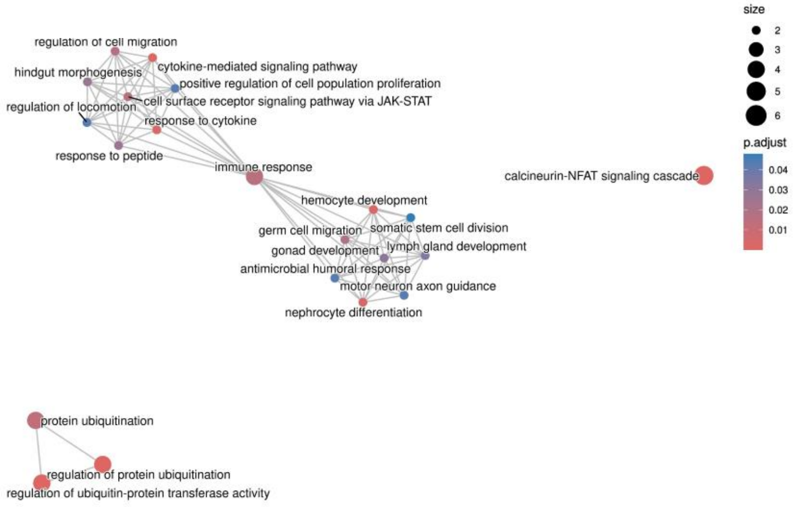

**Fig. S34.** Enrichment map of overrepresented GO Biological Process terms among genes located within CNV regions. Top 50 significantly enriched terms for genes overlapping deletions in (A) E and (B) L lines, and gains in (C) E and (D) L lines. Edges represent ≥20% overlap in associated gene sets between terms.

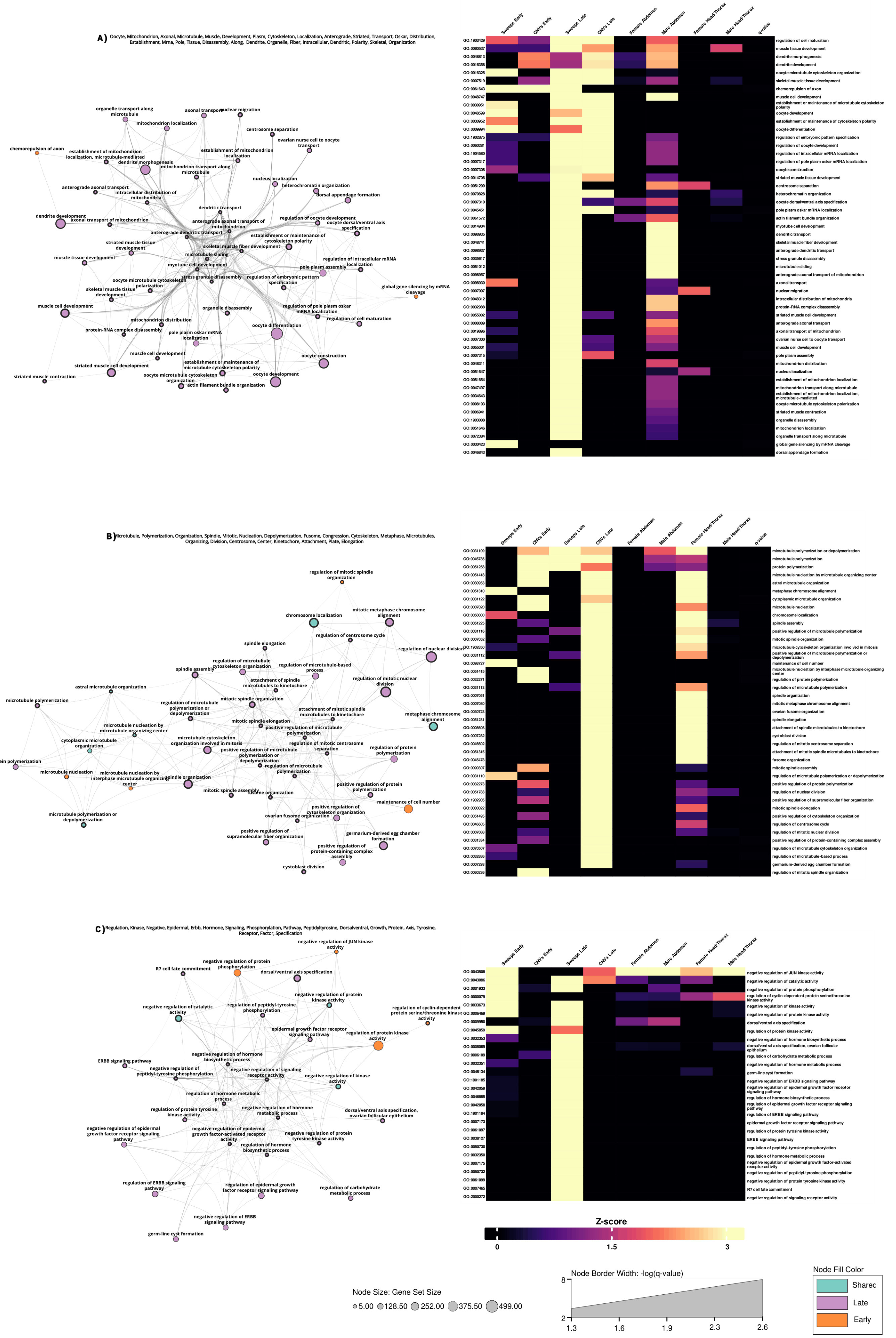
A) Oocyte, Mitochondrion, Axonal, Microtubule, Muscle, Development, Plasm, Cytoskeleton, Localization, Anterograde, Striated, Transport, Oskar, Distribution, Establishment, Mrna, Pole, Tissue, Disassembly, Along, Dendrite, Organelle, Fiber, Intracellular, Dendritic, Polarity, Skeletal, Organization
Sweeps Early
CNVs Early
Sweeps Late
CNVs Late
Female Abdomen
Male Abdomen
Female Head/Thorax
Male Head/Thorax
q-value
regulation of cell maturation
muscle tissue development
dendrite morphogenesis
dendrite development
oocyte microtubule cytoskeleton organization
skeletal muscle tissue development
chemorepulsion of axon
muscle cell development
establishment or maintenance of microtubule cytoskeleton polarity
oocyte development
establishment or maintenance of cytoskeleton polarity
oocyte differentiation
regulation of embryonic pattern specification
regulation of oocyte development
regulation of intracellular mRNA localization
regulation of pole plasm oskar mRNA localization
oocyte construction
striated muscle tissue development
centrosome separation
heterochromatin organization
oocyte dorsal/ventral axis specification
pole plasm oskar mRNA localization
actin filament bundle organization
myotube cell development
dendritic transport
skeletal muscle fiber development
anterograde dendritic transport
stress granule disassembly
microtubule sliding
anterograde axonal transport of mitochondrion
axonal transport
nuclear migration
intracellular distribution of mitochondria
protein-RNA complex disassembly
striated muscle cell development
anterograde axonal transport
axonal transport of mitochondrion
ovarian nurse cell to oocyte transport
muscle cell development
pole plasm assembly
mitochondrion distribution
nucleus localization
establishment of mitochondrion localization
mitochondrion transport along microtubule
establishment of mitochondrion localization, microtubule-mediated
oocyte microtubule cytoskeleton polarization
striated muscle contraction
organelle disassembly
mitochondrion localization
organelle transport along microtubule
global gene silencing by mRNA cleavage
dorsal appendage formation
B) Microtubule, Polymerization, Organization, Spindle, Mitotic, Nucleation, Depolymerization, Fusome, Congression, Cytoskeleton, Metaphase, Microtubules, Organizing, Division, Centrosome, Center, Kinetochore, Attachment, Plate, Elongation
microtubule polymerization or depolymerization
microtubule polymerization
protein polymerization
microtubule nucleation by microtubule organizing center
astral microtubule organization
metaphase chromosome alignment
cytoplasmic microtubule organization
microtubule nucleation
chromosome localization
spindle assembly
positive regulation of microtubule polymerization
mitotic spindle organization
microtubule cytoskeleton organization involved in mitosis
positive regulation of microtubule polymerization or depolymerization
maintenance of cell number
microtubule nucleation by interphase microtubule organizing center
regulation of protein polymerization
regulation of microtubule polymerization
spindle organization
mitotic metaphase chromosome alignment
ovarian fusome organization
spindle elongation
attachment of spindle microtubules to kinetochore
cytoblast division
regulation of mitotic centrosome separation
attachment of mitotic spindle microtubules to kinetochore
fusome organization
mitotic spindle assembly
regulation of microtubule polymerization or depolymerization
positive regulation of protein polymerization
regulation of nuclear division
positive regulation of supramolecular fiber organization
mitotic spindle elongation
positive regulation of cytoskeleton organization
regulation of centrosome cycle
regulation of mitotic nuclear division
positive regulation of protein-containing complex assembly
regulation of microtubule cytoskeleton organization
regulation of microtubule-based process
germarium-derived egg chamber formation
regulation of mitotic spindle organization
C) Regulation, Kinase, Negative, Epidermal, Erbb, Hormone, Signaling, Phosphorylation, Pathway, Peptidyltyrosine, Dorsalventral, Growth, Protein, Axis, Tyrosine, Receptor, Factor, Specification
negative regulation of JUN kinase activity
negative regulation of catalytic activity
negative regulation of protein phosphorylation
regulation of cyclin-dependent protein serine/threonine kinase activity
negative regulation of kinase activity
negative regulation of protein kinase activity
dorsal/ventral axis specification
regulation of protein kinase activity
negative regulation of hormone biosynthetic process
dorsal/ventral axis specification, ovarian follicular epithelium
regulation of carbohydrate metabolic process
negative regulation of hormone metabolic process
germ-line cyst formation
negative regulation of ERBB signaling pathway
negative regulation of epidermal growth factor receptor signaling pathway
regulation of hormone biosynthetic process
negative regulation of epidermal growth factor receptor signaling pathway
regulation of ERBB signaling pathway
epidermal growth factor receptor signaling pathway
regulation of protein tyrosine kinase activity
ERBB signaling pathway
regulation of peptidyl-tyrosine phosphorylation
regulation of hormone metabolic process
negative regulation of epidermal growth factor-activated receptor activity
negative regulation of peptidyl-tyrosine phosphorylation
negative regulation of protein tyrosine kinase activity
R7 cell fate commitment
negative regulation of signaling receptor activity
Z-score
0
1.5
3
Node Size: Gene Set Size
5.00
128.50
252.00
375.50
499.00
Node Border Width: -log(q-value)
8
2
1.3
1.6
1.9
2.3
2.6
Node Fill Color
Shared
Late
Early

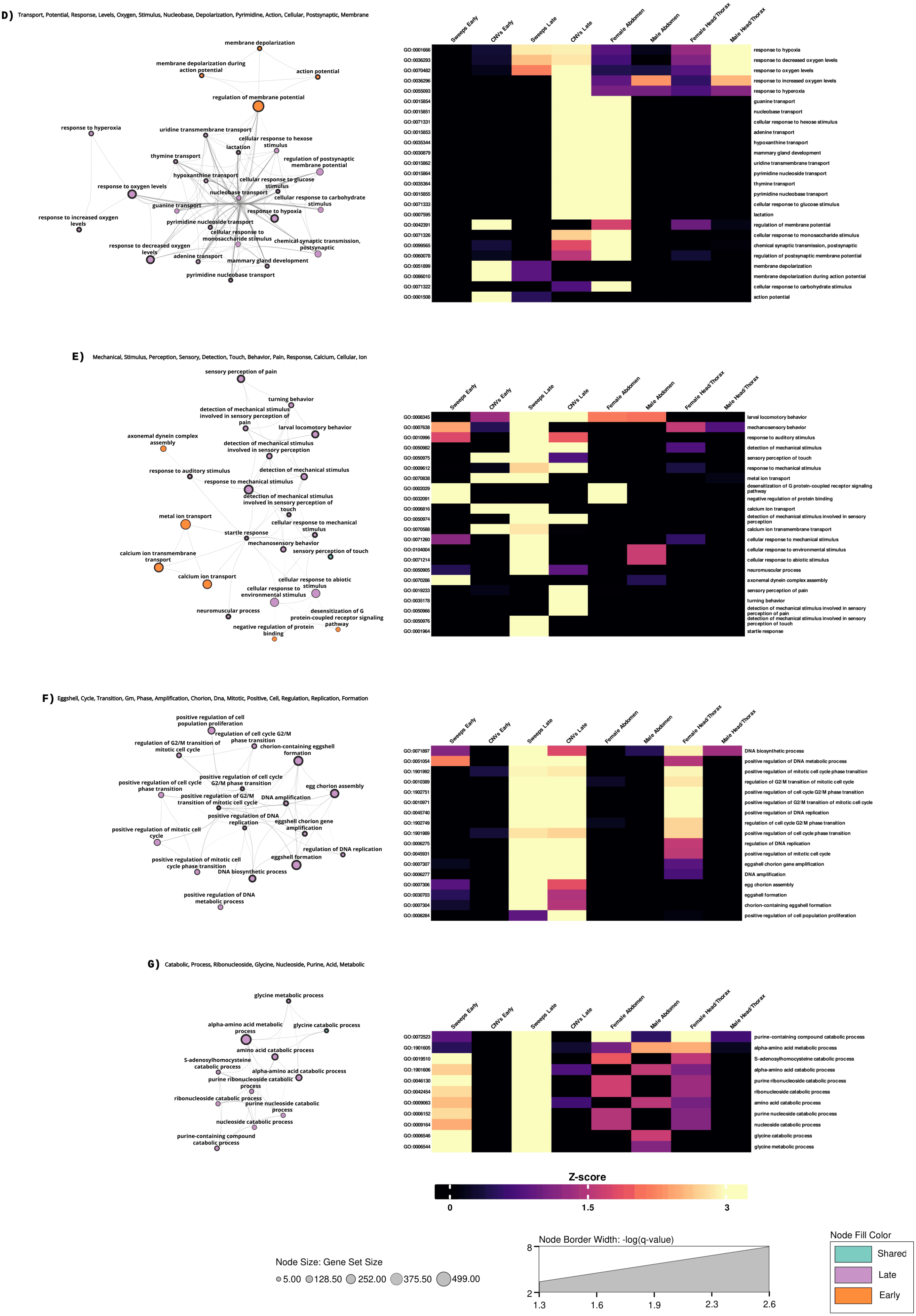

D) Transport, Potential, Response, Levels, Oxygen, Stimulus, Nucleobase, Depolarization, Pyrimidine, Action, Cellular, Postsynaptic, Membrane
Sweeps Early
CNVs Early
Sweeps Late
CNVs Late
Female Abdomen
Male Abdomen
Female Head/Thorax
Male Head/Thorax
GO:0001666 response to hypoxia
GO:0036293 response to decreased oxygen levels
GO:0070482 response to oxygen levels
GO:0036296 response to increased oxygen levels
GO:0055093 response to hyperoxia
GO:0015854 guanine transport
GO:0015851 nucleobase transport
GO:0071331 cellular response to hexose stimulus
GO:0015853 adenine transport
GO:0035344 hypoxanthine transport
GO:0030879 mammary gland development
GO:0015862 uridine transmembrane transport
GO:0015864 pyrimidine nucleoside transport
GO:0035364 thymine transport
GO:0015855 pyrimidine nucleobase transport
GO:0071333 cellular response to glucose stimulus
GO:0007595 lactation
GO:0042391 regulation of membrane potential
GO:0071326 cellular response to monosaccharide stimulus
GO:0099565 chemical synaptic transmission, postsynaptic
GO:0060078 regulation of postsynaptic membrane potential
GO:0051899 membrane depolarization
GO:0086010 membrane depolarization during action potential
GO:0071322 cellular response to carbohydrate stimulus
GO:0001508 action potential
E) Mechanical, Stimulus, Perception, Sensory, Detection, Touch, Behavior, Pain, Response, Calcium, Cellular, Ion
GO:0008345 larval locomotory behavior
GO:0007638 mechanosensory behavior
GO:0010996 response to auditory stimulus
GO:0050982 detection of mechanical stimulus
GO:0050975 sensory perception of touch
GO:0009612 response to mechanical stimulus
GO:0070838 metal ion transport
GO:0002029 desensitization of G protein-coupled receptor signaling pathway
GO:0032091 negative regulation of protein binding
GO:0006816 calcium ion transport
GO:0050974 detection of mechanical stimulus involved in sensory perception
GO:0070588 calcium ion transmembrane transport
GO:0071260 cellular response to mechanical stimulus
GO:0104004 cellular response to environmental stimulus
GO:0071214 cellular response to abiotic stimulus
GO:0050905 neuromuscular process
GO:0070286 axonemal dynein complex assembly
GO:0019233 sensory perception of pain
GO:0035178 turning behavior
GO:0050966 detection of mechanical stimulus involved in sensory perception of pain
GO:0050976 detection of mechanical stimulus involved in sensory perception of touch
GO:0001964 startle response
F) Eggshell, Cycle, Transition, Gm, Phase, Amplification, Chorion, Dna, Mitotic, Positive, Cell, Regulation, Replication, Formation
GO:0071897 DNA biosynthetic process
GO:0051054 positive regulation of DNA metabolic process
GO:1901992 positive regulation of mitotic cell cycle phase transition
GO:0010389 regulation of G2/M transition of mitotic cell cycle
GO:1902751 positive regulation of cell cycle G2/M phase transition
GO:0010971 positive regulation of G2/M transition of mitotic cell cycle
GO:0045740 positive regulation of DNA replication
GO:1902749 regulation of cell cycle G2/M phase transition
GO:1901989 positive regulation of cell cycle phase transition
GO:0006275 regulation of DNA replication
GO:0045931 positive regulation of mitotic cell cycle
GO:0007307 eggshell chorion gene amplification
GO:0006277 DNA amplification
GO:0007306 egg chorion assembly
GO:0030703 eggshell formation
GO:0007304 chorion-containing eggshell formation
GO:0008284 positive regulation of cell population proliferation
G) Catabolic, Process, Ribonucleoside, Glycine, Nucleoside, Purine, Acid, Metabolic
GO:0072523 purine-containing compound catabolic process
GO:1901605 alpha-amino acid metabolic process
GO:0019510 S-adenosylhomocysteine catabolic process
GO:1901606 alpha-amino acid catabolic process
GO:0046130 purine ribonucleoside catabolic process
GO:0042454 ribonucleoside catabolic process
GO:0009063 amino acid catabolic process
GO:0006152 purine nucleoside catabolic process
GO:0009164 nucleoside catabolic process
GO:0006546 glycine catabolic process
GO:0006544 glycine metabolic process
Z-score
0
1.5
3
Node Size: Gene Set Size
5.00
128.50
252.00
375.50
499.00
Node Border Width: -log(q-value)
8
2
1.3
1.6
1.9
2.3
2.6
Node Fill Color
Shared
Late
Early

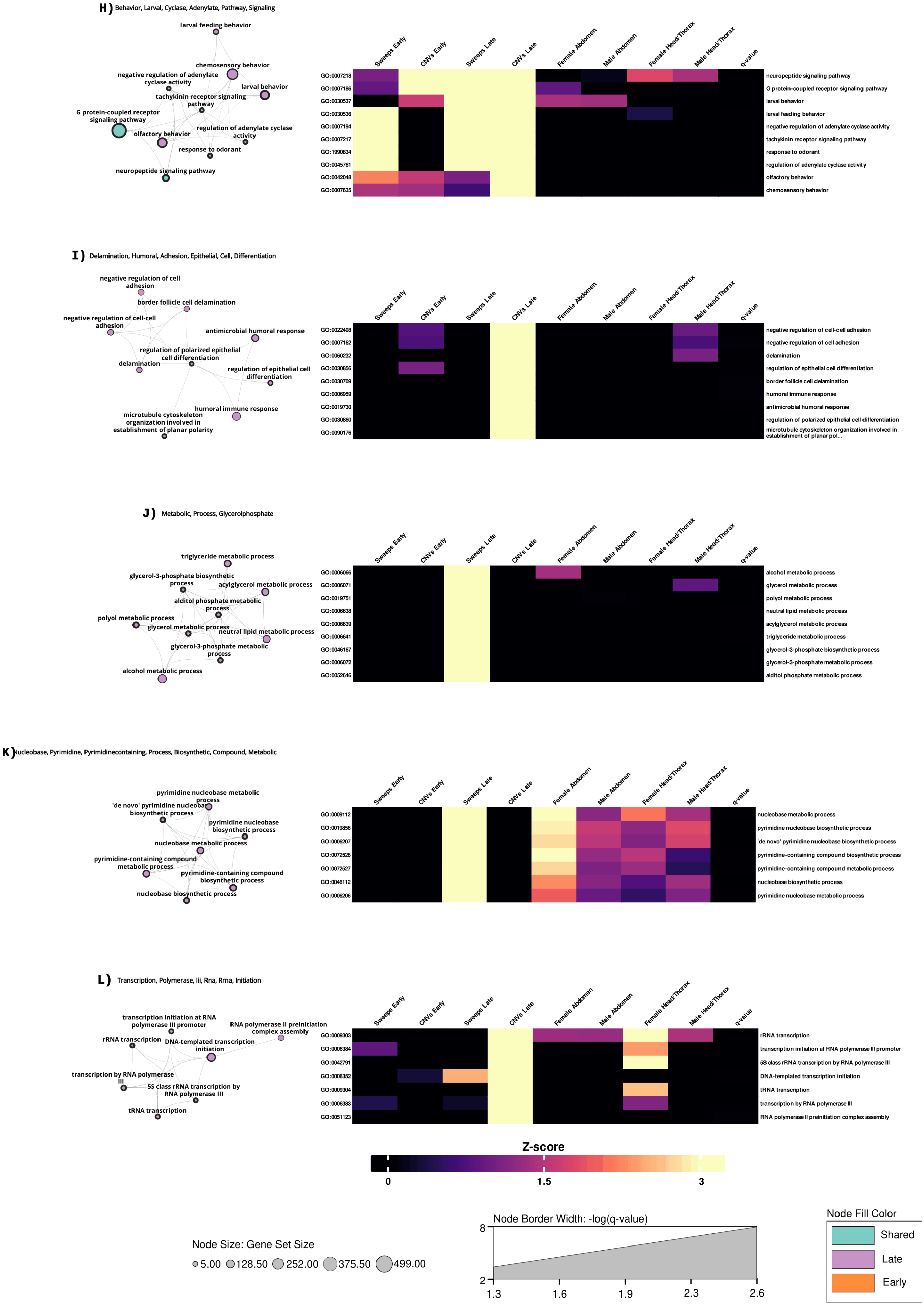
H) Behavior, Larval, Cyclase, Adenylate, Pathway, Signaling
larval feeding behavior
chemosensory behavior
negative regulation of adenylate cyclase activity
larval behavior
tachykinin receptor signaling pathway
G protein-coupled receptor signaling pathway
regulation of adenylate cyclase activity
olfactory behavior
response to odorant
neuropeptide signaling pathway
Sweeps Early
CNVs Early
Sweeps Late
CNVs Late
Female Abdomen
Male Abdomen
Female Head/Thorax
Male Head/Thorax
q-value
GO:0007218 neuropeptide signaling pathway
GO:0007186 G protein-coupled receptor signaling pathway
GO:0030537 larval behavior
GO:0030536 larval feeding behavior
GO:0007194 negative regulation of adenylate cyclase activity
GO:0007217 tachykinin receptor signaling pathway
GO:1990834 response to odorant
GO:0045761 regulation of adenylate cyclase activity
GO:0042048 olfactory behavior
GO:0007635 chemosensory behavior
I) Delamination, Humoral, Adhesion, Epithelial, Cell, Differentiation
negative regulation of cell adhesion
border follicle cell delamination
negative regulation of cell-cell adhesion
antimicrobial humoral response
regulation of polarized epithelial cell differentiation
delamination
regulation of epithelial cell differentiation
humoral immune response
microtubule cytoskeleton organization involved in establishment of planar polarity
GO:0022408 negative regulation of cell-cell adhesion
GO:0007162 negative regulation of cell adhesion
GO:0060232 delamination
GO:0030856 regulation of epithelial cell differentiation
GO:0030709 border follicle cell delamination
GO:0006959 humoral immune response
GO:0019730 antimicrobial humoral response
GO:0030860 regulation of polarized epithelial cell differentiation
GO:0090176 microtubule cytoskeleton organization involved in establishment of planar pol...
J) Metabolic, Process, Glycerolphosphate
triglyceride metabolic process
glycerol-3-phosphate biosynthetic process
acylglycerol metabolic process
alditol phosphate metabolic process
polyol metabolic process
glycerol metabolic process
neutral lipid metabolic process
glycerol-3-phosphate metabolic process
alcohol metabolic process
GO:0006066 alcohol metabolic process
GO:0006071 glycerol metabolic process
GO:0019751 polyol metabolic process
GO:0006638 neutral lipid metabolic process
GO:0006639 acylglycerol metabolic process
GO:0006641 triglyceride metabolic process
GO:0046167 glycerol-3-phosphate biosynthetic process
GO:0006072 glycerol-3-phosphate metabolic process
GO:0052646 alditol phosphate metabolic process
K) Nucleobase, Pyrimidine, Pyrimidinecontaining, Process, Biosynthetic, Compound, Metabolic
pyrimidine nucleobase metabolic process
'de novo' pyrimidine nucleobase biosynthetic process
pyrimidine nucleobase biosynthetic process
nucleobase metabolic process
pyrimidine-containing compound metabolic process
pyrimidine-containing compound biosynthetic process
nucleobase biosynthetic process
GO:0009112 nucleobase metabolic process
GO:0019856 pyrimidine nucleobase biosynthetic process
GO:0006207 'de novo' pyrimidine nucleobase biosynthetic process
GO:0072528 pyrimidine-containing compound biosynthetic process
GO:0072527 pyrimidine-containing compound metabolic process
GO:0046112 nucleobase biosynthetic process
GO:0006206 pyrimidine nucleobase metabolic process
L) Transcription, Polymerase, Iii, Rna, Rrna, Initiation
transcription initiation at RNA polymerase III promoter
RNA polymerase II preinitiation complex assembly
rRNA transcription
DNA-templated transcription initiation
transcription by RNA polymerase III
5S class rRNA transcription by RNA polymerase III
tRNA transcription
GO:0009303 rRNA transcription
GO:0006384 transcription initiation at RNA polymerase III promoter
GO:0042791 5S class rRNA transcription by RNA polymerase III
GO:0006352 DNA-templated transcription initiation
GO:0009304 tRNA transcription
GO:0006383 transcription by RNA polymerase III
GO:0051123 RNA polymerase II preinitiation complex assembly
Z-score
0
1.5
3
Node Size: Gene Set Size
5.00
128.50
252.00
375.50
499.00
Node Border Width: -log(q-value)
8
2
1.3
1.6
1.9
2.3
2.6
Node Fill Color
Shared
Late
Early

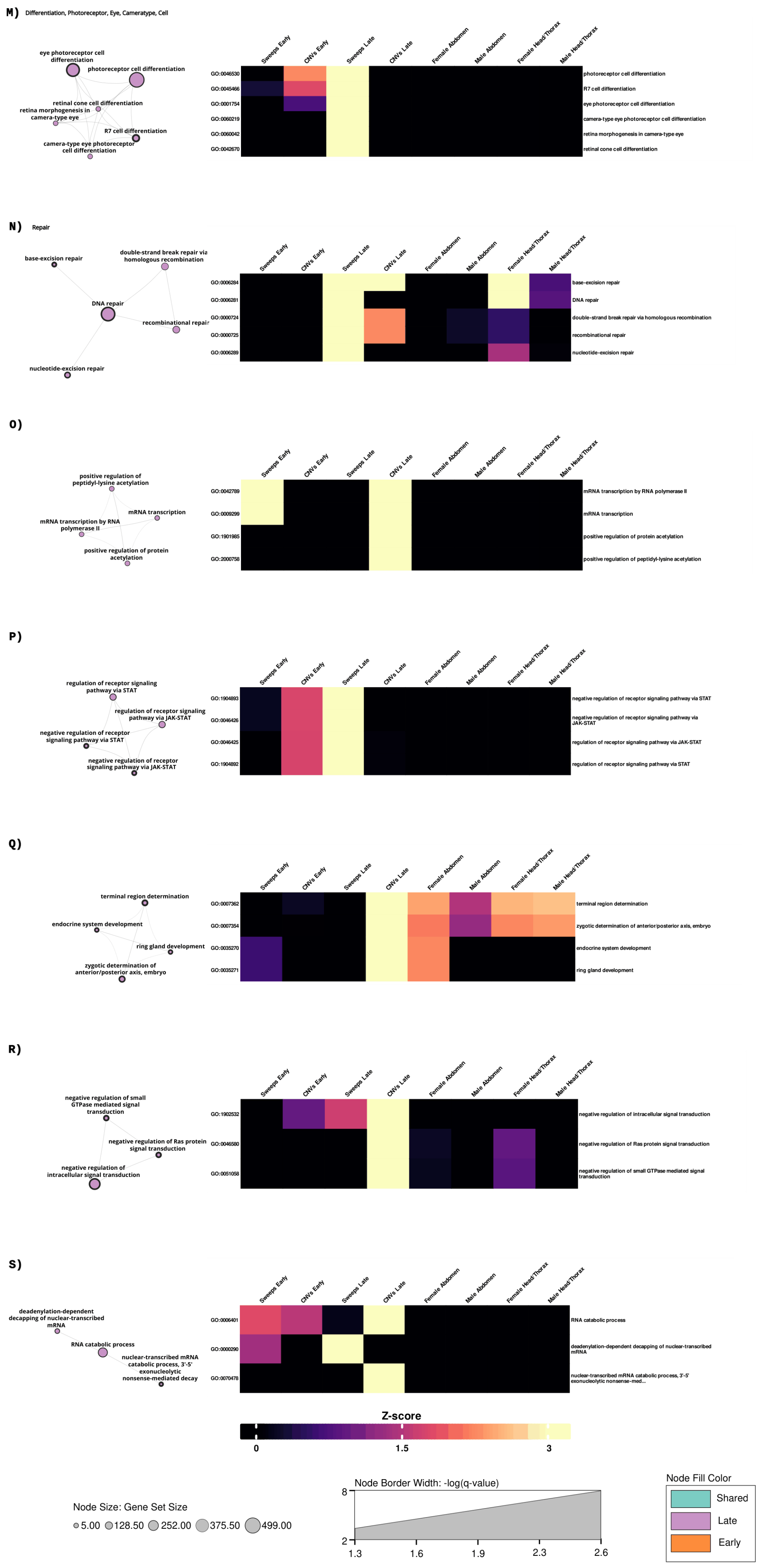
M) Differentiation, Photoreceptor, Eye, Cameratype, Cell
eye photoreceptor cell differentiation
photoreceptor cell differentiation
retinal cone cell differentiation
retina morphogenesis in camera-type eye
R7 cell differentiation
camera-type eye photoreceptor cell differentiation
Sweeps Early
CNVs Early
Sweeps Late
CNVs Late
Female Abdomen
Male Abdomen
Female Head/Thorax
Male Head/Thorax
GO:0046530
GO:0045466
GO:0001754
GO:0060219
GO:0060042
GO:0042670
photoreceptor cell differentiation
R7 cell differentiation
eye photoreceptor cell differentiation
camera-type eye photoreceptor cell differentiation
retina morphogenesis in camera-type eye
retinal cone cell differentiation
N) Repair
base-excision repair
double-strand break repair via homologous recombination
DNA repair
recombinational repair
nucleotide-excision repair
GO:0006284
GO:0006281
GO:0000724
GO:0000725
GO:0006289
base-excision repair
DNA repair
double-strand break repair via homologous recombination
recombinational repair
nucleotide-excision repair
O)
positive regulation of peptidyl-lysine acetylation
mRNA transcription
mRNA transcription by RNA polymerase II
positive regulation of protein acetylation
GO:0042789
GO:0009299
GO:1901985
GO:2000758
mRNA transcription by RNA polymerase II
mRNA transcription
positive regulation of protein acetylation
positive regulation of peptidyl-lysine acetylation
P)
regulation of receptor signaling pathway via STAT
regulation of receptor signaling pathway via JAK-STAT
negative regulation of receptor signaling pathway via STAT
negative regulation of receptor signaling pathway via JAK-STAT
GO:1904893
GO:0046426
GO:0046425
GO:1904892
negative regulation of receptor signaling pathway via STAT
negative regulation of receptor signaling pathway via JAK-STAT
regulation of receptor signaling pathway via JAK-STAT
regulation of receptor signaling pathway via STAT
Q)
terminal region determination
endocrine system development
ring gland development
zygotic determination of anterior/posterior axis, embryo
GO:0007362
GO:0007354
GO:0035270
GO:0035271
terminal region determination
zygotic determination of anterior/posterior axis, embryo
endocrine system development
ring gland development
R)
negative regulation of small GTPase mediated signal transduction
negative regulation of Ras protein signal transduction
negative regulation of intracellular signal transduction
GO:1902532
GO:0046580
GO:0051058
negative regulation of intracellular signal transduction
negative regulation of Ras protein signal transduction
negative regulation of small GTPase mediated signal transduction
S)
deadenylation-dependent decapping of nuclear-transcribed mRNA
RNA catabolic process
nuclear-transcribed mRNA catabolic process, 3'-5' exonucleolytic nonsense-mediated decay
GO:0006401
GO:0000290
GO:0070478
RNA catabolic process
deadenylation-dependent decapping of nuclear-transcribed mRNA
nuclear-transcribed mRNA catabolic process, 3'-5' exonucleolytic nonsense-med...
Z-score
0
1.5
3
Node Size: Gene Set Size
5.00
128.50
252.00
375.50
499.00
Node Border Width: -log(q-value)
8
2
1.3
1.6
1.9
2.3
2.6
Node Fill Color
Shared
Late
Early

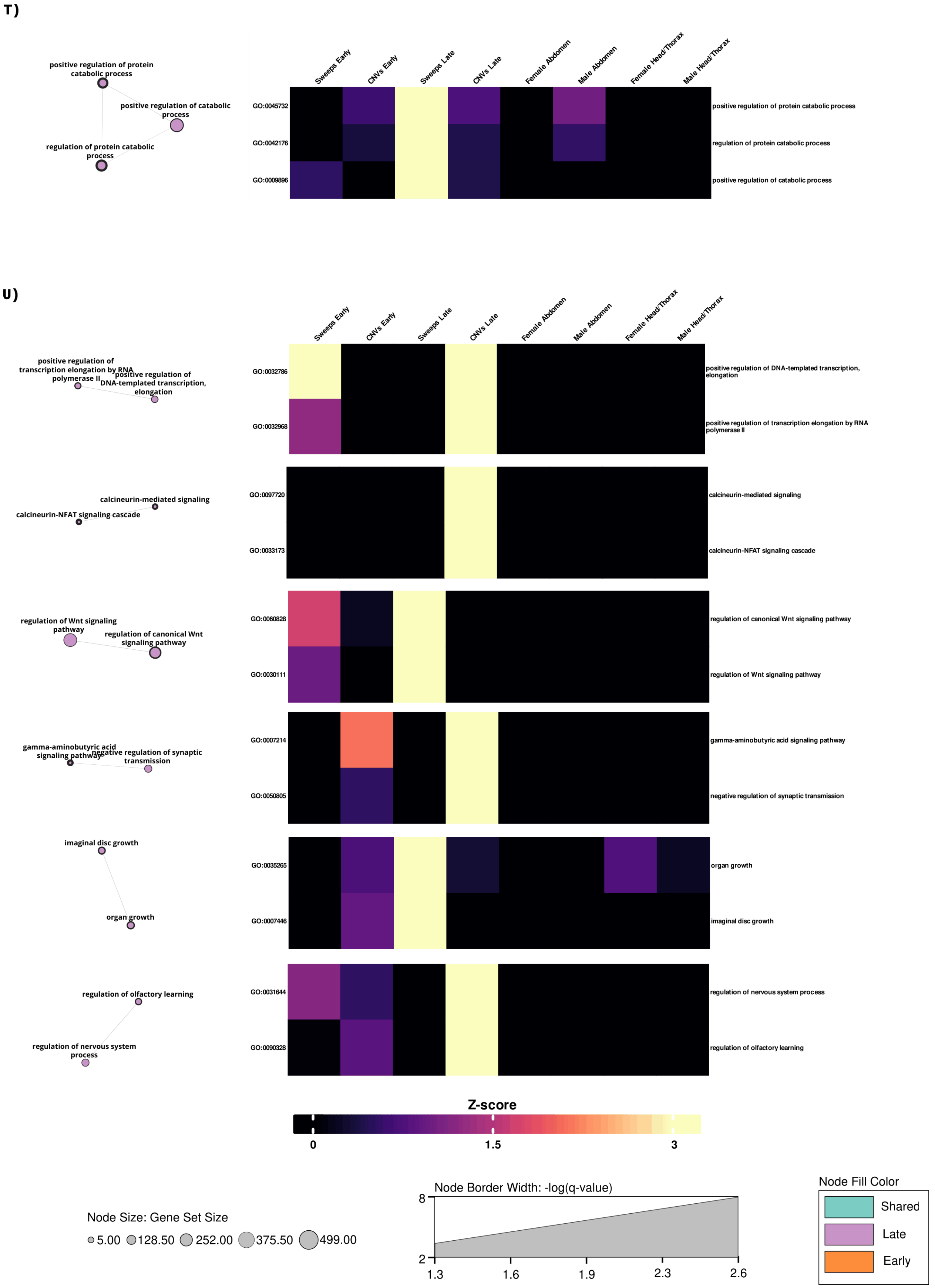

**Fig. S35.** Functional modules (A-U) identified in our multi-evidence enrichment analysis, where evidence for enrichment was present in L lines. The left most four columns in the heat-map represents sweep and CNV calls, the right most four columns differential gene expression (color denotes strength of evidence).

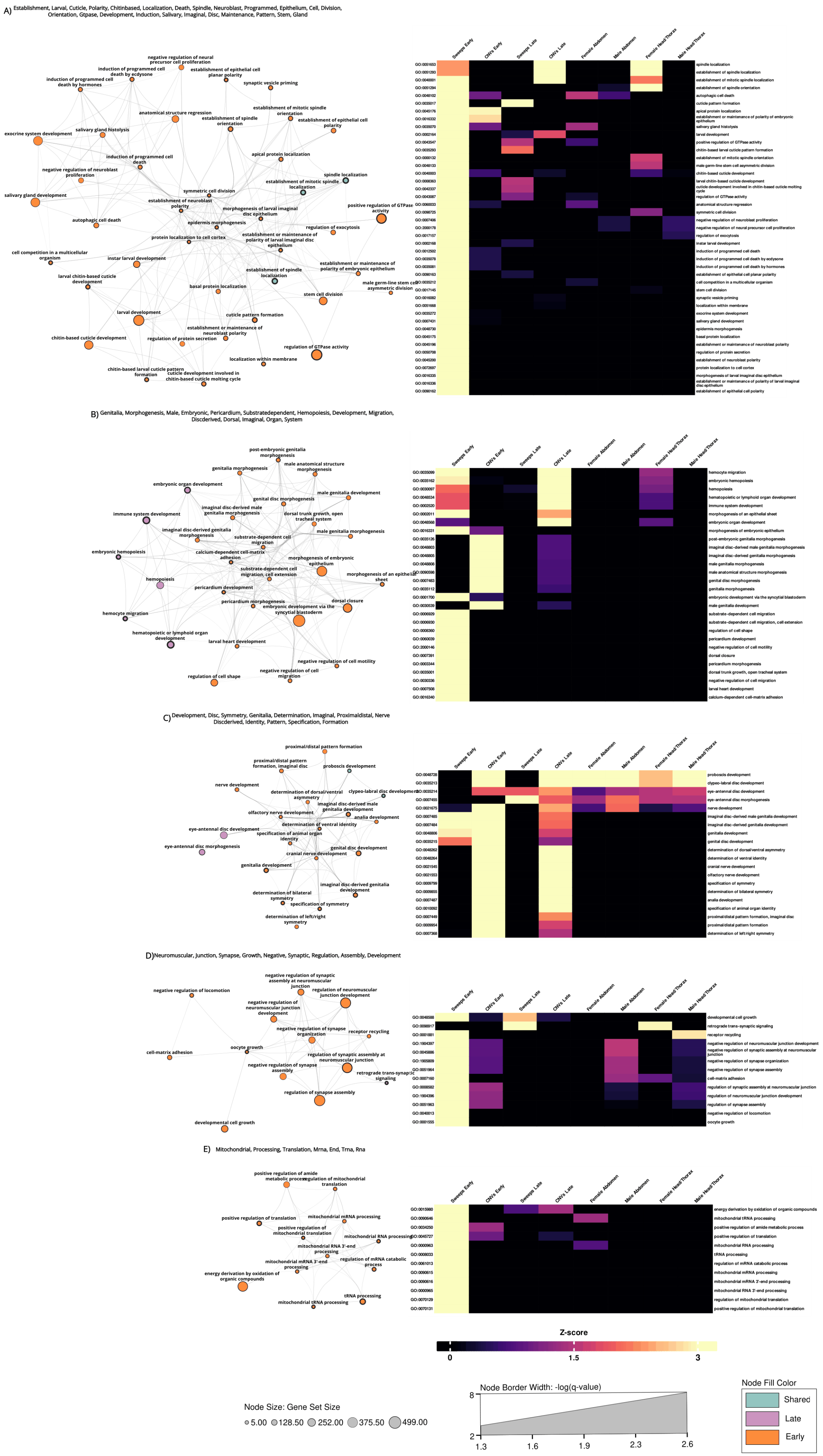

A) Establishment, Larval, Cuticle, Polarity, Chitinbased, Localization, Death, Spindle, Neuroblast, Programmed, Epithelium, Cell, Division, Orientation, Gtpase, Development, Induction, Salivary, Imaginal, Disc, Maintenance, Pattern, Stem, Gland
B) Genitalia, Morphogenesis, Male, Embryonic, Pericardium, Substratedependent, Hemopoiesis, Development, Migration, Discderived, Dorsal, Imaginal, Organ, System
C) Development, Disc, Symmetry, Genitalia, Determination, Imaginal, Proximaldistal, Nerve Discderived, Identity, Pattern, Specification, Formation
D) Neuromuscular, Junction, Synapse, Growth, Negative, Synaptic, Regulation, Assembly, Development
E) Mitochondrial, Processing, Translation, Mrna, End, Trna, Rna
Sweeps Early
CNVs Early
Sweeps Late
CNVs Late
Female Abdomen
Male Abdomen
Female Head/Thorax
Male Head/Thorax
Z-score
0
1.5
3
Node Size: Gene Set Size
5.00
128.50
252.00
375.50
499.00
Node Border Width: -log(q-value)
8
2
1.3
1.6
1.9
2.3
2.6
Node Fill Color
Shared
Late
Early

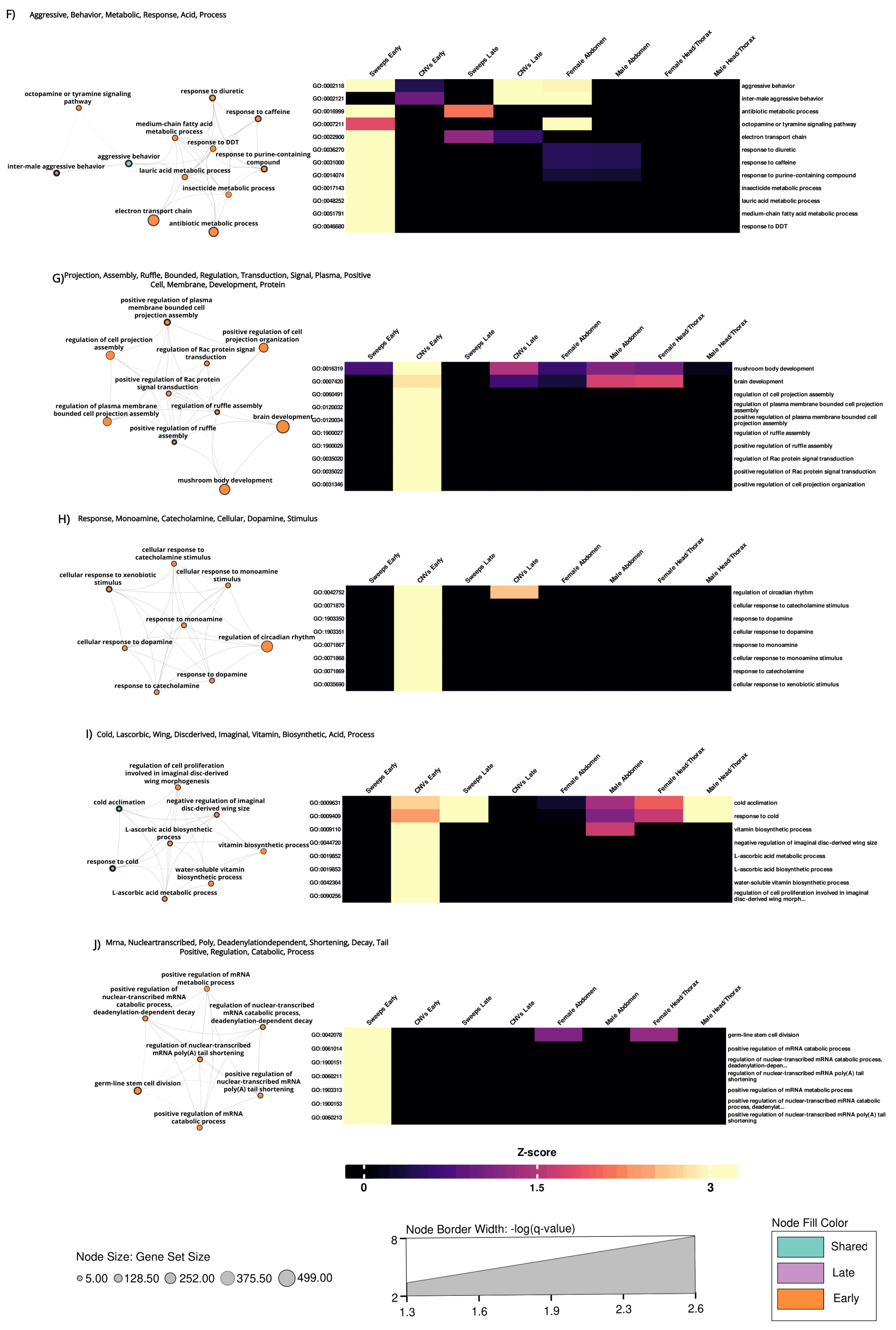

F) Aggressive, Behavior, Metabolic, Response, Acid, Process
octopamine or tyramine signaling pathway
response to diuretic
response to caffeine
medium-chain fatty acid metabolic process
response to DDT
aggressive behavior
response to purine-containing compound
inter-male aggressive behavior
lauric acid metabolic process
insecticide metabolic process
electron transport chain
antibiotic metabolic process
Sweeps Early
CNVs Early
Sweeps Late
CNVs Late
Female Abdomen
Male Abdomen
Female Head/Thorax
Male Head/Thorax
GO:0002118
GO:0002121
GO:0016999
GO:0007211
GO:0022900
GO:0036270
GO:0031000
GO:0014074
GO:0017143
GO:0048252
GO:0051791
GO:0046680
aggressive behavior
inter-male aggressive behavior
antibiotic metabolic process
octopamine or tyramine signaling pathway
electron transport chain
response to diuretic
response to caffeine
response to purine-containing compound
insecticide metabolic process
lauric acid metabolic process
medium-chain fatty acid metabolic process
response to DDT
G) Projection, Assembly, Ruffle, Bounded, Regulation, Transduction, Signal, Plasma, Positive Cell, Membrane, Development, Protein
positive regulation of plasma membrane bounded cell projection assembly
positive regulation of cell projection organization
regulation of cell projection assembly
regulation of Rac protein signal transduction
positive regulation of Rac protein signal transduction
regulation of plasma membrane bounded cell projection assembly
regulation of ruffle assembly
brain development
positive regulation of ruffle assembly
mushroom body development
Sweeps Early
CNVs Early
Sweeps Late
CNVs Late
Female Abdomen
Male Abdomen
Female Head/Thorax
Male Head/Thorax
GO:0016319
GO:0007420
GO:0060491
GO:0120032
GO:0120034
GO:1900027
GO:1900029
GO:0035020
GO:0035022
GO:0031346
mushroom body development
brain development
regulation of cell projection assembly
regulation of plasma membrane bounded cell projection assembly
positive regulation of plasma membrane bounded cell projection assembly
regulation of ruffle assembly
positive regulation of ruffle assembly
regulation of Rac protein signal transduction
positive regulation of Rac protein signal transduction
positive regulation of cell projection organization
H) Response, Monoamine, Catecholamine, Cellular, Dopamine, Stimulus
cellular response to catecholamine stimulus
cellular response to xenobiotic stimulus
cellular response to monoamine stimulus
response to monoamine
regulation of circadian rhythm
cellular response to dopamine
response to dopamine
response to catecholamine
Sweeps Early
CNVs Early
Sweeps Late
CNVs Late
Female Abdomen
Male Abdomen
Female Head/Thorax
Male Head/Thorax
GO:0042752
GO:0071870
GO:1903350
GO:1903351
GO:0071867
GO:0071868
GO:0071869
GO:0035690
regulation of circadian rhythm
cellular response to catecholamine stimulus
response to dopamine
cellular response to dopamine
response to monoamine
cellular response to monoamine stimulus
response to catecholamine
cellular response to xenobiotic stimulus
I) Cold, Lascorbic, Wing, Discderived, Imaginal, Vitamin, Biosynthetic, Acid, Process
regulation of cell proliferation involved in imaginal disc-derived wing morphogenesis
cold acclimation
negative regulation of imaginal disc-derived wing size
L-ascorbic acid biosynthetic process
vitamin biosynthetic process
response to cold
water-soluble vitamin biosynthetic process
L-ascorbic acid metabolic process
Sweeps Early
CNVs Early
Sweeps Late
CNVs Late
Female Abdomen
Male Abdomen
Female Head/Thorax
Male Head/Thorax
GO:0009631
GO:0009409
GO:0009110
GO:0044720
GO:0019852
GO:0019853
GO:0042364
GO:0090256
cold acclimation
response to cold
vitamin biosynthetic process
negative regulation of imaginal disc-derived wing size
L-ascorbic acid metabolic process
L-ascorbic acid biosynthetic process
water-soluble vitamin biosynthetic process
regulation of cell proliferation involved in imaginal disc-derived wing morph...
J) Mrna, Nucleartranscribed, Poly, Deadenylationdependent, Shortening, Decay, Tail Positive, Regulation, Catabolic, Process
positive regulation of mRNA metabolic process
positive regulation of nuclear-transcribed mRNA catabolic process, deadenylation-dependent decay
regulation of nuclear-transcribed mRNA catabolic process, deadenylation-dependent decay
regulation of nuclear-transcribed mRNA poly(A) tail shortening
germ-line stem cell division
positive regulation of nuclear-transcribed mRNA poly(A) tail shortening
positive regulation of mRNA catabolic process
Sweeps Early
CNVs Early
Sweeps Late
CNVs Late
Female Abdomen
Male Abdomen
Female Head/Thorax
Male Head/Thorax
GO:0042078
GO:0061014
GO:1900151
GO:0060211
GO:1903313
GO:1900153
GO:0060213
germ-line stem cell division
positive regulation of mRNA catabolic process
regulation of nuclear-transcribed mRNA catabolic process, deadenylation-depen...
regulation of nuclear-transcribed mRNA poly(A) tail shortening
positive regulation of mRNA metabolic process
positive regulation of nuclear-transcribed mRNA catabolic process, deadenylat...
positive regulation of nuclear-transcribed mRNA poly(A) tail shortening
Z-score
0
1.5
3
Node Border Width: -log(q-value)
8
2
1.3
1.6
1.9
2.3
2.6
Node Size: Gene Set Size
5.00
128.50
252.00
375.50
499.00
Node Fill Color
Shared
Late
Early

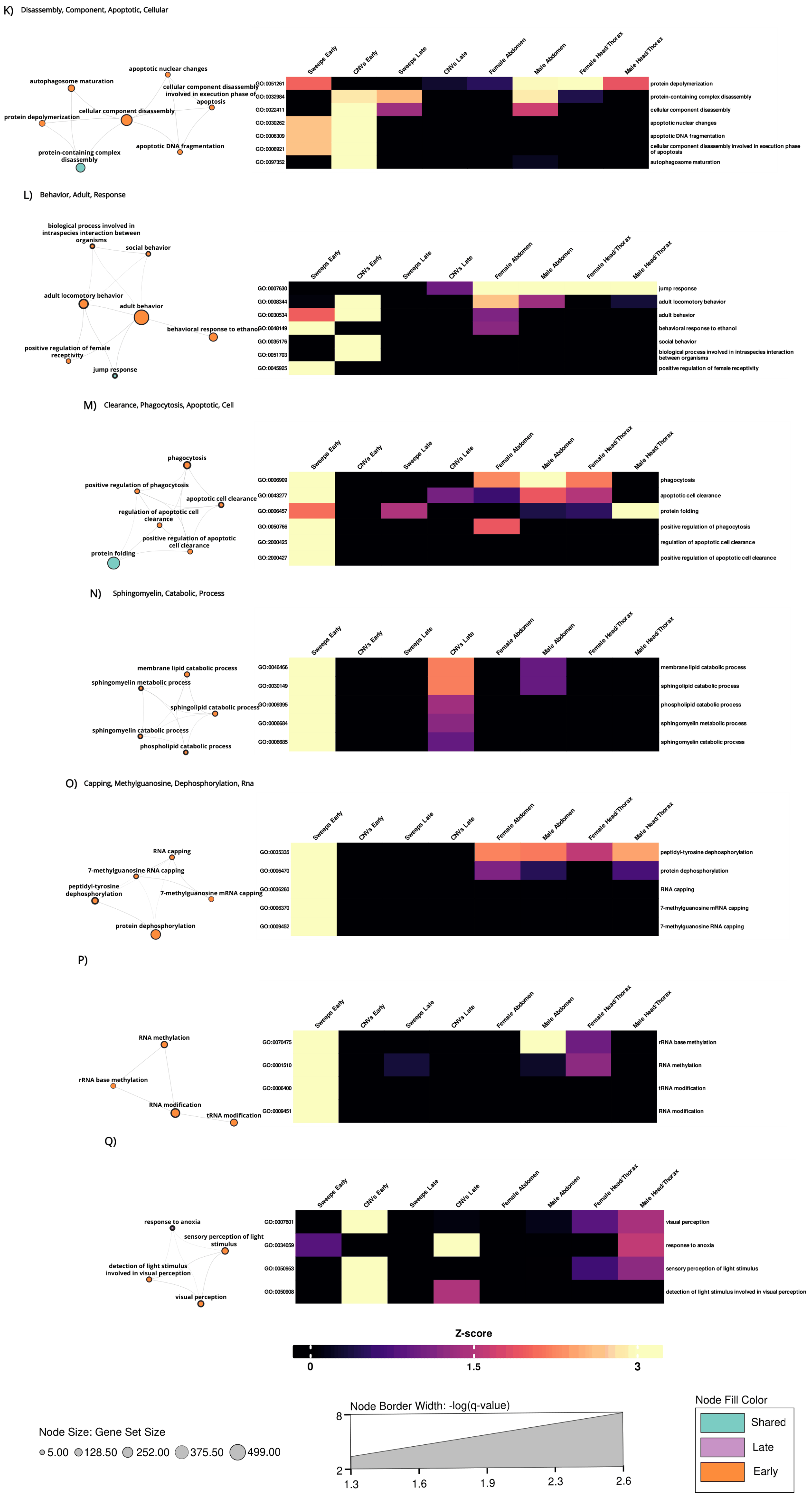
K) Disassembly, Component, Apoptotic, Cellular
apoptotic nuclear changes
autophagosome maturation
cellular component disassembly involved in execution phase of apoptosis
cellular component disassembly
protein depolymerization
apoptotic DNA fragmentation
protein-containing complex disassembly
Sweeps Early
CNVs Early
Sweeps Late
CNVs Late
Female Abdomen
Male Abdomen
Female Head/Thorax
Male Head/Thorax
GO:0051261
GO:0032984
GO:0022411
GO:0030262
GO:0006309
GO:0006921
GO:0097352
protein depolymerization
protein-containing complex disassembly
cellular component disassembly
apoptotic nuclear changes
apoptotic DNA fragmentation
cellular component disassembly involved in execution phase of apoptosis
autophagosome maturation
L) Behavior, Adult, Response
biological process involved in intraspecies interaction between organisms
social behavior
adult locomotory behavior
adult behavior
behavioral response to ethanol
positive regulation of female receptivity
jump response
GO:0007630
GO:0008344
GO:0030534
GO:0048149
GO:0035176
GO:0051703
GO:0045925
jump response
adult locomotory behavior
adult behavior
behavioral response to ethanol
social behavior
biological process involved in intraspecies interaction between organisms
positive regulation of female receptivity
M) Clearance, Phagocytosis, Apoptotic, Cell
phagocytosis
positive regulation of phagocytosis
apoptotic cell clearance
regulation of apoptotic cell clearance
positive regulation of apoptotic cell clearance
protein folding
GO:0006909
GO:0043277
GO:0006457
GO:0050766
GO:2000425
GO:2000427
phagocytosis
apoptotic cell clearance
protein folding
positive regulation of phagocytosis
regulation of apoptotic cell clearance
positive regulation of apoptotic cell clearance
N) Sphingomyelin, Catabolic, Process
membrane lipid catabolic process
sphingomyelin metabolic process
sphingolipid catabolic process
sphingomyelin catabolic process
phospholipid catabolic process
GO:0046466
GO:0030149
GO:0009395
GO:0006684
GO:0006685
membrane lipid catabolic process
sphingolipid catabolic process
phospholipid catabolic process
sphingomyelin metabolic process
sphingomyelin catabolic process
O) Capping, Methylguanosine, Dephosphorylation, Rna
RNA capping
7-methylguanosine RNA capping
peptidyl-tyrosine dephosphorylation
7-methylguanosine mRNA capping
protein dephosphorylation
GO:0035335
GO:0006470
GO:0036260
GO:0006370
GO:0009452
peptidyl-tyrosine dephosphorylation
protein dephosphorylation
RNA capping
7-methylguanosine mRNA capping
7-methylguanosine RNA capping
P)
RNA methylation
rRNA base methylation
RNA modification
tRNA modification
GO:0070475
GO:0001510
GO:0006400
GO:0009451
rRNA base methylation
RNA methylation
tRNA modification
RNA modification
Q)
response to anoxia
sensory perception of light stimulus
detection of light stimulus involved in visual perception
visual perception
GO:0007601
GO:0034059
GO:0050953
GO:0050908
visual perception
response to anoxia
sensory perception of light stimulus
detection of light stimulus involved in visual perception
Z-score
0
1.5
3
Node Size: Gene Set Size
5.00
128.50
252.00
375.50
499.00
Node Border Width: -log(q-value)
8
2
1.3
1.6
1.9
2.3
2.6
Node Fill Color
Shared
Late
Early

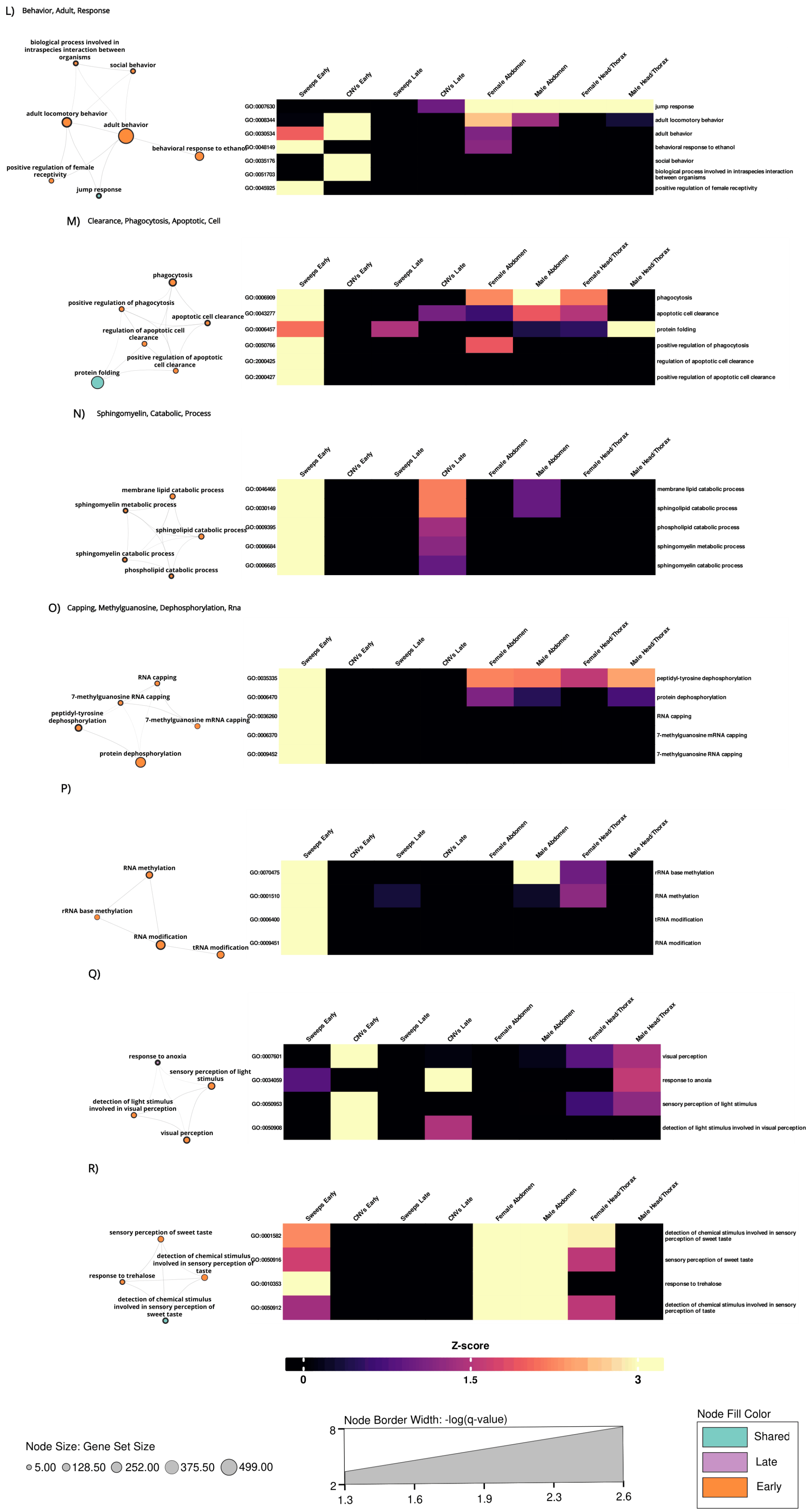

L) Behavior, Adult, Response
M) Clearance, Phagocytosis, Apoptotic, Cell
N) Sphingomyelin, Catabolic, Process
O) Capping, Methylguanosine, Dephosphorylation, Rna
P)
Q)
R)
Sweeps Early
CNVs Early
Sweeps Late
CNVs Late
Female Abdomen
Male Abdomen
Female Head/Thorax
Male Head/Thorax
GO:0007630 jump response
GO:0008344 adult locomotory behavior
GO:0030534 adult behavior
GO:0048149 behavioral response to ethanol
GO:0035176 social behavior
GO:0051703 biological process involved in intraspecies interaction between organisms
GO:0045925 positive regulation of female receptivity
GO:0006909 phagocytosis
GO:0043277 apoptotic cell clearance
GO:0006457 protein folding
GO:0050766 positive regulation of phagocytosis
GO:2000425 regulation of apoptotic cell clearance
GO:2000427 positive regulation of apoptotic cell clearance
GO:0046466 membrane lipid catabolic process
GO:0030149 sphingolipid catabolic process
GO:0009395 phospholipid catabolic process
GO:0006684 sphingomyelin metabolic process
GO:0006685 sphingomyelin catabolic process
GO:0035335 peptidyl-tyrosine dephosphorylation
GO:0006470 protein dephosphorylation
GO:0036260 RNA capping
GO:0006370 7-methylguanosine mRNA capping
GO:0009452 7-methylguanosine RNA capping
GO:0070475 rRNA base methylation
GO:0001510 RNA methylation
GO:0006400 tRNA modification
GO:0009451 RNA modification
GO:0007601 visual perception
GO:0034059 response to anoxia
GO:0050953 sensory perception of light stimulus
GO:0050908 detection of light stimulus involved in visual perception
GO:0001582 detection of chemical stimulus involved in sensory perception of sweet taste
GO:0050916 sensory perception of sweet taste
GO:0010353 response to trehalose
GO:0050912 detection of chemical stimulus involved in sensory perception of taste
Z-score
0
1.5
3
Node Size: Gene Set Size
5.00
128.50
252.00
375.50
499.00
Node Border Width: -log(q-value)
8
2
1.3
1.6
1.9
2.3
2.6
Node Fill Color
Shared
Late
Early

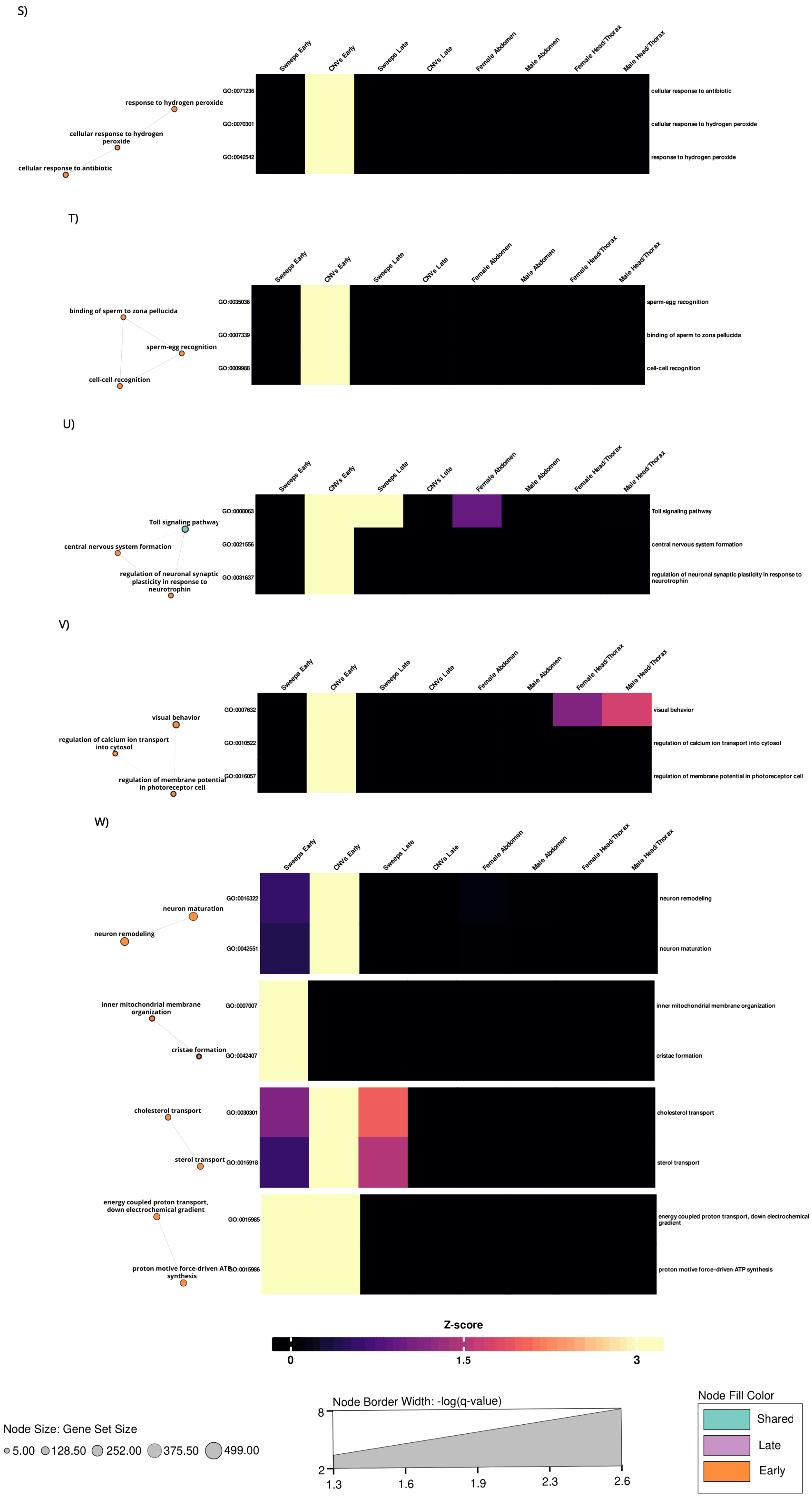

**Fig. S36.** Functional modules (A-W) identified in our multi-evidence enrichment analysis, where evidence for enrichment was present in E lines. The left most four columns in the heat-map represents sweep and CNV calls, the right most four columns differential gene expression (color denotes strength of evidence).

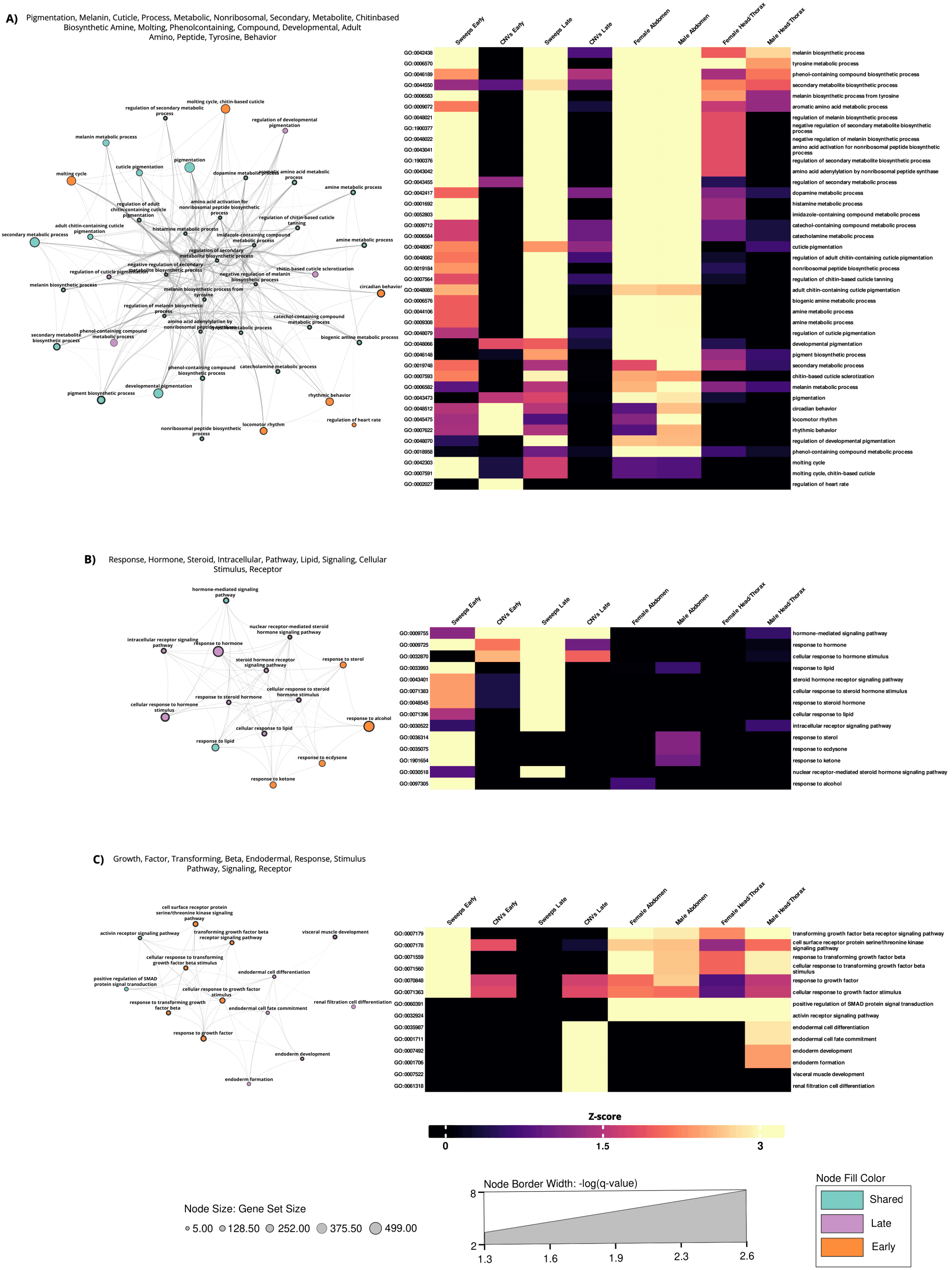
A) Pigmentation, Melanin, Cuticle, Process, Metabolic, Nonribosomal, Secondary, Metabolite, Chitinbased Biosynthetic Amine, Molting, Phenolcontaining, Compound, Developmental, Adult Amino, Peptide, Tyrosine, Behavior
Sweeps Early
CNVs Early
Sweeps Late
CNVs Late
Female Abdomen
Male Abdomen
Female Head/Thorax
Male Head/Thorax
GO:0042438 melanin biosynthetic process
GO:0006570 tyrosine metabolic process
GO:0046189 phenol-containing compound biosynthetic process
GO:0044550 secondary metabolite biosynthetic process
GO:0006583 melanin biosynthetic process from tyrosine
GO:0009072 aromatic amino acid metabolic process
GO:0048021 regulation of melanin biosynthetic process
GO:1900377 negative regulation of secondary metabolite biosynthetic process
GO:0048022 negative regulation of melanin biosynthetic process
GO:0043041 amino acid activation for nonribosomal peptide biosynthetic process
GO:1900376 regulation of secondary metabolite biosynthetic process
GO:0043042 amino acid adenylylation by nonribosomal peptide synthase
GO:0043455 regulation of secondary metabolic process
GO:0042417 dopamine metabolic process
GO:0001692 histamine metabolic process
GO:0052803 imidazole-containing compound metabolic process
GO:0009712 catechol-containing compound metabolic process
GO:0006584 catecholamine metabolic process
GO:0048067 cuticle pigmentation
GO:0048082 regulation of adult chitin-containing cuticle pigmentation
GO:0019184 nonribosomal peptide biosynthetic process
GO:0007564 regulation of chitin-based cuticle tanning
GO:0048085 adult chitin-containing cuticle pigmentation
GO:0006576 biogenic amine metabolic process
GO:0044106 amine metabolic process
GO:0009308 amine metabolic process
GO:0048079 regulation of cuticle pigmentation
GO:0048066 developmental pigmentation
GO:0046148 pigment biosynthetic process
GO:0019748 secondary metabolic process
GO:0007593 chitin-based cuticle sclerotization
GO:0006582 melanin metabolic process
GO:0043473 pigmentation
GO:0048512 circadian behavior
GO:0045475 locomotor rhythm
GO:0007622 rhythmic behavior
GO:0048070 regulation of developmental pigmentation
GO:0018958 phenol-containing compound metabolic process
GO:0042303 molting cycle
GO:0007591 molting cycle, chitin-based cuticle
GO:0002027 regulation of heart rate
B) Response, Hormone, Steroid, Intracellular, Pathway, Lipid, Signaling, Cellular Stimulus, Receptor
GO:0009755 hormone-mediated signaling pathway
GO:0009725 response to hormone
GO:0032870 cellular response to hormone stimulus
GO:0033993 response to lipid
GO:0043401 steroid hormone receptor signaling pathway
GO:0071383 cellular response to steroid hormone stimulus
GO:0048545 response to steroid hormone
GO:0071396 cellular response to lipid
GO:0030522 intracellular receptor signaling pathway
GO:0036314 response to sterol
GO:0035075 response to ecdysone
GO:1901654 response to ketone
GO:0030518 nuclear receptor-mediated steroid hormone signaling pathway
GO:0097305 response to alcohol
C) Growth, Factor, Transforming, Beta, Endodermal, Response, Stimulus Pathway, Signaling, Receptor
GO:0007179 transforming growth factor beta receptor signaling pathway
GO:0007178 cell surface receptor protein serine/threonine kinase signaling pathway
GO:0071559 response to transforming growth factor beta
GO:0071560 cellular response to transforming growth factor beta stimulus
GO:0070848 response to growth factor
GO:0071363 cellular response to growth factor stimulus
GO:0060391 positive regulation of SMAD protein signal transduction
GO:0032924 activin receptor signaling pathway
GO:0035987 endodermal cell differentiation
GO:0001711 endodermal cell fate commitment
GO:0007492 endoderm development
GO:0001706 endoderm formation
GO:0007522 visceral muscle development
GO:0061318 renal filtration cell differentiation
Z-score
0
1.5
3
Node Size: Gene Set Size
5.00
128.50
252.00
375.50
499.00
Node Border Width: -log(q-value)
8
2
1.3
1.6
1.9
2.3
2.6
Node Fill Color
Shared
Late
Early

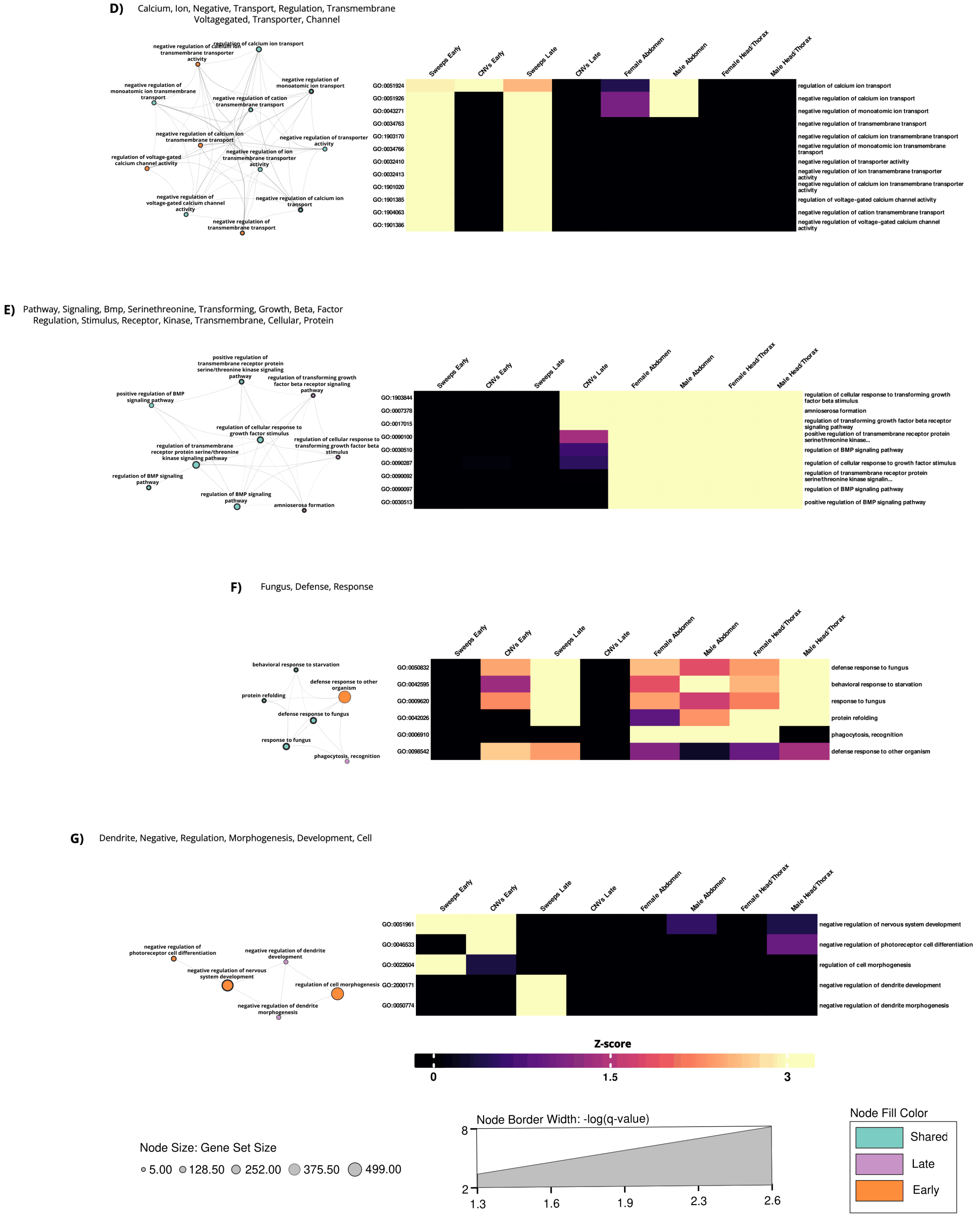
D)
Calcium, Ion, Negative, Transport, Regulation, Transmembrane Voltagegated, Transporter, Channel
regulation of calcium ion transport
negative regulation of calcium ion transmembrane transporter activity
negative regulation of monoatomic ion transmembrane transport
negative regulation of monoatomic ion transport
negative regulation of cation transmembrane transport
negative regulation of calcium ion transmembrane transport
negative regulation of transporter activity
regulation of voltage-gated calcium channel activity
negative regulation of ion transmembrane transporter activity
negative regulation of voltage-gated calcium channel activity
negative regulation of calcium ion transport
negative regulation of transmembrane transport
Sweeps Early
CNVs Early
Sweeps Late
CNVs Late
Female Abdomen
Male Abdomen
Female Head/Thorax
Male Head/Thorax
GO:0051924 regulation of calcium ion transport
GO:0051926 negative regulation of calcium ion transport
GO:0043271 negative regulation of monoatomic ion transport
GO:0034763 negative regulation of transmembrane transport
GO:1903170 negative regulation of calcium ion transmembrane transport
GO:0034766 negative regulation of monoatomic ion transmembrane transport
GO:0032410 negative regulation of transporter activity
GO:0032413 negative regulation of ion transmembrane transporter activity
GO:1901020 negative regulation of calcium ion transmembrane transporter activity
GO:1901385 regulation of voltage-gated calcium channel activity
GO:1904063 negative regulation of cation transmembrane transport
GO:1901386 negative regulation of voltage-gated calcium channel activity
E)
Pathway, Signaling, Bmp, Serinethreonine, Transforming, Growth, Beta, Factor Regulation, Stimulus, Receptor, Kinase, Transmembrane, Cellular, Protein
positive regulation of transmembrane receptor protein serine/threonine kinase signaling pathway
regulation of transforming growth factor beta receptor signaling pathway
positive regulation of BMP signaling pathway
regulation of cellular response to growth factor stimulus
regulation of cellular response to transforming growth factor beta stimulus
regulation of transmembrane receptor protein serine/threonine kinase signaling pathway
regulation of BMP signaling pathway
regulation of BMP signaling pathway
amnioserosa formation
Sweeps Early
CNVs Early
Sweeps Late
CNVs Late
Female Abdomen
Male Abdomen
Female Head/Thorax
Male Head/Thorax
GO:1903844 regulation of cellular response to transforming growth factor beta stimulus
GO:0007378 amnioserosa formation
GO:0017015 regulation of transforming growth factor beta receptor signaling pathway
GO:0090100 positive regulation of transmembrane receptor protein serine/threonine kinase...
GO:0030510 regulation of BMP signaling pathway
GO:0090287 regulation of cellular response to growth factor stimulus
GO:0090092 regulation of transmembrane receptor protein serine/threonine kinase signalin...
GO:0090097 regulation of BMP signaling pathway
GO:0030513 positive regulation of BMP signaling pathway
F)
Fungus, Defense, Response
behavioral response to starvation
defense response to other organism
protein refolding
defense response to fungus
response to fungus
phagocytosis, recognition
Sweeps Early
CNVs Early
Sweeps Late
CNVs Late
Female Abdomen
Male Abdomen
Female Head/Thorax
Male Head/Thorax
GO:0050832 defense response to fungus
GO:0042595 behavioral response to starvation
GO:0009620 response to fungus
GO:0042026 protein refolding
GO:0006910 phagocytosis, recognition
GO:0098542 defense response to other organism
G)
Dendrite, Negative, Regulation, Morphogenesis, Development, Cell
negative regulation of photoreceptor cell differentiation
negative regulation of dendrite development
negative regulation of nervous system development
regulation of cell morphogenesis
negative regulation of dendrite morphogenesis
Sweeps Early
CNVs Early
Sweeps Late
CNVs Late
Female Abdomen
Male Abdomen
Female Head/Thorax
Male Head/Thorax
GO:0051961 negative regulation of nervous system development
GO:0046533 negative regulation of photoreceptor cell differentiation
GO:0022604 regulation of cell morphogenesis
GO:2000171 negative regulation of dendrite development
GO:0050774 negative regulation of dendrite morphogenesis
Z-score
0
1.5
3
Node Size: Gene Set Size
5.00
128.50
252.00
375.50
499.00
Node Border Width: -log(q-value)
8
2
1.3
1.6
1.9
2.3
2.6
Node Fill Color
Shared
Late
Early

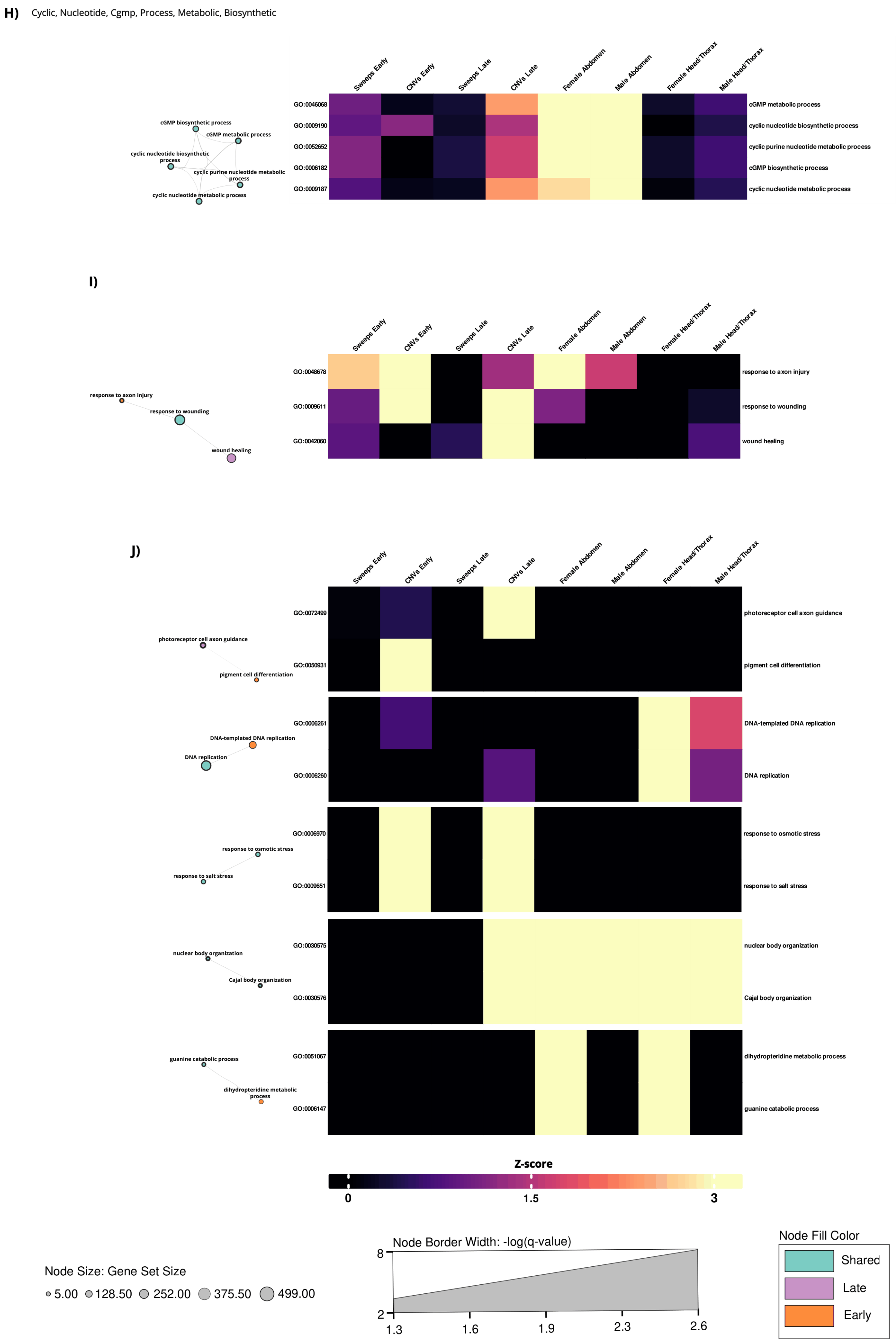

**Fig. S37.** Functional modules (A-J) identified in our multi-evidence enrichment analysis, where evidence for enrichment was present in both E and L lines. The left most four columns in the heat-map represents sweep and CNV calls, the right most four columns differential gene expression (color denotes strength of evidence).

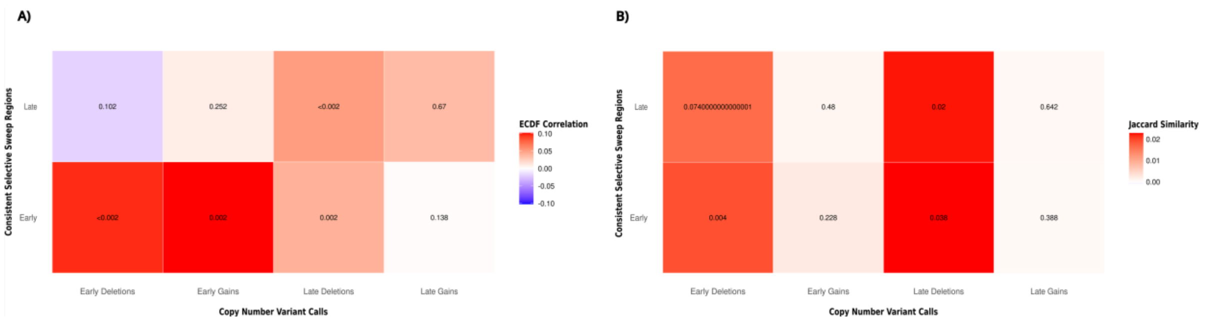

**Fig. S38**. Genomic association between CNV calls and selective sweep intervals. Heatmaps of (A) ECDF-based relative-distance correlation and (B) Jaccard similarity between CNV call categories and selective sweep call intervals (those identified in ≥3 populations per regime). Colour scales are capped to the maximum absolute value of each metric. Numbers indicate raw *P*-values for each pairwise comparison. Significant correlations of interval distributions are observed for selective sweep regions in the E regime and both deletions and gains in the E regime. Deletions in L lines also show significant association with both E and L sweep calls. We note that, despite reaching statistical significance, ECDF correlations are low and the absolute Jaccard overlap between CNV and sweep intervals was generally very low.

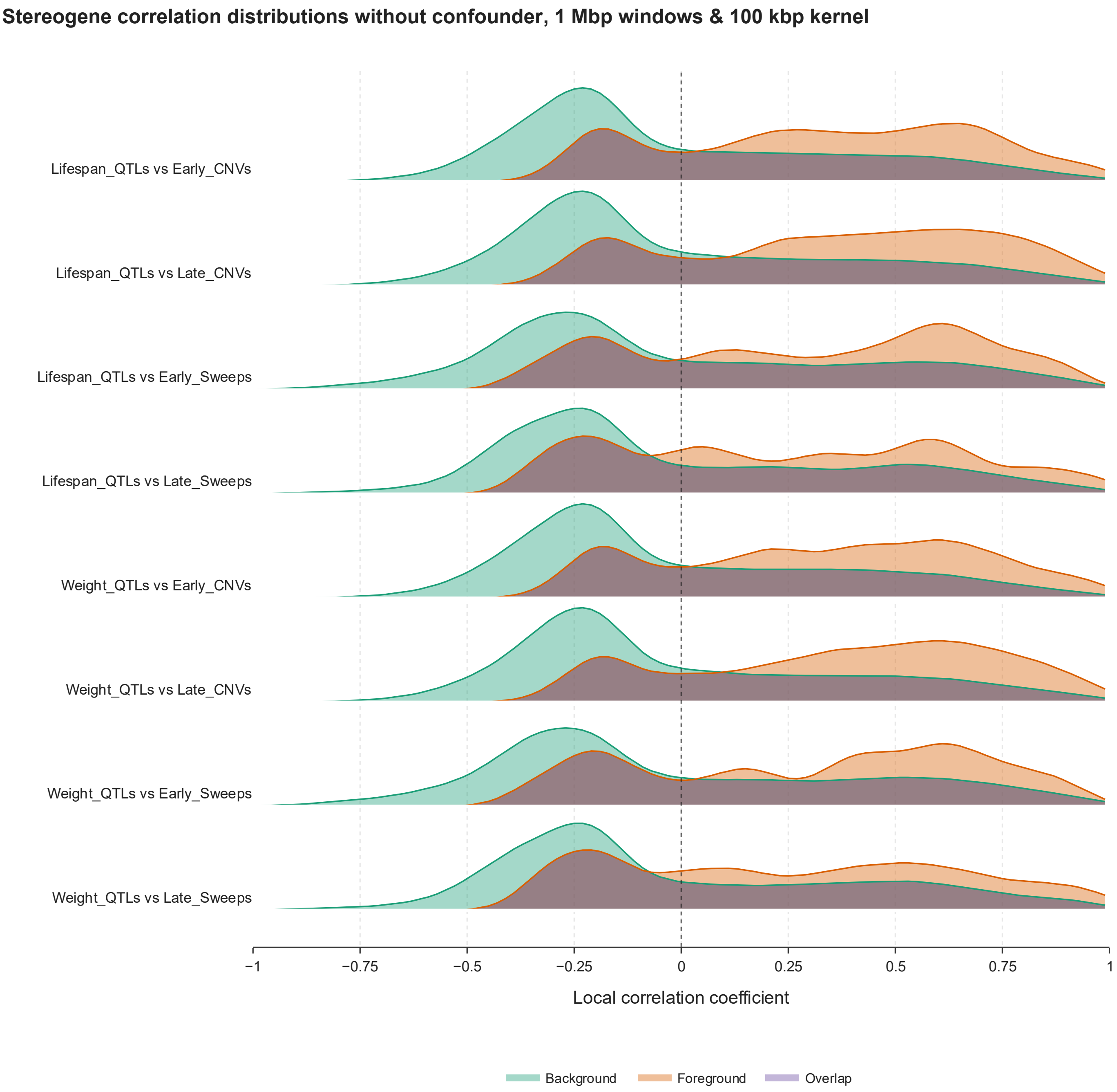

**Fig. S39**. The genome-wide spatial correlations between LOD scores and sweep/CNV calls were sizeable. This figure shows the observed distribution of correlations across the genome (window size 1 Mbp, Kernel width 100 kbp) estimated using using StereoGene (Stavrovskaya et al. 2017) (orange), as well as the random expectation based on spatial permutation (green), for the spatial association between LOD scores for life span and weight versus CNV and sweep calls in L and E lines.

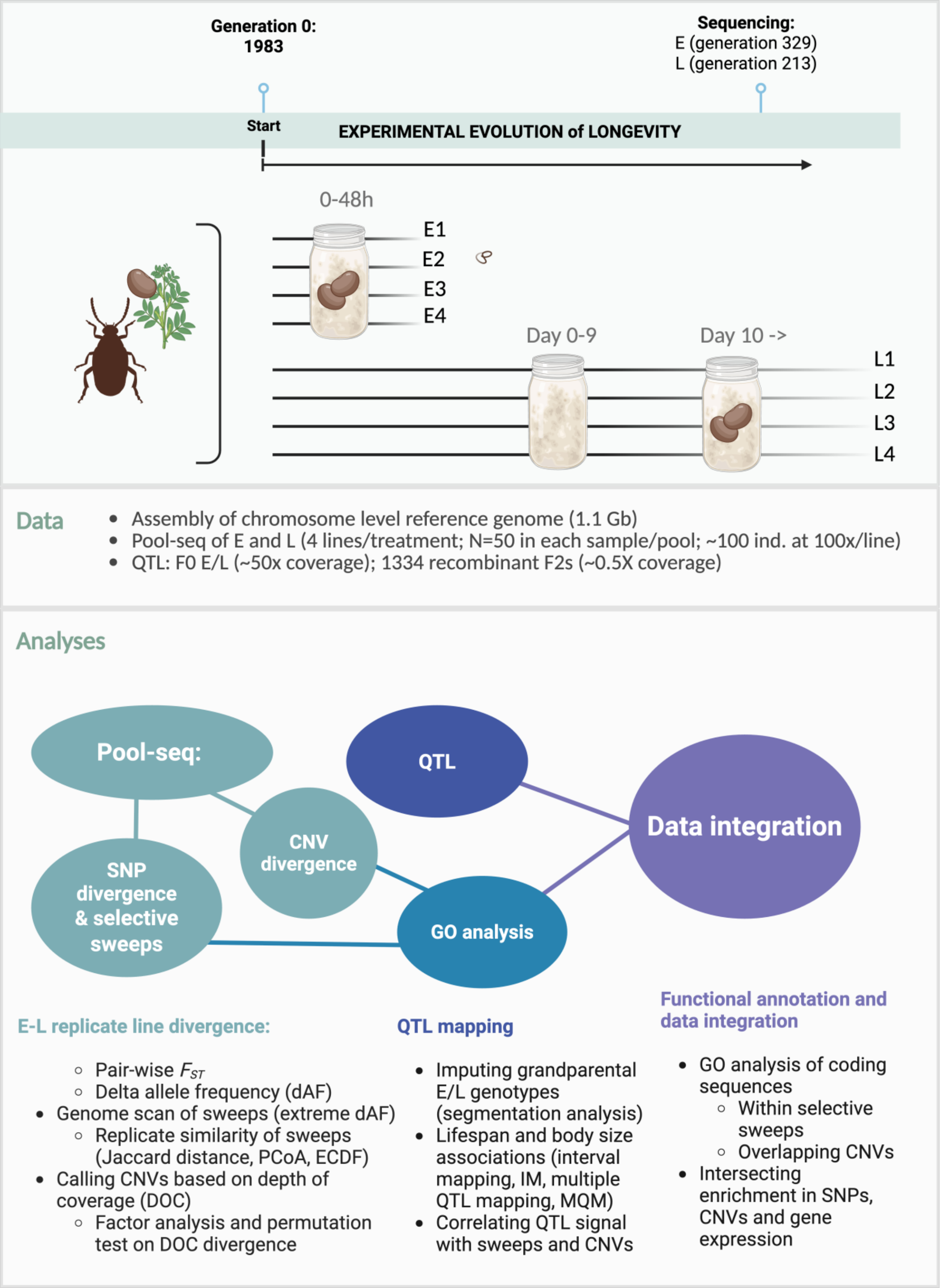

**Fig. S40**. An overview of the experimental procedures, data utilized and analytical steps employed during the study.
